# Sex without crossing over in the yeast *Saccharomycodes ludwigii*

**DOI:** 10.1101/2021.04.22.440946

**Authors:** Ioannis A. Papaioannou, Fabien Dutreux, France A. Peltier, Hiromi Maekawa, Nicolas Delhomme, Amit Bardhan, Anne Friedrich, Joseph Schacherer, Michael Knop

**Author notes:** Faculty of Agriculture, Kyushu University, Fukuoka, Japan (Hiromi Maekawa).

## Abstract

Meiotic recombination is a ubiquitous function of sexual reproduction throughout eukaryotes. Here we report that recombination is extremely suppressed during meiosis in the yeast species *Saccharomycodes ludwigii*. DNA double-strand break formation, processing and repair are required for normal meiosis but do not lead to crossing over. Although the species has retained an intact meiotic gene repertoire, genetic and population analyses suggest the exceptionally rare occurrence of meiotic crossovers. We propose that *Sd. ludwigii* has followed a unique evolutionary trajectory that possibly derives fitness benefits from the combination of frequent fertilization within the meiotic tetrad with the absence of meiotic recombination.

## Main

Sex constitutes the prevailing reproductive mode throughout the eukaryotic tree of life^1^. At its core lies a periodic ploidy cycling, accomplished through meiosis, during which haploid gametes are produced, and mating, which ensures the restoration of the original ploidy level. Meiosis is thought to have arisen early in the eukaryotic evolution and is a ubiquitous attribute of sexual life cycles^2^. During the first meiotic division (meiosis I), parental chromosomes are recombined and separated, while in the second sister chromatids segregate (meiosis II). Random reassortment and recombination of homologous chromosomes in meiosis I lead to novel genetic constellations in the offspring. These are used as substrates for natural selection, for promotion of advantageous and purging of deleterious genetic combinations^3, 4^. However, the significant complexity and biological costs of sex render its widespread occurrence paradoxical, and the questions of its evolutionary origin, persistence and functions have been outstanding enigmas in biology^5–7^.

*Saccharomyces cerevisiae* (baker’s yeast) is a premier model organism for the study of cellular, molecular and evolutionary biology, and it has been used to gain major insights into the molecular aspects of meiosis^8, 9^. Following pairing of homologous chromosomes, meiotic recombination is initiated in prophase I with the formation of DNA double-strand breaks (DSBs) by the topoisomerase-like protein Spo11^10^. Several pathways mediate the subsequent repair of these DSBs and pathway choice is regulated by a multitude of meiosis-specific factors that act in concert with the DNA repair machinery. One possibility involves the use of the homologous chromosome as repair template, which can ultimately lead to the generation of chimeric DNA molecules. This process may involve either reciprocal exchange of the chromosomal regions that flank the DSB site (crossover, CO) or gene conversion without reciprocal exchange (non-crossover, NCO)^8, 9^. Crossing over leads to the establishment of chiasmata (Fig. 1d), which ensure the physical interconnection of bivalents that is important for the faithful segregation of homologous chromosomes in meiosis I^11^. In many, but not all, organisms this is facilitated by the synaptonemal complex, which mediates stable synapsis^9^.

**Fig. 1:**
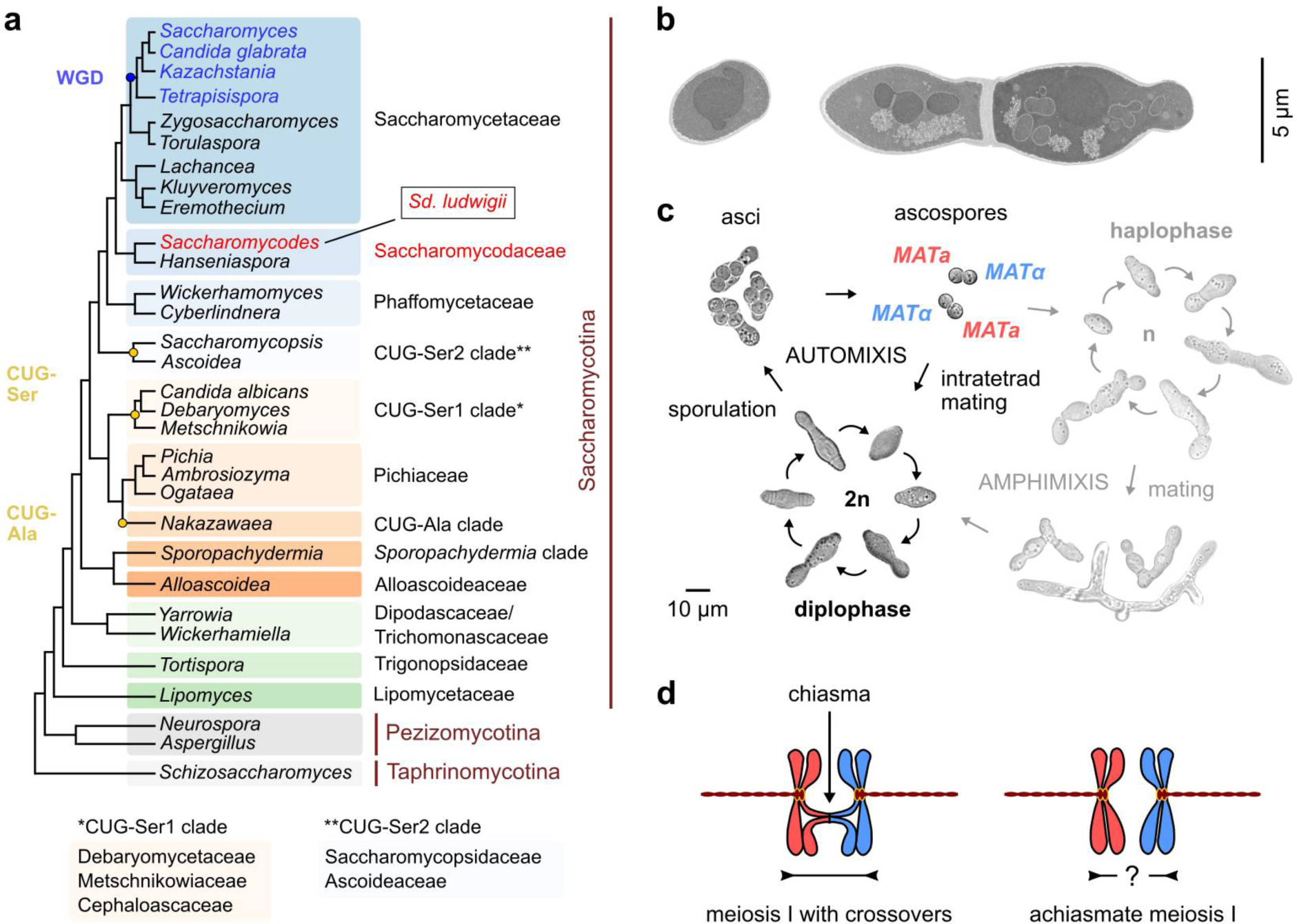
The budding yeast *Saccharomycodes ludwigii*. **a**, Classification of *Sd. ludwigii* in the yeast phylogeny. WGD: whole-genome duplication; CUG-Ser1/2 and CUG-Ala: clades deviating from the universal genetic code. **b**, Transmission electron microscopy images of representative *Sd. ludwigii* vegetative cells (reference strain NBRC 1722). **c**, Life cycle of *Sd. ludwigii*. Brightfield images of strains NBRC 1722 and its diploid progenitor NBRC 1721 are shown. **d**, Chiasmata in meiotic prophase I contribute to spindle stabilization and faithful segregation of homologous chromosomes. Meiosis in *Sd. ludwigii*, however, has been suggested to be achiasmate^38, 39^.

The extent of genetic exchange in meiosis and thus the generation of genomic diversity are strongly influenced by the frequency and distribution of COs along chromosomes. Homeostatic regulation controls these parameters in many organisms^12, 13^, with different outcomes in different species. These range from one obligatory CO per chromosome in *Caenorhabditis elegans*^14^ to 10 or more in *Schizosaccharomyces pombe*^15^. Among yeasts, the frequency of meiotic COs per chromosome ranges from an average of 2.5 COs in *Lachancea kluyveri* to its highest level in *Sc. pombe*^15–19^. Furthermore, generation of diversity is influenced by the mating behavior (breeding), which can involve gametes of variable genetic relatedness. Mating occurs between unrelated gametes in outbreeding, which ensures high genetic diversity in the offspring. On the other hand, inbreeding (self-fertilization) refers to mating between genetically related gametes that originate from the same individual or clonal line. Inbreeding is generally considered to lead to lower heterozygosity, which can compromise the adaptive potential of populations^20^.

Mating of gametes from the same meiotic event represents a particular type of inbreeding referred to as intratetrad mating or automixis^21^. This appears to be the most frequent breeding strategy in *S. cerevisiae*^22–24^. If automixis brings together chromosomes that had been separated during meiosis I (non-sister components), it is described as “central fusion” or “first division restitution”. This has important genetic consequences, since it maintains parental heterozygosity around the centromeres, reducing the risk of deleterious alleles being exposed due to homozygotization^25–27^. The degree to which parental heterozygosity is restituted upon intratetrad mating correlates inversely with the frequency of COs during meiosis. In the extreme case of absent crossing over parental genomes would be fully reconstituted and their heterozygosity maintained. Evolutionary models suggested that high frequency of deleterious mutations could promote automixis in the absence of meiotic recombination^28^.

Chromosome segregation in meiosis I without any recombination (achiasmate meiosis) has been observed in one of the two sexes^6, 29^ or in individual chromosomes^30, 31^ of a few organisms. Furthermore, unusually low overall recombination rates have been reported in some species^32, 33^. However, these data should be interpreted with caution since sampling bias, insufficient marker coverage and incomplete genome assemblies may have hindered the identification of COs in many of these cases. In addition, most of these studies could not exclude the possibility of crossing over near chromosome ends, which has been observed in a number of species across kingdoms^32^.

The budding yeast species *Saccharomycodes ludwigii* (Fig. 1a-b) preferentially undergoes intratetrad mating (Fig. 1c), ensured by strong inter-spore bridges that efficiently keep spores together in pairs of opposite mating types during germination and subsequent mating^34, 35^. This organism was used in the early days of yeast genetics for the pioneering description of heterothallism by Øjvind Winge at the Carlsberg laboratory^35, 36^. During his studies, Winge also observed in this species an unusual segregation pattern of two cell morphology markers^35^. The subsequent interpretation by Lindegren was that the two genes were not linked and that “each is so close to the spindle attachment that segregation invariably occurs at the Meiosis I without crossing over”^37^. While it could have been a coincidence that both markers were linked to centromeres, later work by the Oshima lab surprisingly revealed that the same behavior was displayed by any combination of more than 20 genetic markers tested. These findings led to the conclusion that this “may be due to the absence of crossing over in *Sd. ludwigii*”^38, 39^.

Here we continued this work by exploring the extent and types of meiotic interactions between homologous chromosomes in *Sd. ludwigii*. For this, we performed whole-genome sequencing and high-contiguity *de novo* assembly of wild-type strains, which enabled a high-resolution DNA variant segregation analysis of meiosis. We combined bioinformatic analyses of the meiotic gene repertoire with a functional study of key meiotic components in order to derive a better understanding of the meiotic mechanisms in this species. We also searched for signs of recombination on an evolutionary scale between divergent strains of *Sd. ludwigii* in comparison to other species with different levels of meiotic recombination. In order to unravel the relative contributions of mutational pressure and recombination to genome evolution, we determined the genome-wide mutation rate and bias using a mutation accumulation experiment. Our results provide insights into the unique sexual lifestyle of the yeast species *Sd. ludwigii*, and they propose this organism as a particularly suitable model system for the study of the evolution of recombination rates and their impact on genome evolution.

## Results

### The *Sd. ludwigii* genome

To investigate meiotic recombination in *Sd. ludwigii*, we first used long-read PacBio sequencing to generate a high-contiguity *de novo* genome assembly of our reference haploid strain NBRC 1722 (Supplementary Table 1). The 12.5-Mb assembly consists of 7 chromosomal scaffolds, in concordance with PFGE karyotyping^39, 40^ (Fig. 2a) and previous genetic mapping^38, 39^. Gene synteny and DNA motif analyses (Extended Data Fig. 1a) enabled the identification of putative point centromeres on all chromosomes. The extremities of all scaffolds share subtelomere-related repetitive sequences and genes, while telomeric DNA repeats were also detected in 9 cases (Extended Data Fig. 1b). A single mating-type locus (*MATalpha*) was identified in the centromeric vicinity of chrE (Fig. 2a), in congruence with the known heterothallic nature of the species^35, 36^.

**Fig. 2:**
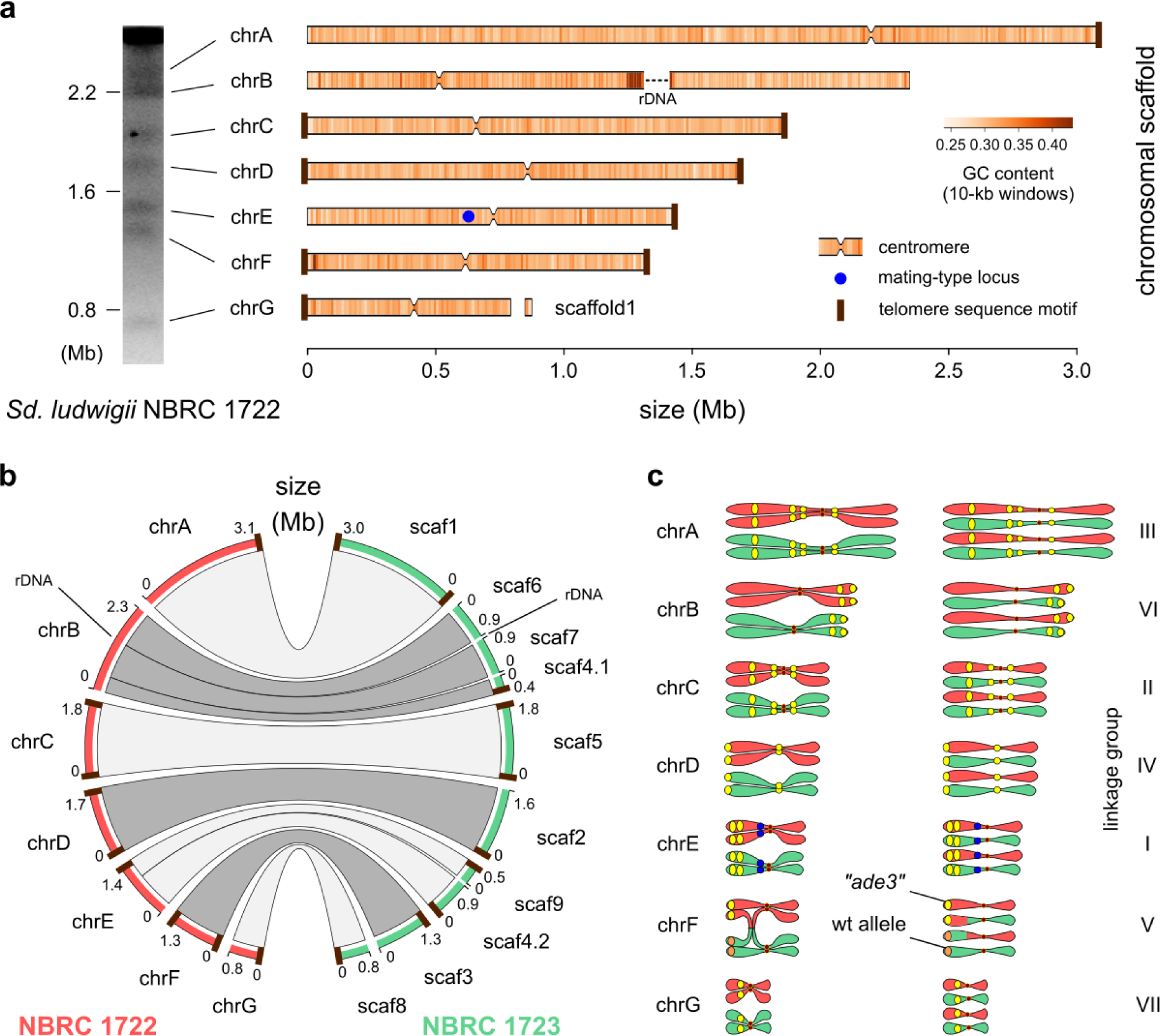
Genome structure of *Sd. ludwigii* and mapping of previously used genetic markers. **a**, The nuclear genome of the haploid reference strain NBRC 1722 is organized in 7 chromosomes, resolved as chromosomal bands on a PFGE gel (left) and depicted as bars (right). The only gap in the assembly (in chrB) corresponds to the internal part of the rDNA region, which is flanked by assembled rDNA repeats. **b**, Genomic comparison of the *Sd. ludwigii* parental strains used in the previous genetic analyses. **c**, The 7 previously defined *Sd. ludwigii* linkage groups39 were matched to the 7 chromosomes of the reference genome assembly, and 16 of the previously used markers were unambiguously mapped on the assembly (yellow circles). The *MAT* locus is shown as a blue circle (in chrE). One of the rare COs that were previously detected using tetrad analysis was tracked down to a reciprocal exchange event between the marker “*ade3*” (corresponding to the *ADE4* gene) and the centromere of chrF, depicted here as an example.

The *Sd. ludwigii* genome shares many features with Saccharomycetaceae members that diverged before the whole-genome duplication event (Fig. 1a), including genome size, number of chromosomes, point centromeres, overall gene synteny, number of genes (5,031 ORFs) and frequency of introns (3.3%). Remarkably, *Sd. ludwigii* chromosomes exhibit GC levels that are unusually low for a yeast species (30.9% on average; Fig. 2a), as shown by a comparison to 100 yeast and other fungal genomes (Extended Data Fig. 2a; Supplementary Table 2). The comparative analysis further revealed exceptionally high coverage in *Sd. ludwigii* by AT-rich low-complexity regions and simple sequence repeats (microsatellites), as well as enrichment in transposable elements (Extended Data Fig. 2b-e).

Meiotic crossing over could be hindered in chromosomal regions of *Sd. ludwigii* that do not align during meiotic prophase I due to major sequence variation or structural differences of the homologous chromosomes. To investigate whether high genomic dissimilarity could be responsible for the scarcity of COs observed in previous analyses^38, 39^ that used our reference strain as one of the parents, we also sequenced and assembled *de novo* the genome of the second parent used in those experiments (NBRC 1723; Supplementary Table 1). The two parental genomes are highly collinear (Fig. 2b) and similar at the DNA sequence level (99.6% identity on average), apart from the terminal region of chrA and the longest part of chrF, which exhibit lower degrees of sequence identity (95.3% on average). These findings exclude the possibility that the absence of COs could be due to major differences between homologous chromosomes.

Another possibility for the explanation of the extreme rarity of previously detected COs could be a non-random chromosomal distribution of the UV-generated genetic markers used in those analyses^38, 39^. By combining the results of our gene prediction and annotation of the reference assembly with the phenotypic gene annotations of *S. cerevisiae* (www.yeastgenome.org), we identified the chromosomal positions of 16 out of the 24 used markers (Fig. 2c). This analysis revealed that all chromosomes were covered by markers, most of which were quite distant from the corresponding centromeres. Therefore, the previously used experimental setup^38, 39^ appears sufficient for capturing the majority of COs in the largest part of the genome.

### The *Sd. ludwigii* meiotic gene machinery

To gain insight into the genetic causes of the unusual meiotic behavior of *Sd. ludwigii*, we manually refined the gene prediction and we curated the annotation of *Sd. ludwigii* genes based on the available *S. cerevisiae* dataset (www.yeastgenome.org). Gene prediction yielded a total of 5,347 genes, of which 5,031 are protein-coding ORFs. Among these genes, we identified homologs of 272 out of the 284 genes of *S. cerevisiae* with annotated functions in meiosis (Gene Ontology terms; www.yeastgenome.org), using protein sequence similarity and gene synteny as criteria (Fig. 3a; Supplementary Table 3). All genes that are required for wild-type levels of meiotic recombination in *S. cerevisiae* have homologs in *Sd. ludwigii*, with the only exception of *MER1*^41^ (Fig. 3a). This gene codes for a meiosis-specific splicing activator for introns in the genes *AMA1*, *HFM1*, *REC107* and *SPO22*, which function in chromosome pairing, meiotic recombination and cell cycle regulation. Deletion of *MER1* in *S. cerevisiae* abrogates the expression of these 4 target genes and causes major defects in meiotic recombination and progression^41, 42^. However, the Mer1 regulon of *Sd. ludwigii* appears largely rescued, as no introns were detected in 3 of its target genes (*AMA1*, *HFM1* and *REC107*), while *SPO22* has only a minor role in *S. cerevisiae* meiosis^43^. By investigating the phylogenetic distribution of *MER1* we confirmed its ancestral origin as syntenic homologs were identified in the distant families Phaffomycetaceae, Ascoideaceae and Saccharomycopsidaceae (Fig. 1a; 3a). Therefore, its absence from *Sd. ludwigii* and the closely related *Hanseniaspora* species must be due to a secondary loss early in the evolution of their family (i.e. Saccharomycodaceae; Fig. 3a).

**Fig. 3:**
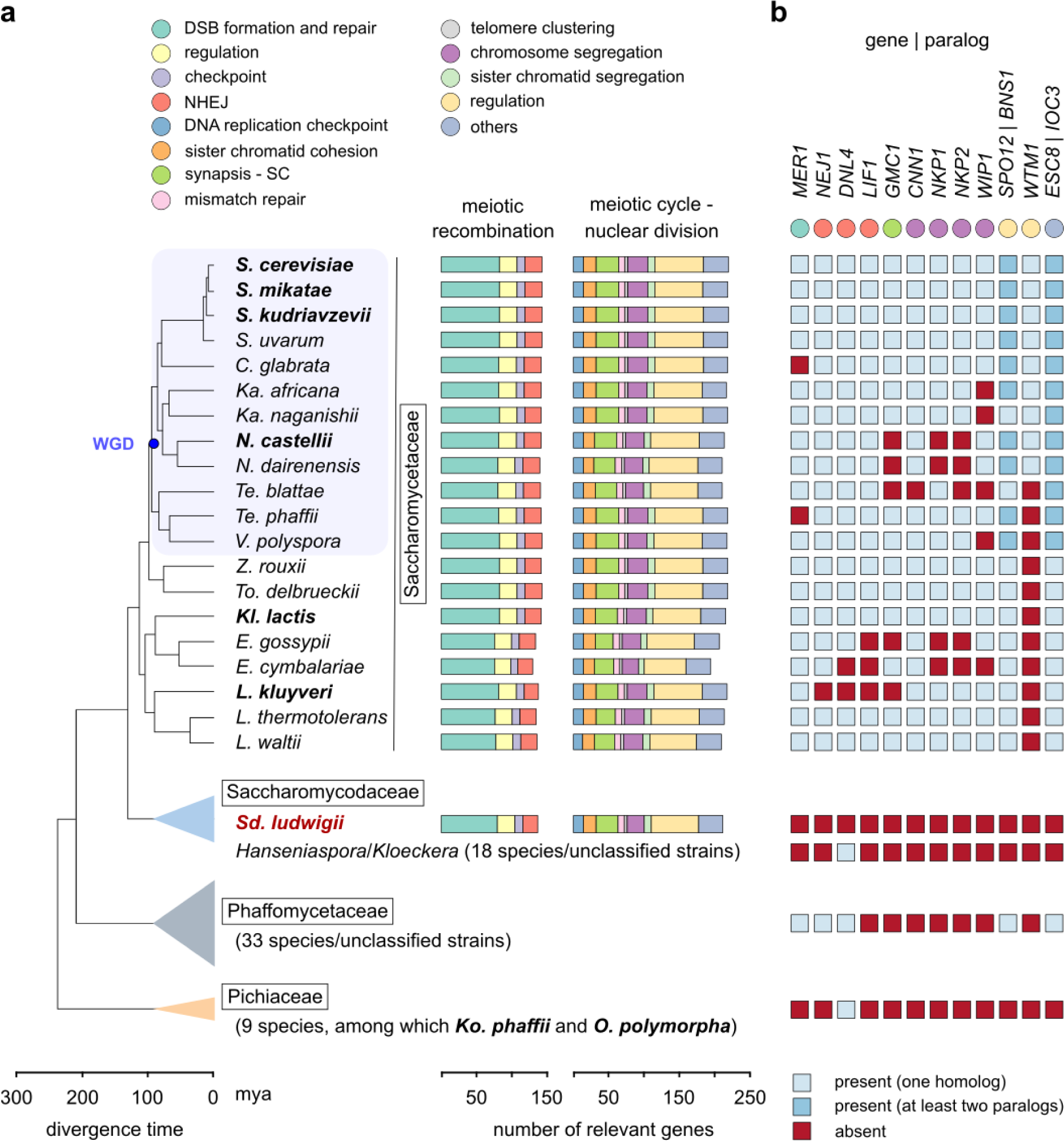
*Sd. ludwigii* possesses a nearly full meiotic gene complement. **a**, Meiotic genes of Sd. ludwigii and other yeast representatives. Genes were classified into functional groups (top) based on the associated GO terms of their *S. cerevisiae* homologs (www.yeastgenome.org), and the total gene count for each of them is plotted for each species (right). Apart from the presumably asexual species *C. glabrata*^49^, Hanseniaspora jakobsenii, for which sexual sporulation has not been reported to date^50^, and the mitosporic yeasts Candida mycetangii, *C. orba, C. stellimalicola, C. ponderosae, C. montana* and *C. vartiovaarae*, all other 71 species included in this analysis are sexual. The names of species with documented meiotic recombination appear in bold^17,19,44,51,52^. **b**, Meiosis/meiotic recombination-related genes that are absent from the reference *Sd. ludwigii* strain are summarized here. Presence or absence of homologs is also indicated for all other species, for comparison. The function of each gene is indicated by a colored circle beneath its name (color code as in **a**).

Apart from *MER1*, no homologs were detected in *Sd. ludwigii* for 11 additional genes with meiotic functions, namely *NEJ1*, *DNL4*, *LIF1*, *GMC1*, *CNN1*, *NKP1*, *NKP2*, *WIP1*, *SPO12/BNS1* (paralogs in *S. cerevisiae*), *WTM1* and *ESC8/IOC3* (paralogs in *S. cerevisiae*) (Fig. 3b). In contrast to *MER1*, inactivation of any of these genes in *S. cerevisiae* does not abrogate meiotic crossing over. With the exception of *DNL4*, these genes are also absent from members of the Pichiaceae that are known to form COs in meiosis, such as *Komagataella phaffii* and *Ogataea polymorpha*^19, 44^. A phylogenetic analysis suggested that 7 of these genes (i.e. *LIF1*, *NKP1*, *NKP2*, *CNN1*, *WIP1*, *GMC1* and *WTM1*) are present only in Saccharomycetaceae members, and thus have probably emerged after the separation of the *Sd. ludwigii* lineage. Among the remaining 4 genes, *DNL4* and *NEJ1* (involved in non-homologous end joining, NHEJ^45^) are also missing from *Lachancea kluyveri*, which is capable of meiotic crossing over^17^, while *SPO12*^46^ and *ESC8*^47^ have only minor and indirect roles in *S. cerevisiae* meiosis. Overall, our results demonstrate very limited meiotic gene loss and the retention in *Sd. ludwigii* of the essential gene machinery for meiotic recombination. Eight of the putative *Sd. ludwigii* meiotic proteins have little or no similarity to their *S. cerevisiae* homologs, and the identification of their genes was mostly based on synteny. These are *MEI4*, *NDJ1*, *PSY3*, *SPO16*, *REC104*, *POL4*, *IML3* and *HED1* (Supplementary Table 3). Using similarity searches with either the *S. cerevisiae* or *Sd. ludwigii* protein sequences as queries, most of these genes could not be detected in the closely related *Hanseniaspora* species, which is consistent with extensive loss of genes involved in DNA repair, cell cycle regulation and meiosis from those species^48^.

### Meiotic recombination components are required for *Sd. ludwigii* meiosis

The presence of a nearly intact meiotic gene machinery suggests that these genes are functional in *Sd. ludwigii*. This is further supported by our finding that deletion of *SPO11*, which in other organisms initiates meiotic recombination by generating DNA DSBs, led to significantly reduced sporulation and spore viability (Fig. 4a), similarly to *spo11* hypomorphs of *S. cerevisiae*^53, 54^. These results indicate an important function of this protein in meiosis. Since *S. cerevisiae* Δ*spo11* mutants also exhibit defects in chromosome synapsis^10^, we also deleted *SAE2*, an endonuclease that operates downstream of Spo11 in DSB processing and cleavage of DNA-Spo11 intermediates^55^. This led to a similar sporulation defect to that of the Δ*spo11* strain (Extended Data Fig. 3), suggesting a role for the endonuclease function of Spo11 in *Sd. ludwigii* meiosis. In order to confirm directly the occurrence of Spo11-dependent DSBs, we investigated the meiotic localization of Rad51, a recombinase that is involved in DSB repair^8^. Using an anti-Rad51 antibody, we detected discrete foci in spreads of meiotic *Sd. ludwigii* chromosomes. No such foci were observed in Δ*rad51* or Δ*spo11* mutants, indicating the specificity of detection and the dependence of DSBs on Spo11 (Fig. 4b). We detected 5-12 foci per nucleus, which would be consistent with the formation of 1-2 Spo11-dependent DSBs per meiotic chromosome. Our results overall suggest that Spo11 is required for normal meiosis in *Sd. ludwigii*, through the generation of meiotic DNA DSBs.

**Fig. 4:**
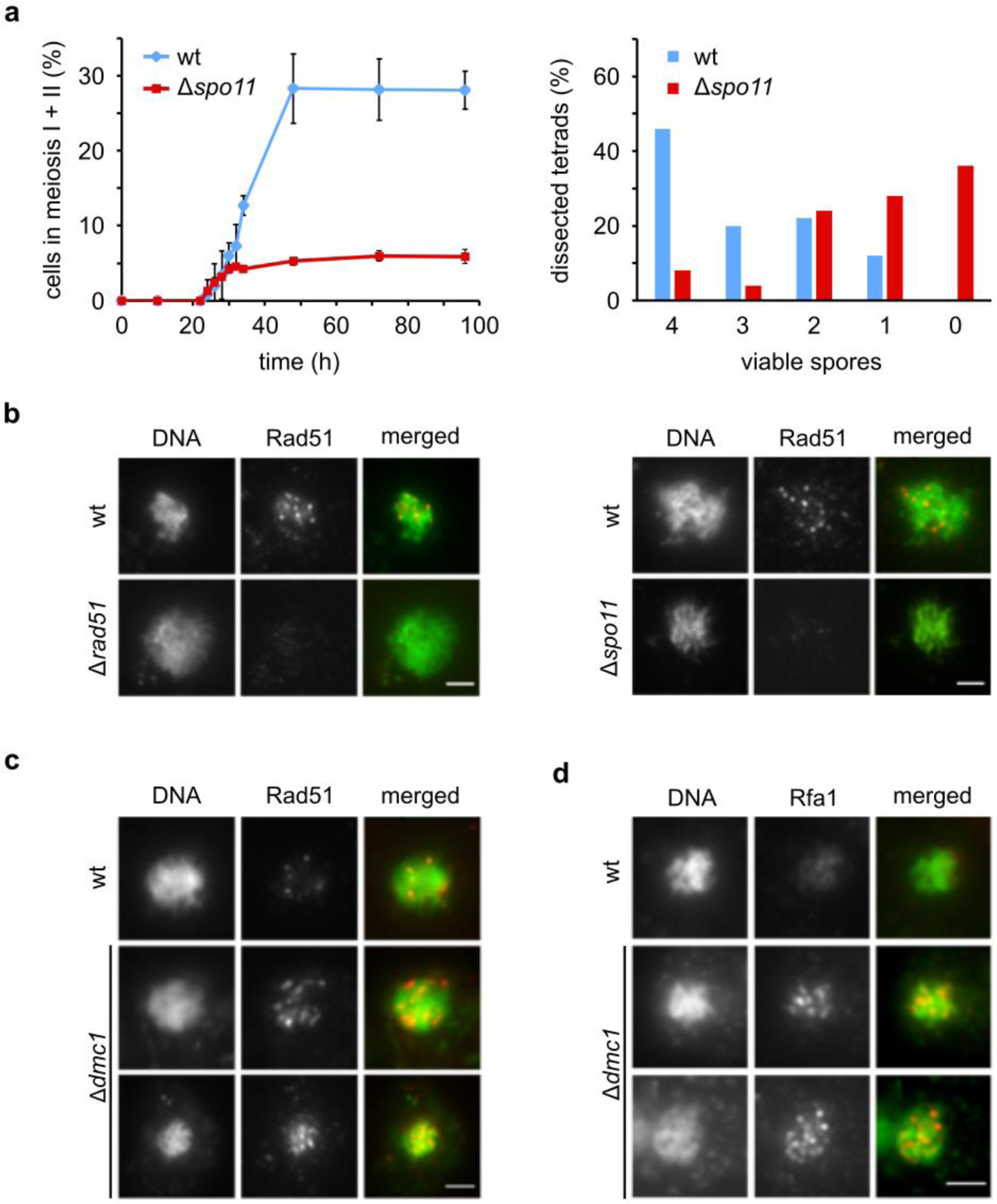
Core meiotic recombination components are required for normal meiosis in *Sd. ludwigii*. **a**, Meiotic time-course analysis of a homozygous *Δspo11 Sd. ludwigii* mutant in comparison to its wild type (left). After induction of meiosis, samples were withdrawn at the indicated time points and their cellular DNA content was stained with Hoechst 33258 to determine the fractions of binucleate (meiosis I) and tetranucleate (meiosis II) cells. Error bars: SD (3 replicates for each strain). Spore viability was examined using tetrad dissection and viable colony counting (right). A total of 50 tetrads of each strain were analyzed. b-d, Meiotic nuclear spreads of sporulating cells with the indicated genotypes (homozygous diploids). Cellular DNA was stained with Hoechst 33258. Immunostaining was performed using anti-Rad51 or anti-Rfa1 antibodies. Bars: 1 µm.

We also performed Rad51 immunostaining in meiotic spreads of *Sd. ludwigii* cells deleted for *DMC1*, a ubiquitous meiosis-specific recombinase involved in DSB repair as well as in homolog interactions. In this case, we observed elongated, filamentous foci (Fig. 4c), which is consistent with the anticipated function of Dmc1 as an inhibitor of the Rad51 strand exchange activity^56^. Another conserved single-stranded (ss) DNA-binding protein that is involved in DNA replication, repair and recombination, is Rfa1. We detected expanded Rfa1 foci in Δ*dmc1* meiotic spreads, whereas no foci were detected in wild-type spreads (Fig. 4d). This is congruent with the normally transient nature of ssDNA-binding activity of Rfa1, which is replaced by Dmc1, and the accumulation of ssDNA at DSBs in Δ*dmc1* cells, similarly to *S. cerevisiae*^57^. Overall, formation of meiotic DSBs and their recombinational repair, mediated by key meiotic components, are required for normal meiosis in *Sd. ludwigii*, consistently with what is known from *S. cerevisiae*.

### High-resolution analysis of meiotic segregation in *Sd. ludwigii*

Our findings suggest that meiotic recombination might still be occurring in *Sd. ludwigii*, albeit with patterns that perhaps prevented detection by segregation analyses using only few markers. To address this, we performed a high-resolution genome-wide SNP segregation analysis of *Sd. ludwigii* meiosis (Fig. 5a; Extended Data Fig. 4a). We used as parents the reference strain and a spore that we isolated from a tetrad of a geographically distant strain (spore 122; Supplementary Table 1). Crosses between these two haploid strains frequently formed tetrads with 4 viable spores. Long-read sequencing and *de novo* assembly of the second parent’s genome did not reveal significant karyotypic differences that could prevent pairing of homologous chromosomes in this cross (Fig. 5b). Variant calling between the two strains resulted in a total of 199,392 high-quality SNPs, with an average density of 34 SNPs per kb of genomic sequence on chrC, chrF, chrG and parts of chrA and chrB, whereas the remaining parts of the genome exhibited lower heterozygosity levels, of approx. 3 SNPs per kb on average (Extended Data Fig. 4b).

**Fig. 5:**
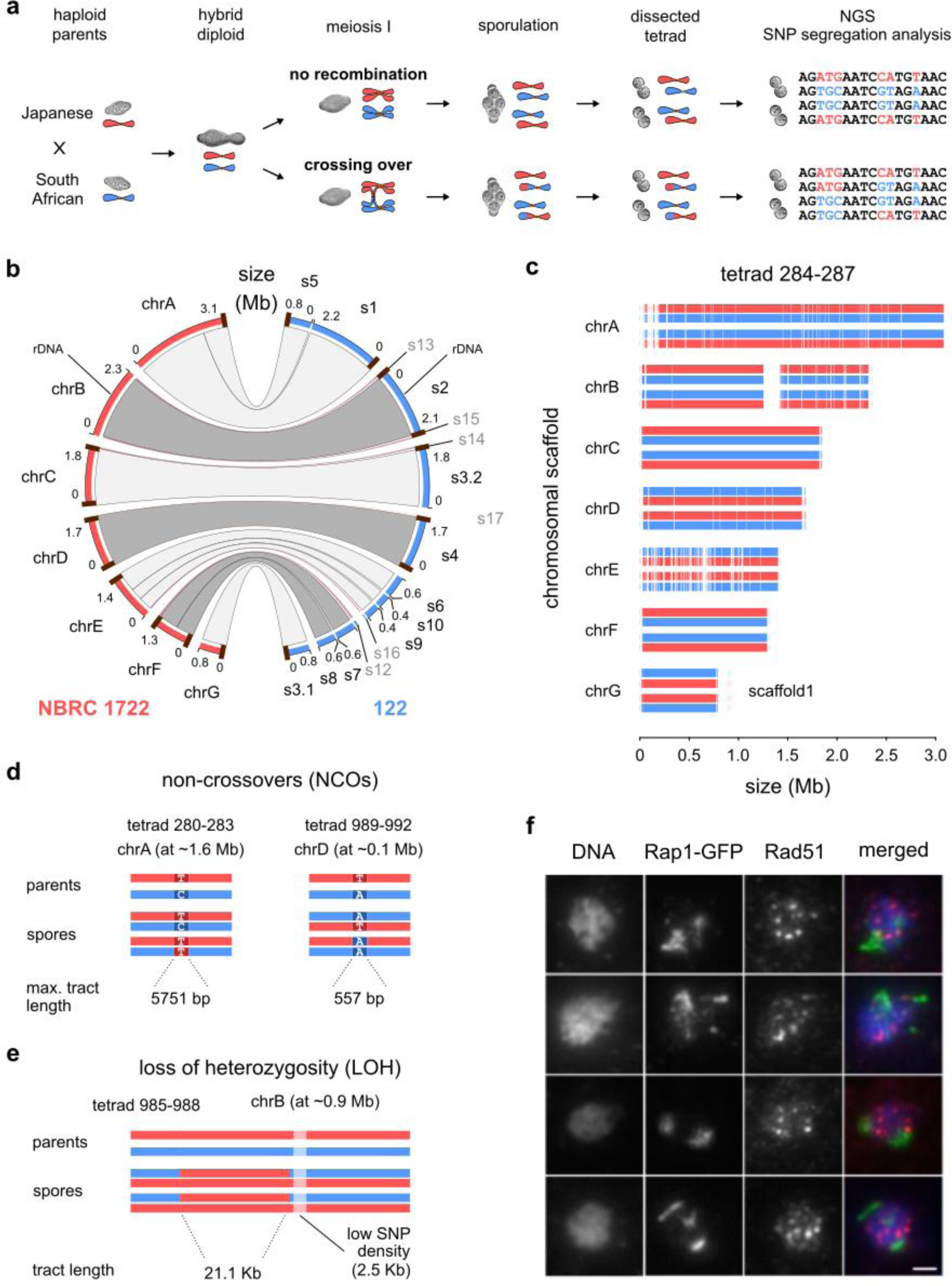
A high-resolution segregation analysis revealed the absence of crossing over in *Sd. ludwigii* meiosis. **a**, Experimental setup (further details in Extended Data Fig. 4a). **b**, Genomic comparison of the haploid parental strains. **c**, Results of the genome-wide SNP segregation analysis for a representative tetrad. Each dimorphic position of the genome was assigned to one of the parental haplotypes (red for the reference strain, blue for spore 122). **d**, Two verified single-marker NCOs were detected by the SNP segregation analysis. **e**, A single LOH event was detected in tetrad 985-988, in close proximity to another shorter region with unusually low SNP density. **f**, Immunostaining of meiotic nuclear spreads of a *Sd. ludwigii* strain expressing Rap1-GFP using anti-Rad51 and anti-GFP antibodies. Bar: 1 μm.

Sequencing of all spores from 5 full tetrads from this cross followed by analysis of the SNP segregation patterns revealed complete absence of meiotic COs (Fig. 5c). The same was observed when we extended our analysis to 2 tetrads from a cross between 2 spores of the second parents’ lineage (spores 100 and 102; Supplementary Table 1; Extended Data Fig. 4a). These results confirm the previous genetic evidence that suggested extreme rarity of meiotic crossing over in *Sd. ludwigii*^38, 39^. Among the 7 tetrads, we detected 2 independent single-marker NCOs (gene conversion tracts with 3:1 segregation patterns; Fig. 5d), which were validated by PCR amplification and sequencing. Their maximum tract lengths were 5.8 and 0.6 kb, respectively (calculated based on the distance between their closest flanking SNPs with 2:2 segregation patterns). This suggests that at least some of the meiotic DSBs are processed by interactions that use the homologous chromosome as repair template without, however, leading to CO formation. Finally, our analysis revealed a 21.1 kb-long loss-of-heterozygosity (LOH) tract (Fig. 5e), which could have resulted from a mitotic gene conversion event during diploid growth preceding sporulation. This LOH event occurred in close proximity to a 2.5 kb-long region of distinctly lower SNP density than that of the flanking chromosomal regions (Fig. 5e), indicating that another ancient event had occurred in the same region.

Telomeric regions of chromosomes are highly repetitive (Extended Data Fig. 1b) and this could compromise the sensitivity of our method for the detection of COs in these regions^16, 17, 19, 32^. To investigate the possibility of crossing over in telomeric or telomere-proximal regions, we used meiotic chromosome spreads to compare the localization of GFP-tagged Rap1, a cytological marker of telomeres^58^, and that of Rad51 foci as an indicator of meiotic DSBs^8^. Our experiments consistently demonstrated absence of co-localization of the 2 proteins (Fig. 5f). Therefore, initiation of recombination is not biased towards telomeres or adjacent regions in *Sd. ludwigii*, and the extreme suppression of crossing over appears to affect the entire genome.

### Search for signs of historical crossing over in *Sd. ludwigii*

The unusual suppression of homolog interactions in *Sd. ludwigii* meiosis motivated us to search for signs of recombination by comparing divergent strains of international origin. For this, we sequenced 10 available haploid and diploid strains (Supplementary Table 4), two of which were found to be aneuploids (2n+1 trisomies) by a read coverage analysis (Extended Data Fig. 5a). Variant calling and comparison to our reference strain revealed variable sequence divergence (∼18,000-308,000 SNPs) and heterozygosity levels in diploids (∼1,300-60,000 heterozygous SNPs) (Supplementary Table 4). Apart from the very divergent haploid strain PC99_R_1, sequence variation showed non-uniform patterns of distribution across the genome, being mostly restricted to particular chromosomes (spore 122) or, in most strains, to chromosomal segments (Extended Data Fig. 5b).

Phylogenetic analysis of all strains based on their genome-wide SNP content revealed the presence of a major cluster of 7 strains, and 4 more divergent ones (Fig. 6a). When we compared the dendrograms of individual chromosomes, we observed chromosome-dependent topologies and distances for particular strains (Fig. 6a-b; Extended Data Fig. 5c). Our analyses revealed that chromosomes of strains with unusually unstable topologies differ significantly in SNP density from their genomic average (Extended Data Fig. 5e). One possible explanation for these contrasting chromosome-specific signals could be that absence or rarity of meiotic recombination over an extended period of time has deprived chromosomes of the corresponding homogenizing effect, allowing them to accumulate SNPs independently from their homologs to a certain extent. We gained further support for this hypothesis by performing the same analysis in the yeast *Lachancea kluyveri*, a Saccharomycetaceae member with a similar genome size and chromosome number to *Sd. ludwigii*, but with demonstrated capability of meiotic recombination^17^. The analysis revealed very stable topologies in *L. kluyveri*, with all chromosomes yielding the same topology as the whole genome (Extended Data Fig. 5d, 5f; Fig. 6b).

**Fig. 6:**
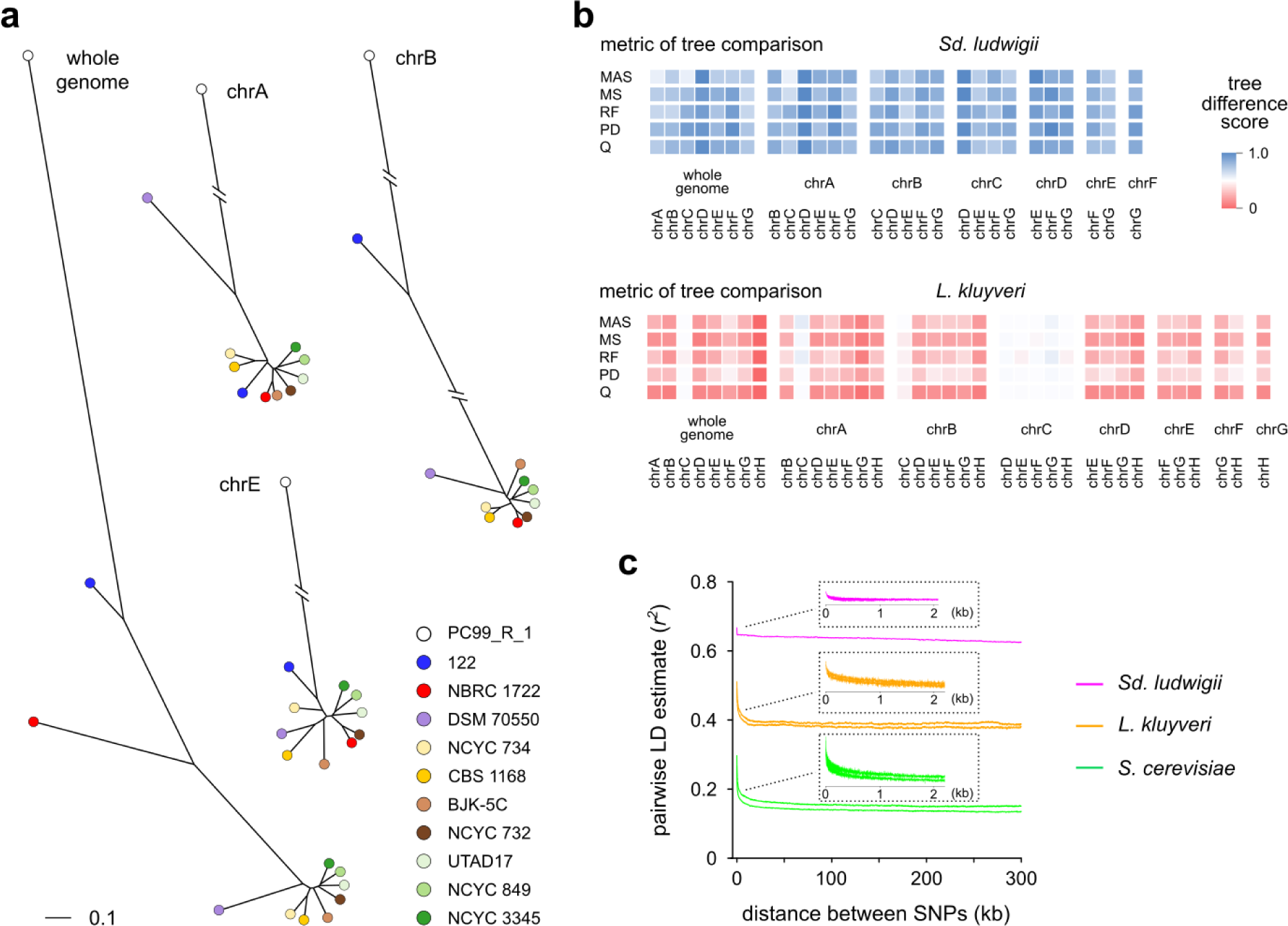
Crossing over has been absent or unusually rare in *Sd. ludwigii* over evolutionary time. **a**, Dendrograms of *Sd. ludwigii* strains, constructed by neighbor-joining analysis of all SNPs genome-wide (“whole genome”) or SNPs of individual chromosomes. The dendrograms of the remaining chromosomes are shown in Extended Data Fig. 5c. **b**, Scores of tree difference (5 topological, unrooted metrics) between all combinations of whole-genome SNPs and SNPs of individual chromosomes, for *Sd. ludwigii* and *L. kluyveri*. The analyzed *L. kluyveri* strains and their corresponding dendrograms are provided in Extended Data Fig. 5d. The results shown here are based on normalized distances to the average values of 1,000 randomly generated tree pairs (uniform average method). MAS: unrooted maximum agreement subtree distance; MS: matching split distance; RF: Robinson-Foulds distance; PD: path difference distance; Q: Quartet distance. **c**, Decay of LD as a function of physical distance between SNP marker pairs, for *Sd. ludwigii* in comparison to representative *L. kluyveri* and *S. cerevisiae* groups of strains. The moving averages of the LD estimate *r*^2^ values are plotted for all SNP marker pairs (genome-wide) of different physical distances. All data (no line smoothing) for the first 2 kb of distance are plotted in the insets.

Next, we performed a genome-wide linkage disequilibrium (LD) analysis in *Sd. ludwigii*, in comparison to *L. kluyveri* and *S. cerevisiae*, which are characterized by low and high meiotic recombination activity, respectively^16, 17^. Generation of LD decay curves, for which the LD estimate *r*^2^ for each pair of SNPs is plotted versus their physical distance, revealed striking differences between the species (Fig. 6c). The overall levels of LD were unusually high in *Sd. ludwigii*, intermediate in *L. kluyveri*, and lowest in *S. cerevisiae*. Furthermore, LD decayed extremely fast in *Sd. ludwigii*, where it reached its plateau within the first ∼50 bp, in contrast to the more gradual decay of the 2 other species, which is typical of recombining organisms^59, 60^. Four of the *Sd. ludwigii* chromosomes exhibited even higher LD levels and faster decay than the genomic average (Extended Data Fig. 6a). To address the limitation of the small sample size in these analyses (due to the limited availability of wild-type *Sd. ludwigii* strains), we always compared groups of the same size (11 strains for all species), and we also compared multiple groups of randomly selected strains of *L. kluyveri* and *S. cerevisiae*. Curves of LD decay were quite stable and the observed differences between species were independent of the particular group considered (Extended Data Fig. 6b). These results are consistent with the hypothesis of absent or very rare meiotic recombination in *Sd. ludwigii* over evolutionary time, which would decrease the overall LD levels and decelerate its decay.

### Mutation rate and genomic evolution in *Sd. ludwigii*

A hypothetically elevated mutation rate could be a compensatory solution for the generation of genetic diversity in *Sd. ludwigii* in the absence of meiotic recombination, or even serve as a driver of achiasmate meiosis^28^. Furthermore, we reasoned that a significant AT bias of spontaneous mutations could explain the particularly low GC content of the *Sd. ludwigii* genome (Extended Data Fig. 2a). We investigated these hypotheses directly by a mutation accumulation analysis in 3 founder *Sd. ludwigii* strains: our haploid reference strain, a corresponding homozygous diploid strain, and the hybrid diploid strain used in our SNP segregation experiment (Table 1; Supplementary Table 1). We adopted a single-cell population bottlenecking regime of growth on agar plates to evolve 60 independent mutation accumulation lines for ∼2,000 generations each. Such an experimental setup ensures fixation of the majority of spontaneous mutations and minimizes elimination of non-lethal deleterious mutations due to selection^61^. Sequencing and genome-wide analysis revealed a total of 186 line-exclusive single nucleotide mutations (SNMs) (Supplementary Table 5). This corresponds to an overall base-substitutional mutation rate (*μ*_bs_) of 5.70 × 10^-11^ - 1.20 × 10^-10^ mutations per base per generation, for the 3 strain genealogies (Table 1). These rates, which are in line with the rate of another recently analyzed *Sd. ludwigii* diploid strain^62^ (7.3 × 10^-11^), fall within the range of unicellular eukaryotes, and they are even slightly lower than those of other yeast species^62, 63^. Therefore, mutation rate in *Sd. ludwigii* is not unusually elevated. Similarly, the overall transition-to-transversion (Ts:Tv) ratio (1.21 in *Sd. ludwigii*) is comparable to the values reported for *S. cerevisiae* and expected for organisms with no cytosine methylation^63^.

**Table 1:** Mutation accumulation analysis. The base-substitutional mutation rate (*μ*_bs_), its Poisson 95% confidence interval (CI), the AT bias (weighted by genomic nucleotide composition), and the transition:transversion ratio (Ts:Tv) of spontaneous single-nucleotide mutations are provided for each strain genealogy.

| Strain | # generations (average) | # mutations (# lines) | $\mu_{bs}$ (/site/generation) | Poisson 95% CI | AT bias | mutation-driven equilibrium GC content | observed GC content | Ts:Tv ratio |
| --- | --- | --- | --- | --- | --- | --- | --- | --- |
| haploid (NBRC 1722) | 2,037 | 85 (30) | $1.20 \times 10^{-10}$ | $9.62 \times 10^{-11}$ - $1.49 \times 10^{-10}$ | 2.92 | 0.255 | 0.296 | 1.18 |
| isogenic diploid (YLFP17-4) | 1,920 | 48 (19) | $5.70 \times 10^{-11}$ | $4.20 \times 10^{-11}$ - $7.55 \times 10^{-11}$ | 5.85 | 0.146 | 0.296 | 1.53 |
| hybrid diploid (YLFP188-1) | 2,016 | 53 (11) | $9.55 \times 10^{-11}$ | $7.16 \times 10^{-11}$ - $1.25 \times 10^{-10}$ | 2.63 | 0.276 | 0.296 | 0.96 |

The *Sd. ludwigii* genome has a remarkably low GC content. We determined that, similarly to the majority of eukaryotes, *Sd. ludwigii* has a strong AT bias of spontaneous mutations, with SNMs that convert GC to AT occurring on average 3.6 times more frequently than SNMs in the opposite direction (Table 1). Based on this, the theoretical equilibrium GC content of the *Sd. ludwigii* genome should be 23.3%, if it were determined by spontaneous mutations alone. In these calculations we excluded repeat regions (∼7% of the genome), since they are difficult to analyze and are associated with a high indel formation frequency^64^. The observed genomic GC content of the analyzed genomic fraction was 29.6%, which cannot be explained by the effect of spontaneous mutations only. Therefore, balancing forces in the opposite direction, such as selection on GC or GC-biased gene conversion^63^, are probably also present in *Sd. ludwigii*.

### Mitotic LOH in *Sd. ludwigii*

The striking rarity of homolog interactions in *Sd. ludwigii* meiosis prompted us to investigate the levels of mitotic homologous recombination, to understand whether the observed strong suppression is specific to meiosis. Further analysis of the 11 mutation accumulation lines of the hybrid diploid strain (Table 1) revealed a total of 22 LOH events (Supplementary Table 6), half of which occurred on chrA. Chromosome A is the largest chromosome (representing ∼25% of the *Sd. ludwigii* genome). Whether this relative enrichment of events on this chromosome explains its lower SNP density compared to other chromosomes (Extended Data Fig. 4b) is unclear. Similarly non-uniform distributions of LOH events have been observed in *S. cerevisiae*^65, 66^. Among the detected events, we identified one long terminal event (in chrC; ∼183 kb), which could be attributed to either a mitotic crossover or a break-induced replication (BIR) event^67^. The remaining events were shorter interstitial LOH regions with a maximum length of 12.5 kb (Supplementary Table 6). These may be associated with mitotic crossing over or gene conversion^65, 66, 68^. The total genomic rate of LOH events was 9.9 × 10^-4^ events per generation, which is moderately lower than the 3-10 times higher frequencies reported for *S. cerevisiae*^65, 66^. Therefore, the core machinery for repair of spontaneous DSBs during the vegetative life phase of *Sd. ludwigii* is present and functional. The extreme suppression of recombination is, therefore, limited to meiosis.

## Discussion

Intermixing of genomes for the generation of variability is the unifying theme of sexual reproduction. Although sexual life cycles appear very diverse in nature^6^, genomic reshuffling through meiotic reassortment and recombination of homologous chromosomes is regarded as a common denominator of sexual cycles^2, 7^. The level of heterozygosity present in individuals and, therefore, the extent of genetic diversity in populations are largely dependent on mating strategies. This is clearly illustrated in the case of yeasts that can alternate between outcrossing and inbreeding reproductive regimes^23^. Frequent intratetrad mating, in particular, preserves high levels of heterozygosity in parts of the genome (Fig. 7a), ensures efficient purging of deleterious mutations, and provides fitness advantages^25, 27, 28, 69^. Suppression of recombination in organisms that often engage in intratetrad mating could be beneficial for their evolution by extending the preservation of heterozygosity to larger parts of their genomes^26^ (Fig. 7b). Here, we describe the yeast *Saccharomycodes ludwigii* as an example of an organism with extremely suppressed meiotic recombination in a sexual cycle that is predominated by intratetrad mating. Our study shows that this species suppresses meiotic crossing over throughout its genome, and this behavior has persisted over evolutionary time. Our findings corroborate the hypothesis of Yamazaki and coworkers, who found that tetratype tetrads, putatively indicative of crossing over, are extremely rare in *Sd. ludwigii* crosses (i.e. 22 tetratypes out of a total of 2,234 analyzed tetrads)^38, 39^.

**Fig. 7:**
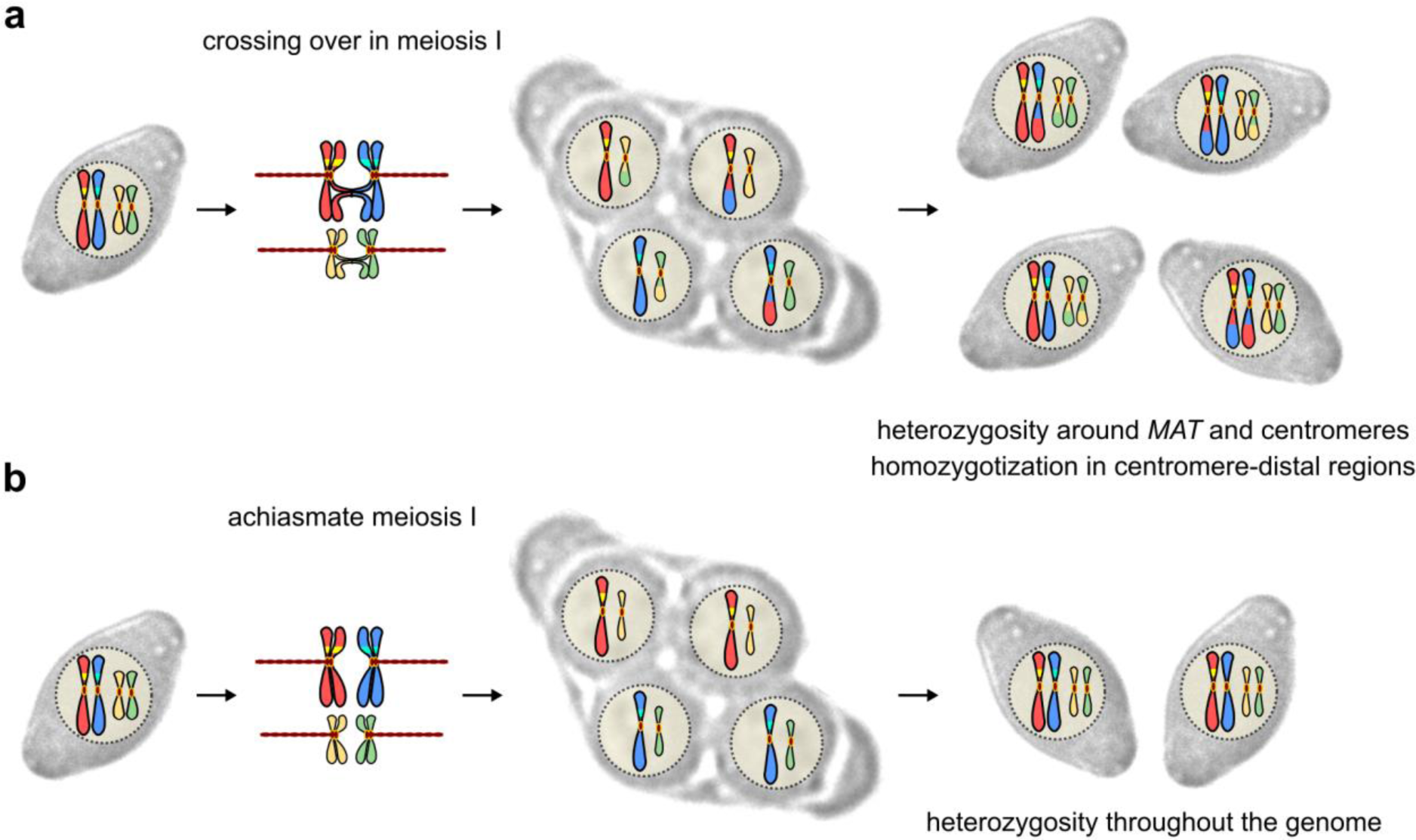
The evolutionary coupling of high rates of non-sister intratetrad mating with extremely suppressed meiotic recombination in *Sd. ludwigii* maximizes preservation of heterozygosity. **a**, Non-sister intratetrad mating with crossing over in meiosis I. The mating-type locus is linked to the centromere of its chromosome (which is often observed in species that engage in intratetrad mating, e.g. *S. cerevisiae* and *Sd. ludwigii*). **b**, Non-sister intratetrad mating with achiasmate meiosis I, which normally happens in *Sd. ludwigii*.

Meiotic recombination is initiated during the prophase of meiosis I of sexual eukaryotes with the programmed introduction of DNA DSBs by the conserved transesterase Spo11^8, 10, 52^. It is thought that meiotic cells induce those dangerous lesions that put their genomic integrity at risk to induce the formation of COs that promote accurate segregation of homologous chromosomes by the establishment of chiasmata and exchange genetic information between them^8, 9, 11^. Given the high toxicity of DNA DSBs, organisms have evolved multiple pathways for their repair^70^. Meiotic cells preferentially rely on homologous recombination, which is the only pathway that can repair a subset of these DSBs to form COs, whereas the remainder are repaired as NCOs or using the sister chromatid as template for repair^70, 71^. The pathway choice decision is controlled by multiple factors, including those that determine DSB positioning and the formation and processing of recombination intermediates^72^, and it can be subject to evolutionary selection. Our experiments revealed that Spo11-mediated introduction of DSBs is required for normal meiosis in *Sd. ludwigii*. Given that immunostaining of Rad51 for visualizing DSBs may underestimate their number by half or more^73^, we conclude that the observed 5-12 Rad51 foci could correspond to at least 1-2 DSBs per chromosome in *Sd. ludwigii* meiosis. The demonstrated involvement of Dmc1 in DSB processing suggests that at least a fraction of these DSBs are processed through homolog-templated repair pathways^8, 9, 11^, but with a strong bias towards NCOs as suggested by the absence of COs. This is supported by the detection of 2 NCOs in our meiotic SNP segregation analysis, which *per se* provides evidence for the presence of interhomolog interactions in this species. Since such events typically involve short gene conversion tracts (usually 1-2 kb in *S. cerevisiae*^16, 74^), more short-tract NCOs might have occurred in our analyzed tetrads but evaded detection due to the lower SNP density in some genomic regions, masking of low-complexity fractions of the genome (∼7%) in our analysis, or restoration of the original genotype by the mismatch repair system. On the other hand, our study conclusively demonstrated absence of crossing over throughout the genome, which confirms its previous detection by tetratype analyses at extremely low levels^38, 39^. These data suggest that meiotic interactions between homologs are needed for meiosis I in *Sd. ludwigii*, but DSB repair is biased towards NCOs, without reciprocal exchange of the flanking regions. Finally, considerable involvement of NHEJ in the repair of meiotic DSBs is unlikely, given that *Sd. ludwigii* misses 4 components of the major NHEJ pathway (i.e. Nej1, Dnl4, Lif1 and Pol4)^75^, which is anyway suppressed during meiosis of other organisms^76, 77^.

In the absence of crossing over in *Sd. ludwigii* meiosis I, compensatory mechanisms must be present to prevent nondisjunction of homologous chromosomes due to the lack of chiasmata^9^. In other organisms with sex- or chromosome-specific suppression of crossing over, components of the synaptonemal complex (SC) or specific cohesins facilitate segregation of meiotic chromosomes^78, 79^. The unique meiotic behavior of *Sd. ludwigii* renders the functional dissection of its meiotic mechanisms an interesting goal of future research. In this context, the investigation of ZMM proteins that provide links between the SC and repair of meiotic DSBs^80, 81^, as well as the characterization of components that regulate homolog bias and pathway choice^9, 72, 82^, could provide useful insights. Preliminary analyses in our lab revealed the presence of a SC and considerable roles of its core components in meiosis.

In order to understand the particular regulation of meiotic recombination in *Sd. ludwigii*, an interesting subject might be the fast evolving meiotic proteins, i.e. the ones that have retained little homology to the corresponding *S. cerevisiae* genes: Rec104 participates in a complex with Spo11 during the initiation of recombination^83^; Mei4 also interacts with DSB formation proteins^84, 85^; and Ndj1 and Spo16 are involved in the regulation of CO formation and distribution^86, 87^. In the cases of Mei4, Rec104 and Pol4, the characteristic protein domains were not identifiable in the *Sd. ludwigii* putative homologs and the *Sd. ludwigii* Pol4 (component of NHEJ^75^) appears truncated (41.2% of the *S. cerevisiae* Pol4 length) and is predicted to be intrinsically disordered, suggesting loss of function and pseudogenization. However, further functional investigations depend on the development of *Sd. ludwigii* strains with higher sporulation rates and better synchronization of meiosis I than the currently available ones.

Our findings support the hypothesis that achiasmate meiosis might have evolved in the *Sd. ludwigii* lineage on the basis of mutual selection between suppression of meiotic recombination and frequent intratetrad mating. The fixation of such a sexual lifestyle suggests that suppression of crossing over must have provided advantages to the organism, which must also have developed sufficient mechanisms or behaviors to compensate for the absence of crossing over. Low recombination rates could have arisen through the accidental loss of an important meiotic gene, e.g. *MER1*^42^, which might have spread to fixation possibly due to genetic hitchhiking by suppressing recombination along its entire chromosome. The need to overcome the negative consequences of the loss of crossing over could have driven the selection of secondary mutations that rescued the ability of the species to complete meiosis without the detrimental effects of frequent chromosome nondisjunction. One possible strategy that is often observed in *Sd. ludwigii* strains^88^ is the completion of meiosis with only one division, which leads to two-spore asci with diploid spores exhibiting excellent viability, according to our observations. Secondary loss of *SPO12*, which is absent from *Sd. ludwigii*, might have helped to achieve this and prevent a meiotic arrest in cell cycle progression, since natural *S. cerevisiae* strains lacking *SPO12* and *SPO13* exhibit the same behavior^89^. Alternatively, the risk of inviable spores due to chromosomal mis-segregation might be alleviated by mating in the tetrad with non-sister meiotic components, which would restore a full diploid chromosome complement (Fig. 7b) even in the case of aneuploid individual spores, which are still likely capable of mating^28, 90^. The absence of recombination would not permit the exchange of mating types between non-sister chromatids in meiosis I and, therefore, the requirement for opposite mating types would lead to full reconstitution of the parental diploid genomic content upon mating. If intratetrad mating is prevented, e.g. by experimental dissection of tetrads, frequent spore aneuploidy as well as accumulation of haploid lethal mutations would be expected. This organism appears to be avoiding these risks by particularly enforcing intratetrad mating in its life cycle through the development of robust interspore bridges that keep non-sister spores tightly linked together in its asci^91^. These structures efficiently promote intratetrad mating^35, 37, 92^, and our investigations have revealed that their mechanical integrity depends on chitin synthase III, since deletion of *CHS3* renders them much more fragile, similarly to *L. kluyveri*^17^. In this scenario, the duet of suppressed recombination and frequent intratetrad mating could have persisted because of the efficient preservation of heterozygosity and the fitness advantages that it could offer, according to previous research^26–28^.

The establishment of a lifestyle that successfully copes with the lack of meiotic recombination does not exclude the possibility of rare outbreeding^91^ that would occasionally permit intermixing of chromosomes, as was observed in our study (Fig. 6a; Extended Data Fig. 6c). In addition, the species seems to not have completely abolished its ability to rarely recombine its genome^38, 39^, which is reflected in the preservation of the corresponding gene arsenal, and the partial rescuing of its Mer1 regulon, possibly to compensate for the loss of *MER1*. However, those processes would require the maintenance of chromosomal structural stability, without which the accumulation of gross rearrangements in homologs would further hinder synapsis in meiosis I and preclude recombination. The loss of the classical NHEJ pathway from the *Sd. ludwigii* lineage (Fig. 3) could contribute to the preservation of genomic collinearity between different lineages of this species over longer evolutionary periods.

Recombination, genomic GC content and mutation rate are thought to be linked, although the causality of their relationships is still not fully understood^63, 93^. We performed a mutation accumulation analysis to investigate the reason for the particularly low genomic GC level in *Sd. ludwigii*, which is reflected in numerous low-complexity regions and AT-rich microsatellites across its genome. Our study revealed that the base-substitution mutation rate is within the normal range of yeasts and other unicellular eukaryotes^63^. On the other hand, the strong detected AT bias of spontaneous mutations –similarly to many other organisms^63, 94^– in conjunction with the presumably low relevance of GC-biased gene conversion^61, 63^ due to the rarity of recombination, provides an explanation for the particularly low GC level of its genome. Nevertheless, balancing forces in the opposite direction must also act, given that the expected equilibrium genomic GC level would be 6.3% lower than the actual one if it were determined solely by the mutational load. Such forces could include selection on GC^95^, especially in the coding fraction of the genome due to codon usage constraints, and temporary or conditional changes of the mutation rate and/or bias^96^.

Loss of heterozygosity has been suggested to offer rapid evolutionary solutions for the increase of phenotypic diversity and the adaptation to changing environments, by exposing recessive beneficial alleles which are masked in the heterozygous diploid state, or by alleviating negative epistasis of alleles in the heterozygous state^66, 97, 98^. Such adaptive flexibility could be very important in the case of *Sd. ludwigii*, which is predicted to generally preserve heterozygosity throughout its genome due to intratetrad mating without recombination. Our results indicate that LOH events, often resulting from mitotic COs or NCOs, occur in *Sd. ludwigii* at frequencies comparable to the ones observed in *S. cerevisiae*^65, 66^, which are significantly higher than the base-substitution mutation rate. The suppression of recombination in *Sd. ludwigii* is, therefore, limited to meiosis, and LOH could be important for its adaptive potential.

Long-standing theories about sex suggest advantages of recombination for adaptation to changing environments by combining beneficial alleles and disengaging them from deleterious mutations. Our findings open up the possibility that the extreme suppression of crossing over in meiosis has been coupled with a high intratetrad mating rate in *Sd. ludwigii* to ensure adaptive genome evolution in a trajectory that does not rely on meiotic recombination. Therefore, we envision that *Sd. ludwigii* provides a new paradigm and an excellent natural forum, in which to study experimentally the relationship of genome evolution and the evolution of recombination rate by manipulating factors that could influence the frequency of recombination, such as the rate of outbreeding.

## Data availability

All sequencing datasets generated in this work have been deposited at the Sequence Read Archive (NCBI GenBank), included in the BioProjects with accession numbers PRJNA28063, PRJNA578491 and PRJNA639224. All genome assemblies have been deposited at GenBank as Whole Genome Shotgun projects, under the accession numbers JACTOA000000000 (version JACTOA010000000) for *Sd. ludwigii* strain NBRC 1722 (annotated genome assembly of our reference strain), JACTNW000000000 (version JACTNW010000000) for *Sd. ludwigii* strain NBRC 1723, and JACTNV000000000 (version JACTNV010000000) for *Sd. ludwigii* spore 122.

## Acknowledgements

We would like to acknowledge T. Yamazaki (University of Yamanashi, Japan) for his valuable feedback and useful advice on handling *Sd. ludwigii*. We are also grateful to A. Shinohara (Osaka University, Japan) for the anti-Rad51 antibody, E. Mancera (National Polytechnic Institute of Mexico) and L. Steinmetz (EMBL, Heidelberg, Germany) for the anti-Rfa1 antibody, J. Trček (University of Maribor, Slovenia) for the *Sd. ludwigii* isolate BJK_5C, and A. Giacomini (University of Padua, Italy) for the *Sd. ludwigii* isolate PC99/R. Funding/further support: This work was supported by a postdoctoral fellowship grant from the Cluster of Excellence *CellNetworks* (University of Heidelberg) to I.A.P. We also acknowledge support by the state of Baden-Württemberg through bwHPC for high-performance computing and SDS@hd for data storage (grant INST 35/1314-1 FUGG). Additional help was provided by the Flow Cytometry and the Deep Sequencing Core Facilities of the University of Heidelberg (both supported by *CellNetworks*).

## Author contributions

Conceptualization: M.K.; Methodology: M.K., I.A.P., J.S. and A.F.; Investigation: I.A.P., F.A.P. and H.M.; Formal analysis: I.A.P., F.D., A.F. and N.D.; Supervision: M.K. and I.A.P.; Visualization: I.A.P., F.D. and A.F.; Writing – Original Draft: I.A.P.; Writing – Review & Editing: I.A.P., M.K., J.S. and A.F.; Funding acquisition: M.K. and I.A.P.

## Competing interests

The authors declare no competing interests.

## Methods

### Yeast and bacterial strains – strain construction

All yeast strains used in this study are listed in Supplementary Table 1. Growth media were as previously described for *S. cerevisiae*^99^. All yeast strains were incubated at 30 °C (with shaking of liquid cultures at 230 rpm) for vegetative growth or at 23 °C for sporulation.

Gene deletion and tagging were performed using the PCR targeting method^100^. To increase transformation efficiency in *Sd. ludwigii*, 180-300 bp-long homology flanks were used, by using *S. cerevisiae* strain ESM356-1 for *in vivo* recombinational cloning of the DNA constructs or the NEBuilder HiFi DNA Assembly Cloning kit (New England Biolabs) for their *in vitro* assembly. High-fidelity DNA polymerases were always used for amplification of PCR cassettes for strain construction, whereas *Taq* DNA polymerases were routinely used for validation of engineered strains using colony PCR.

All plasmids used in this study are provided in Supplementary Table 7. They were maintained and propagated in *E. coli* strain DH5α and carried the ampicillin resistance cassette for selection in lysogeny broth (LB) supplemented with ampicillin (100 μg ml^-1^). All DNA oligonucleotides used for plasmid/strain construction are listed in Supplementary Table 8. Preparation and transformation of *E. coli* competent cells were performed using standard procedures^101^.

### Transformation of *Sd. ludwigii*

A lithium acetate (LiAc)-based method was optimized and used to transform *Sd. ludwigii*. For preparation of competent cells, a 50-ml YPD culture was inoculated from a saturated starter culture at an initial OD_600_ of 0.1 and was incubated at 30 °C (shaking at 230 rpm) until OD_600_ was in the range 0.8-1.0. Cells were harvested by centrifugation (500 × g for 5 min) and washed once in 25 ml of sterile distilled water and once in 12 ml of LiSorb buffer (100 mM LiAc, 1 M sorbitol, 10 mM Tris-HCl pH 8.0, 1 mM EDTA-NaOH pH 8.0) at room temperature. Cells were resuspended in 360 μl of LiSorb buffer, followed by the addition of 40 μl of denatured carrier DNA solution (salmon sperm ssDNA, 10 mg ml^-1^; Invitrogen). Aliquots of 50 μl of these competent cells were used immediately for transformation (for maximum efficiency) or stored for future use at -80 °C (no snap freezing). For each transformation, an aliquot of competent cells was thawed at room temperature and up to 20 μg of DNA (in max. 5 μl of water) were added. After mixing, 300 μl of buffer SdlLiPEG (50 mM LiAc, 45% w/v PEG-3350, 10 mM Tris-HCl pH 8.0, 1 mM EDTA-NaOH pH 8.0) were added, and the suspension was mixed well before being incubated at room temperature for 2 h. Following this incubation, the sample was subjected to heat shock at 38 °C for 40 min; alternatively, DMSO was added to a final concentration of 15% and heat shock was performed at 38 °C for 15 min. Subsequently, the cells were washed twice in YPD medium (4,000 rpm for 2 min), before resuspension in 2 ml of this medium and incubation at 30 °C for at least 6 h (with shaking at 230 rpm). Finally, cells were pelleted again, resuspended in 100 μl of YPD medium, plated on selective medium and incubated at 30 °C for 3-4 days. Three to six independent transformants were streaked to single colonies on fresh selective medium and the genomic alterations were validated by colony PCR.

### Mating and sporulation of *Sd. ludwigii*

Mating of *Sd. ludwigii* strains was performed by mixing equal numbers of haploid cells of opposite mating types on YPD plates and incubating them at 30 °C for 16 h. The cell mixture was subsequently transferred directly to sporulation medium or was streaked to single colonies on double selective YPD plates that support growth of diploid cells only (in case of strains containing different markers). When wild-type strains without suitable markers were mated, diploid colonies were identified by visual inspection, based on their brittle and matte morphology. For sporulation in liquid cultures, diploid cells were pre-grown in YPD medium to early stationary phase and then transferred to liquid sporulation medium (1% potassium acetate, 1% glucose, 0.25% yeast extract) at an initial density of OD_600_=0.6. The culture was incubated at 23 °C (with shaking at 230 rpm) for 3 days. For sporulation on plates, freshly grown cells from YPD plates were transferred to sporulation plates (2% w/v potassium acetate, 0.1% glucose, 2% agar) and incubated at 23 °C for up to 5 days.

### Construction of isogenic Sd. ludwigii strains of opposite mating types

Strain *Sd. ludwigii* NBRC 1722 (*MATalpha*) was used as the background strain for strain construction. To generate an isogenic *MATa* strain, a two-step replacement of the *MATalpha* locus was performed. Briefly, the *URA3* gene of strain NBRC 1722 was replaced with the *kanMX4* cassette. Then, the *MATalpha* locus was replaced with *URA3*, followed by replacement of the *URA3* marker with a cloned copy of the *MATa* locus (from strain NBRC 1723), using selection on 5-FOA. This resulted in strain Sdl-339. All strains and their genotypes are listed in Supplementary Table 1.

### *Sd. ludwigii* tetrad dissection

For tetrad dissection, a loopful of sporulated cells were transferred to 100 μl of sterile water and 1 μl of a zymolyase 100T solution (10 mg ml^-1^; Amsbio) was added. The suspension was gently mixed, incubated at room temperature for 10 min, and afterwards stored on ice. Droplets (15 μl) of this suspension were spread in the middle of YPD plates, which were used for tetrad dissection with a dissection microscope (Singer Instruments). Compared to *S. cerevisiae*, tetrads of *Sd. ludwigii* exhibit strong interspore bridges that link pairs of spores (dyads). In order to dissect tetrads, these bridges have to be broken mechanically. This can be tedious, and meticulous manipulation is necessary to prevent spore damage. To facilitate breakage of interspore bridges, the method developed by Yamazaki^38, 39^ was adopted. This involved placing a hydrated sterile cellophane sheet on the surface of the agar as a solid yet flexible support surface for dissection. Individual tetrads were carefully transferred onto it and spores were manipulated by lateral pushing rather than applying any pressure directly on them, since this could damage them. In addition, we used self-made glass needles with a little dent at one side in order to facilitate separation of dyads. Only with this setup, precise positioning of the dyads and exact application of forces to the bridges did we manage to dissect tetrads without damaging the spores. For analysis of mutants, we always dissected an equal number of control tetrads from wild-type cells.

### Pulsed-field gel electrophoresis (PFGE)

Spheroplasts were prepared from *Sd. ludwigii* cells, embedded in agarose plugs, lysed and processed for electrophoresis of intact chromosomes as previously described^39^, using the CHEF-DR II Pulsed Field Electrophoresis System (Bio-Rad).

### Transmission electron microscopy (TEM)

The glutaraldehyde-KMnO_4_ fixation method, followed by embedding in Agar 100, was used for electron microscopy of vegetative *Sd. ludwigii* cells as previously described^102^.

### Immunostaining of meiotic spreads

Spreads of *Sd. ludwigii* meiotic nuclei were prepared from sporulating cells in meiotic time courses using the procedure previously described for *S. cerevisiae*^103^ with some modifications. For spheroplast formation, 10 ml of a liquid sporulation culture were centrifuged at 700 × g for 5 min and the cell pellet was resuspended in 1 ml resuspension buffer (2% potassium acetate, 0.8 M sorbitol). Following the addition of 20 μl of a 0.5 M DTT solution and 25 μl of a zymolyase 100T (10 mg ml^-1^) solution, the sample was placed on a rotating wheel at 37 °C for digestion of cells; the process was monitored microscopically every 5 min. Cell shape changes during digestion; at first, cells become small and round, then their size increases by 2 to 3-fold and, finally, the cell wall disappears but the cell content is still held together by the plasma membrane. When at least 70% of the cells were at this stage, digestion was arrested by transferring the cells to a 15-ml tube containing 10 ml of ice-cold stop solution (0.1 M MES, 1 M sorbitol, 1 mM EDTA, 0.5 mM MgCl_2_, pH 6.4). The suspension was then centrifuged at 900 × g for 7 min (at 4 °C), the supernatant was discarded and the cell pellet was gently resuspended in 30 μl of ice-cold stop solution without sorbitol (absence of sorbitol causes hypotonic burst of the cells). This was followed by the addition of 60 μl of fixative (4% formaldehyde, 1.5% sucrose, pH 7.3) and 30 μl of the suspension were pipetted onto a microscope slide previously coated with 0.1% w/v poly-L-lysine. The suspension was spread by tilting the slide, which was followed by the addition of 80 μl of 1% Lipsol and mixing by tilting the slide again. Another 80 μl of fixative were added and the slide was tilted again for mixing. The slide was then dried overnight under a fume hood, and it was subsequently stored in a Coplin jar at -20 °C or processed directly for immunostaining.

Before immunostaining the slide was washed by immersion in PBS buffer for 10 min, which was followed by the addition of 100 μl of blocking buffer (0.5% BSA, 0.2% gelatin) and incubation for 10 min. The slide was covered with a coverslip during the incubation; to remove it before the next step of the procedure, the slide was immersed in PBS and gently shaken. For immunostaining, 40 μl of a primary antibody solution in blocking buffer were added, the slide was covered with a coverslip again and was incubated for 2 h in a humid chamber. The coverslip was then removed in PBS buffer and the slide was washed by immersion in PBS buffer for 5 min. Following that, 40 μl of a secondary antibody solution in blocking buffer were added on the slide, a coverslip was placed and the sample was incubated in a humid chamber for 1 h. The primary antibodies used were: rabbit anti-ScRfa1 (gift from E. Mancera), sheep anti-GFP (generated in our lab) at 1:500 dilution each, and rabbit anti-ScRad51 (gift from A. Shinohara) at dilution 1:1,000. The latter antibody104 was used for detection of discrete Rad51 foci, while the anti-GFP antibody was used for localization studies of GFP-tagged Rap1 in spreads of meiotic nuclei. The secondary antibodies used were: Alexa Fluor 488-conjugated donkey anti-rabbit IgG (Dianova), Cy2-conjugated donkey anti-sheep IgG (Dianova) and Cy3-conjugated donkey anti-rabbit IgG (Dianova), at 1:500 dilution each. For DNA staining, the coverslip was removed in PBS buffer and the slide was washed by immersion in fresh PBS buffer for 10 min (twice). The sample was then mounted with 20 μl of a Hoechst 33258 solution (0.5 μg ml^-1^) in 60% glycerol, a coverslip was added and sealed with nail polish. Single-plane epifluorescence images were acquired using an upright fluorescence microscope and a 100x Plan-Apochromat oil immersion objective. Image processing and merging of channels was performed using ImageJ^105^.

### Mutation accumulation experiment

Three Sd. ludwigii strains were used for the mutation accumulation study: the haploid reference strain NBRC 1722, the nearly isogenic diploid strain YLFP17-4 (homozygous diploid with the reference strain’s background) and the hybrid diploid strain YLFP188-1 (heterozygous diploid) (Supplementary Table 1). Clonal stocks of these parental strains were streaked on YPD plates, and randomly selected single colonies of each one served as founders of independent mutation accumulation lines (MALs). Spontaneous mutations were accumulated in 30, 19 and 11 MALs of the haploid, the homozygous and the heterozygous diploid strain, respectively, while they were growing under favorable conditions (YPD medium, at 30 °C). Each MAL was passed through a single-cell population bottleneck every 48 h by picking a random single colony and streaking it to single colonies on fresh YPD plates again. The plates were pre-marked with a target, and the single colony closest to the target was picked for each cycle, to ensure random colony selection. The experiment was performed for a total of 96-97 cycles, which corresponds to 2,037 generations for the haploid, 1,920 generations for the homozygous diploid, and 2,016 generations for the heterozygous diploid strain (each cycle corresponds to 20 generations for the homozygous diploid strain or 21 for the two other strains, as determined by resuspending 10 independent colonies of each one in water and counting cells using a Neubauer-improved hemocytometer, at the beginning of the experiment).

### Ploidy determination using flow cytometry

The cellular DNA content was analyzed by propidium iodide staining following ethanol fixation and flow cytometry on a FACSCanto II (BD Biosciences) instrument, according to standard protocols^106^.

### Total DNA extraction from *Sd. ludwigii* for next-generation sequencing

We optimized and used in this study two alternative methods that yielded total DNA of sufficient quality and quantity from *Sd. ludwigii*, as follows:

#### DNA isolation with commercial kits

Liquid *Sd. ludwigii* cultures in YPD medium (early stationary phase) were processed for DNA extraction using the Genomic-tip 20/G (mini prep, starting with approx. 10^8^ cells) or 100/G (midi prep, 10^9^ cells) kit (QIAGEN), according to the manufacturer’s recommendations. An additional washing step with buffer QC was performed before the final elution step. Elution buffer QF was pre-warmed to 50 °C for maximum yield. To recover DNA following isopropanol precipitation, spooling with a pipette tip was preferably used, or alternatively DNA was pelleted by centrifugation at 4,500 × g for 40 min (at 4 °C). A washing step was performed using 500 μl of ice-cold 70% ethanol. Following centrifugation (13,000 rpm for 10 min at 4 °C), ethanol was completely removed, the sample was air-dried at room temperature for 10 min, and DNA was then resuspended in 100 μl (mini prep) or 200 μl (midi prep) TE buffer (pH 8.0). To further improve the quality of the isolated DNA, 100 μl of each DNA sample were further processed using the DNeasy PowerClean Pro Cleanup kit (QIAGEN), according to the manufacturer’s instructions. Total DNA was finally eluted in 50 μl of elution buffer (10 mM Tris-HCl, pH 8.0) and stored at -20 or -80 °C.

#### Alternative DNA isolation method

Cells were harvested from a liquid culture in YPD medium (OD_600_ = 1), washed in 1 ml of resuspension buffer (0.9 M sorbitol, 0.1 M EDTA pH 7.5), pelleted again by centrifugation and finally resuspended in 0.4 ml of resuspension buffer amended with 1.4 mM β-mercaptoethanol. Subsequently, 20 μl of a zymolyase 100T solution (10 mg ml^-1^) were added, and the suspension was incubated at 37 °C for 30 min. Digested cells were layered over a sorbitol cushion (1.8 M) in a 50-ml tube, which was then centrifuged at 2,000 rpm for 15 min (at 4 °C). Sorbitol was discarded and the pellet was resuspended in 0.4 ml TE buffer (pH 8.0) after the addition of 90 μl of a solution containing 0.5 M EDTA, 2 M Tris and 10% SDS (pH 8.0). The sample was incubated at 65 °C for 30 min before the addition of 80 μl of a 5 M potassium acetate solution and incubation at 4 °C for at least 1 h (usually overnight). The sample was then centrifuged at 13,000 rpm for 15 min and the supernatant was mixed with 1 ml of ethanol. The precipitate was recovered by brief centrifugation (5 s) at maximum speed, and the pellet was then rinsed with 70% ethanol, air-dried and gently resuspended in 0.5 mL TE buffer (pH 8.0). The sample was centrifuged again (13,000 rpm for 15 min) to remove any insoluble material, and 5 μl of an RNase A solution (5 mg ml^-1^) were added to the supernatant, which was followed by incubation at 37 °C for 30 min. Two consecutive phenol-chloroform-isoamyl alcohol (25:24:1) extraction steps followed (the 5Prime Phase Lock Gel system was used for maximum recovery), and 0.6-0.8 volumes of isopropanol were added for DNA precipitation by centrifugation (13,000 rpm for 10 min, at 4 °C). Total DNA was washed with 70% ethanol, dried and resuspended in 50 μl molecular grade water. For removal of polysaccharides, the DNA sample was mixed with 0.1 volume of a hexaamminecobalt(III) chloride solution (100 mM) and centrifuged at 13,000 rpm for 5 min. The pellet was washed with 200 μl of water and dissolved in 50-100 μl of exchange buffer (100 mM EDTA pH 8.0, 2 M guanidinium thiocyanate) at 37 °C for 1 h. Once the pellet was completely dissolved, total DNA was precipitated, washed, dried and finally resuspended in 50 μl TE buffer (pH 8.0).

Agarose gel electrophoresis, UV spectrophotometry and fluorometry (Qubit 4.0, using the Qubit dsDNA HS Assay Kit; Invitrogen) were used for the assessment of the quality and quantity of the isolated DNA. During optimization of the extraction protocols, DNA quality was characterized using the Fragment Analyzer (with the Genomic DNA 50 kb kit; Agilent).

### Next-generation sequencing (NGS)

Short-read (Illumina) and long-read (Pacific Biosciences, Oxford Nanopore Technologies) sequencing methods were used in this study for addressing different questions. Library preparation workflows and sequencing procedures are summarized below.

#### Illumina sequencing

For sequencing on an in-house Illumina NextSeq instrument, libraries were prepared using the NEBNext Ultra II FS DNA Library Prep Kit for Illumina (New England Biolabs), using the appropriate protocols for DNA samples of 50 ng (lower input, without size selection) or 250 ng (higher input, with size selection), according to the manufacturer’s recommendations. Enzymatic fragmentation of DNA was controlled to generate fragment size distributions in the range of 300-500 bp. For PCR enrichment of adaptor-ligated DNA, 6 or 7 amplification cycles were used for samples with higher and lower starting amounts of DNA, respectively. Multiple DNA samples were barcoded with unique dual indices using the NEBNext Multiplex Oligos for Illumina (96 Unique Dual Index Primer Pairs) (New England Biolabs) and pooled at equimolar concentrations. The NEBNext Library Quant Kit for Illumina (New England Biolabs) was used for the qPCR-based quantification of the pooled library, which was then loaded on a high-output flow cell (NextSeq 500/550 High Output Kit v2.1, 300 cycles; Illumina) for sequencing on an Illumina NextSeq 550 sequencer, in paired-end mode (2×150 bp).

Alternatively, Illumina sequencing was performed on MiSeq and HiSeq 2000 instruments, at the EMBL Genomics Core Facility (Heidelberg, Germany), in paired-end mode (2×150 and 2×100 bp for MiSeq and HiSeq systems, respectively). Sequencing libraries were prepared by the Deep Sequencing Core Facility (University of Heidelberg, Germany). Briefly, mechanical shearing using a Covaris ultrasonicator was used for DNA fragmentation to a size range of 300-500 bp. The NEBNext Ultra II DNA Library Prep Kit for Illumina (New England Biolabs) was used for library preparation, and 8 PCR cycles were performed for amplification of adaptor-ligated DNA. The quality and quantity of libraries were assessed by using a Qubit fluorometer (Invitrogen) and a Bioanalyzer instrument (Agilent).

#### Pacific Biosciences SMRT (single-molecule real-time) sequencing (PacBio)

For the generation of the de novo genome assembly of our reference Sd. ludwigii strain we used PacBio sequencing, which was performed at GATC Biotech AG (now Eurofins Genomics). Library preparation briefly involved DNA fragmentation, size selection (using a BluePippin instrument; Biozym Scientific), DNA end repair and adaptor ligation, annealing of the primer and the polymerase. The PacBio library was then sequenced on 2 SMRT cells in a PacBio RS II instrument (run mode: 240 min. movie) using the P6-C4 chemistry. This yielded a total of 2.52 Gb of pass-filter sequence data, which corresponds to an average genomic sequencing depth of approx. 200x.

#### Oxford Nanopore Technologies sequencing (ONT)

Generation of genome assemblies of other *Sd. ludwigii* strains was based on ONT sequencing on a MinION sequencer. The Ligation Sequencing Kit (for 1D experiments; SQK-LSK109) was used for library construction, starting from 1 μg of total DNA of each sample, and different samples were barcoded using the Native Barcoding Expansion 1-12 (PCR-free) kit (both kits were purchased from Oxford Nanopore Technologies) and pooled for multiplexed sequencing (according to the manufacturer’s instructions). No fragmentation of the input DNA was performed. The sequencing libraries were loaded on R9.4.1 flow cells and sequenced on a MinION device. Each strain was sequenced until the desired amount of sequencing data was gathered, corresponding to average genomic sequencing depths of approx. 30-40x.

### Bioinformatics

#### Genome assemblies

The *de novo* genome assembly of the reference *Sd. ludwigii* strain NBRC 1722 was based on high-coverage PacBio sequencing data. A total of 193,117 reads (totalling 2,518,671,875 bp and corresponding to an average genomic sequencing depth of approx. 200x) with mean length 13,041 bp and mean q-score 0.827 passed the filter (SMRT Portal, Pacific Biosciences). The PacBio HGAP.3 pipeline (Hierarchical Genome Assembly Process) within SMRT Analysis v2.3.0 (Pacific Biosciences) was used for the assembly (mapping single-pass reads to seed reads, generating and quality trimming pre-assembled consensus sequences, assembling long high-quality reads using the Celera Assembler^107^ and consensus polishing using Quiver^108^). One of the resulting scaffolds (with length 22,745 bp and GC content 11.7% was determined using BLAST+ v2.8.0^109^ to be part of the mitochondrial genome and was removed from the nuclear assembly. Subsequently, we used an Illumina MiSeq 2×150-bp paired-end read dataset generated from the same DNA sample for further polishing of the assembly using Pilon^110^ (version 1.22), to generate the final assembly version (with an N50 value of 1,848,403 bp). The genome is organized in 7 chromosomes, with a total size of 12,500,424 bp and an overall GC content of 30.9%. Chromosome B is interrupted by the single rDNA locus, which is estimated to be at least 100 kb-long (based on the PFGE profile of the strain) and is represented by the only gap in the assembly. Only one small (26,468 bp-long) subtelomeric scaffold (“scaffold1”) could not be unambiguously linked to any of the chromosomal scaffolds, but it is presumed to be part of the right extremity of chrG (based on repetitive subtelomeric content analysis, gene synteny and co-segregation in meiosis).

For whole-genome comparisons of the reference genome with those of *Sd. ludwigii* strains 122 and NBRC 1723, we used ONT sequencing and generated *de novo* assemblies of their genome sequences. For this, we used 93,607 filter-passing reads (totalling 407,656,509 bp and corresponding to an average genomic sequencing depth of approx. 33x) with an N50 read length of 6.89 kb and a mean q-score of 13.242 for strain 122, and 102,478 filter-passing reads (totalling 526,253,091 bp and corresponding to a genomic depth of approx. 42x) with an N50 read length of 7.76 kb and a mean q-score of 13.586 for strain NBRC 1723. We used the GPU-accelerated base-caller Guppy (v2.3.5; Oxford Nanopore Technologies) with the high-accuracy model (“flip-flop” algorithm) for base-calling. Demultiplexing of barcoded reads and trimming of ONT adaptors were performed using Porechop (v0.2.4; https://github.com/rrwick/Porechop). Genome assembly were then constructed using SMARTdenovo (https://github.com/ruanjue/smartdenovo), which generates assemblies from all-vs-all raw read alignments, with parameters “-c 1 -k 14 -J 500 -e zmo”. Scaffolds of the strain 122 assembly were polished using an Illumina MiSeq 2×150-bp paired-end sequencing dataset of the same strain and Pilon^110^ (v1.23). Subsequently, the assemblies were scaffolded further using ONT long read information with SSPACE-LongRead^111^ (v1-1) and, in the case of strain 122, gaps were closed using GapFiller^112^ (v1-10) and the previously mentioned paired-end Illumina MiSeq reads. The NBRC 1723 assembly was corrected by mapping the raw ONT reads using minimap2^113^ (2.17) and then using Racon^114^ (v1.3.3). The final assemblies were generated after a second round of scaffolding using SSPACE-LongRead (v1-1), followed by polishing using Pilon (v1.23) for strain 122 or Nanopolish^115^ (v0.11.0) for strain NBRC 1723. The assemblies consist of 18 scaffolds (12,368,365 bp, N50 = 1,655,802 bp) for strain 122 and 10 scaffolds (12,251,049 bp; N50 = 1,643,865 bp) for strain NBRC 1723.

#### Whole-genome comparison (alignment)

For genomic comparisons between *Sd. ludwigii* strains we used the software MUMmer^116^ (v4.0.0). Alignments of genomic sequences were performed using the program nucmer (with the --maxmatch option). The output files were passed on to the script mummerplot for visualization. Comparisons between genomes were plotted as chord diagrams using the R package circlize^117^ (v0.4.6). The dnadiff (v1.3) wrapper of the genome alignment system MUMmer^116^ was used for the calculation of average nucleotide identity between whole genomes. Structural variations were called using Sniffles^118^ (v1.0.10) from the sorted output of the BWA-MEM aligner^119^ (v0.7.17).

#### Gene prediction and annotation

For the generation of a comprehensive *Sd. ludwigii* gene annotation dataset, we used the reference genome sequence and combined the results from different tools and pipelines, followed by extensive manual curation. Firstly, we used the Yeast Genome Annotation Pipeline (YGAP^120^), which is based on homology and gene synteny information from previously analyzed yeast species, available in the Yeast Gene Order Browser (YGOB) database^121^. Secondly, AUGUSTUS^122^ (v3.2.1) was trained on the latest (as of September 2018) available protein datasets of several annotated yeast species (*Saccharomyces cerevisiae*, *Naumovozyma dairenensis*, *Tetrapisispora blattae*, *Vanderwaltozyma polyspora*, *Kluyveromyces lactis*, *Eremothecium gossypii*, *Hanseniaspora uvarum*) and all resulting training datasets were independently used for *ab initio* gene prediction in *Sd. ludwigii*. Thirdly, the MAKER pipeline^123^ (v2.30), with predictors Augustus^122^ (v2.5.5), Genemark-ES^124^ (v2.3) and SNAP^125^, was used for gene prediction using the protein datasets of all yeast species available in the YGOB database^121^ as evidence. Custom scripts were then used for comparing the aforementioned prediction datasets and integrating all non-overlapping predicted gene models into a union dataset. The most probable gene model was retained for each gene, based on manual curation (using the genome browser IGV^126^; v2.6.2) and BLAST+^109^ (v2.8.0) comparisons. Finally, all genomic regions without predicted genes were extracted from the genomic sequence (using BEDtools^127^; v2.27.0), and custom scripts were used for retrieving all open reading frames in these regions that were predicted to code for proteins with a minimum size of 30 aa. These were analysed using BLAST+^109^ (v2.8.0) and were retained in the final gene prediction dataset only if they were homologous to known proteins of other yeast species or similar to conserved hypothetical proteins. Furthermore, we used tRNAscan-SE^128^ (v2.0) for the identification of tRNA genes, BLAST+^109^ (v2.8.0) and RNAmmer^129^ (v1.2) for the annotation of rRNA genes, and the program cmscan (v1.1) of Rfam^130^ (v12.1) for prediction of other RNA genes. For the assessment of gene prediction completeness, BUSCO^131^ (v3) was used, against the “Saccharomycetales” and “Fungi” databases of conserved single-copy orthologs. Functional annotation and Gene Ontology (GO) assignment of the final gene dataset was performed using the software suite Blast2GO PRO^132^ (v5), integrating evidence from BLAST+^109^ (v2.8.0), InterProScan^133^ (v5) and eggNOG^134^ (v4.5) searches, and manual curation for resolving conflicting results. The consensus features of point centromeres of Saccharomycetaceae representatives, that were compared to the *Sd. ludwigii* predictions, were retrieved from Gordon *et al*.^135^.

#### Analysis of meiotic gene content

All *S. cerevisiae* genes that are known to be involved in meiosis and recombination were retrieved from the Saccharomyces Genome Database (SGD^136^, https://www.yeastgenome.org) by searching for genes with relevant GO terms. These searches retrieved a total of 284 genes, which were then used as queries for the identification of homologs in the *Sd. ludwigii* gene annotation dataset using BLAST+^109^ (v2.8.0) analyses. For all genes that remained undetected after similarity searches, we performed detailed gene synteny comparisons with annotated Saccharomycetaceae species that are included in the Yeast Gene Order Browser (YGOB^121^) database (v7; http://ygob.ucd.ie). The meiotic gene complement of *Sd. ludwigii* was compared to that of the 20 Saccharomycetaceae species that are included in the YGOB (v7) database, as well as to that of 9 Pichiaceae species that are included in the Methylotroph Gene Order Browser^137^ (http://mgob.ucd.ie). In order to gain a better understanding of how the missing meiotic genes from *Sd. ludwigii* evolved along the yeast phylogeny, we also studied the distribution of their homologs in the families Saccharomycodaceae and Phaffomycetaceae. For this, we used the *S. cerevisiae* protein sequences as queries in tBLASTn^109^ searches (with the Expect threshold set at 50) against all available genome assemblies from these two families in the NCBI Genome database on the 18th August 2019 (i.e. 28 genome assemblies from 18 *Hanseniaspora*/*Kloeckera* species or unclassified strains from the Saccharomycodaceae, and 42 assemblies from 33 species/strains of the Phaffomycetaceae, excluding the *Komagataella* species that have been classified in the Pichiaceae clade^138^). Representative positive hits were further examined using reciprocal BLASTp searches against the annotated *S. cerevisiae* proteome (SGD^136^). In those cases of putative homologs with marginally detectable similarity (often limited to short protein regions), synteny with *S. cerevisiae* S288C (YGOB database; v7) was also investigated and used as additional evidence for inferring homology. To confirm the absence of detectable homologs of *MER1* from the *Hanseniaspora* species (family Saccharomycodaceae), we used the synteny information from *S. cerevisiae* to identify the expected positions in the genomes of 10 representative species (i.e. *H. vineae*, *H. osmophila*, *H. occidentalis*, *H. lachancei*, *H. pseudoguilliermondii*, *H. guilliermondii*, *H. opuntiae*, *H. thailandica*, *H. jakobsenii* and *H. singularis*) and we analysed all predicted ORFs in these regions for weak similarity to the *S. cerevisiae* Mer1 sequence. The online search engine of the Pfam database^139^ (v32.0; https://pfam.xfam.org) was used for the identification of protein domains. Prediction of intrinsically unstructured regions in proteins was performed with the online tool IUPred2A^140^ (https://iupred2a.elte.hu).

#### Read mapping and variant calling

Reads were mapped to the *Sd. ludwigii* reference genome (strain NBRC 1722), which was previously masked with RepeatMasker (v4.0.7, default parameters; http://www.repeatmasker.org), using bwa mem^141^ (v0.7.17). Resulting bam files were sorted and indexed using SAMtools^142^ (v1.9). Duplicate reads were marked and sample names were assigned using Picard (v2.18.14; https://broadinstitute.github.io/picard/). The GATK pipeline^143^ (v3.7.0) was used to realign remaining reads. Variants were then called using GATK UnifiedGenotyper. Calling was performed simultaneously for all spores from the same tetrad or all lines from the same background.

#### Identification of recombination events

For the single-nucleotide polymorphism (SNP) segregation analysis of the hybrid cross, SNPs called from the respective reads mapping were first filtered (bcftools view; v1.9, https://github.com/samtools/bcftools) in order to define a set of high-confidence discriminant markers. Only positions with a single alternate allele, supported by at least 10 reads across both parents and with >90% of the reads covering either the reference or alternate allele, were selected. For each tetrad of the cross, SNPs located at the aforementioned marker positions were extracted, and the parental origin was assigned based on SNP correspondence between parents and spores at those positions. The result was formatted as a Seg file and used as input of the CrossOver pipeline (ReCombine suite)^144^, using default parameters and modified chromosome sizes/coordinates to match the reference genome of *Sd. ludwigii*. The reported events were individually validated by visual inspection using the Integrative Genomics Viewer (IGV). The same method was applied for the SNP segregation analysis of the South African cross, except that the minimum amount of reads supporting a marker position was lowered to 3 for each parent, as the coverage was reduced for one of the parents.

#### Mutation accumulation analysis

Filters were applied to the SNPs called from the mutation accumulation lines to identify SNPs that were present only in a single line, as we expected these events to be line-exclusive. First, only positions covered by more than 10 reads in each sample with a single alternate allele were kept. Then, filters based on the numbers of lines per background and the type of conversion event expected to occur in a given background (homozygous to homozygous, homozygous to heterozygous, heterozygous to homozygous) were applied. Heterozygous SNPs were only retained if their allele balance was between 0.4 and 0.6. In the case of heterozygous-to-homozygous substitution, likely LOH events were filtered out to prevent false-positive SNP calls. The base-substitutional mutation rate (*μ*_bs_) of each strain genealogy was calculated as the ratio of the total number of line-exclusive novel mutations divided by the size of the analyzed fraction of the genome (after masking), the total number of generations and the number of respective lines. The 95% Poisson confidence intervals of mutation rates were computed as in Long *et al*.^145^, and the mutation bias (corrected for the genomic GC content) as well as the theoretical equilibrium GC level (under mutation pressure alone) were calculated as in Krasovec *et al*.^146^.

#### Population analyses

A total of 558,629 biallelic segregating sites were used to construct the neighbor-joining trees using the R^147^ packages ape and SNPrelate. The .gvcf matrices (of the whole genome or individual chromosomes) were converted into .gds files. Individual dissimilarities were estimated using the snpgdsDiss function, and the bionj algorithm was run on the distance matrices. Tree comparison was performed using the Visual TreeCmp package^148^ (web application; v2.0.76), using 7 unrooted metrics (topological or weighted). For normalization of distances, the results of the topological metrics were compared to the average values of 1,000 randomly generated tree pairs (uniform average method). The software PopLDdecay^149^ (v3.41) was used for the calculation of the LD decay data, which were then smoothed (moving average method) and plotted using R^147^.

#### Other genomic analyses

We used RepeatMasker (v4.0.7; http://www.repeatmasker.org) with combined Dfam^150^ (v2.0) and Repbase^151^ (Genetic Information Research Institute - GIRI) databases, NCBI RMBLAST^109^ (v.2.6.0+) as the search engine and Tandem Repeats Finder^152^ (v4.0.9), for identification of genomic repeats, low-complexity regions and transposable elements. Other genomic data analyses were performed using R^147^ and the Bioconductor framework^153^.

## Extended Data - Figures

**Extended Data Fig. 1:**
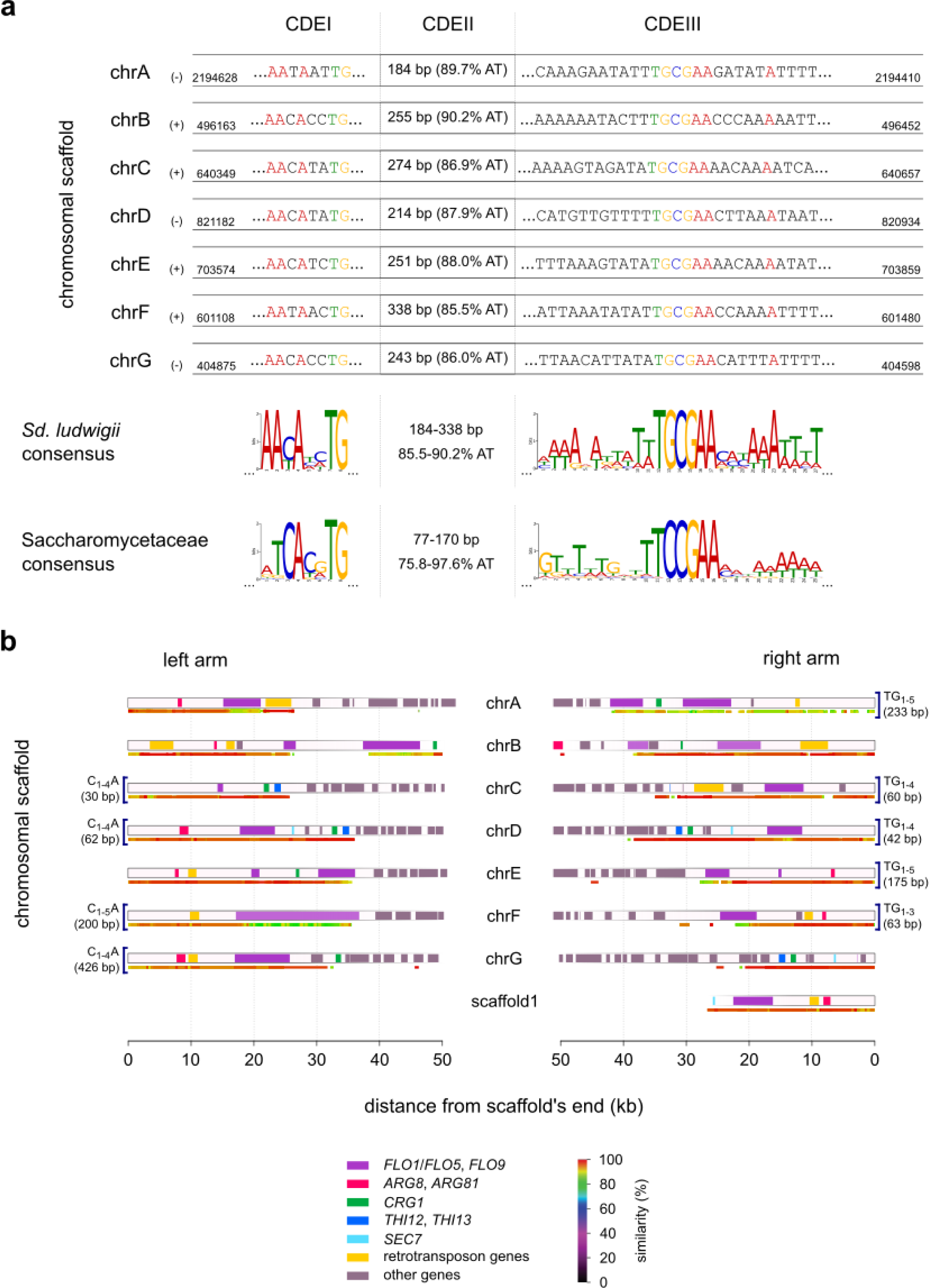
Features of *Sd. ludwigii* point centromeres and repetitive (sub)telomeres. **a**, Sequences of the centromere DNA elements (CDEs) I and III, and length and AT content of CDE II of each predicted centromere. The direction of each centromere with reference to the genomic scaffold (+/-) is provided next to its bar (left). The consensus DNA motif sequences of CDEs I and III and the ranges of length and AT content of CDEs II are shown (bottom), in comparison to the respective Saccharomycetaceae data. **b**, The terminal regions of all chromosomal scaffolds are shown. A horizontal line beneath each bar indicates coverage by subtelomere-specific DNA repeats that are shared between different subtelomeres. These coverage bars are color-coded on the basis of the highest sequence similarity to other subtelomeres, according to the color key (bottom right). Detected telomere-specific DNA motif sequences at the ends of chromosomal scaffolds are indicated by square brackets. Gene content/locations for each subtelomere are shown as boxes within the bars, colored according to the respective code (bottom left). Gene homologs typical of *S. cerevisiae* subtelomeres, such as *FLO* and *ARG* gene family members, are present in all *Sd. ludwigii* subtelomeres, while retrotransposons are also present in most of them.

**Extended Data Fig. 2:**
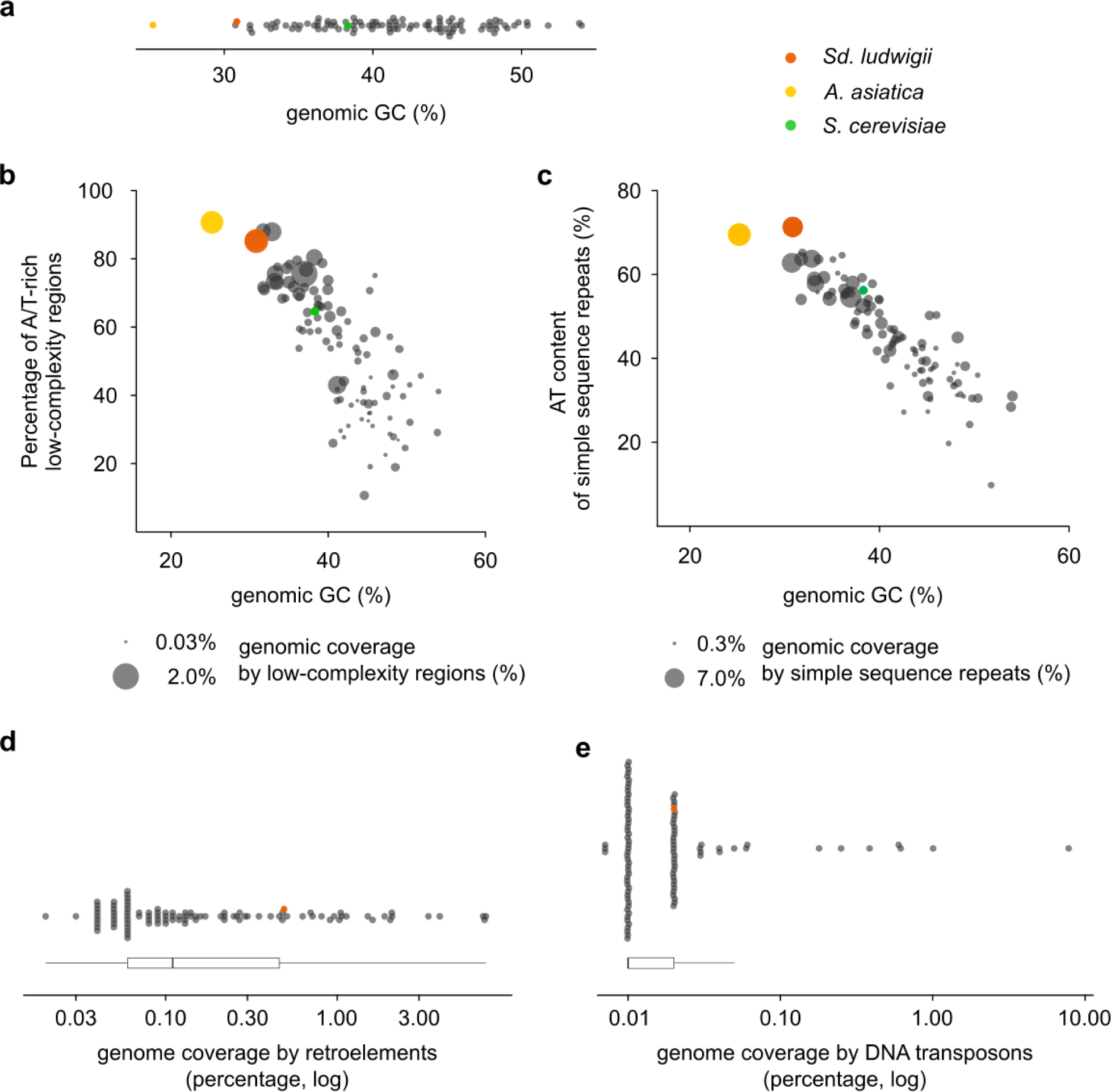
A comparative genomic analysis with 100 yeast and other fungal genomes revealed that the *Sd. ludwigii* genome is exceptionally AT-rich and enriched in low-complexity, repetitive, and transposable elements. **a**, Overall genomic GC content. **b**, Genomic coverage by low-complexity regions (dot width) and the proportion of those regions that are AT-rich (>70%; y axis) versus the overall genomic GC content (x axis). **c**, Genomic coverage by simple sequence repeats (SSRs or microsatellites; dot width) and the total AT content of SSRs (y axis) versus the mean genomic GC content (x axis). **d and e**, Genomic coverage by retroelements (d) and DNA transposons (e). All genomes included in the comparisons here are listed in Supplementary Table 2.

**Extended Data Fig. 3:**
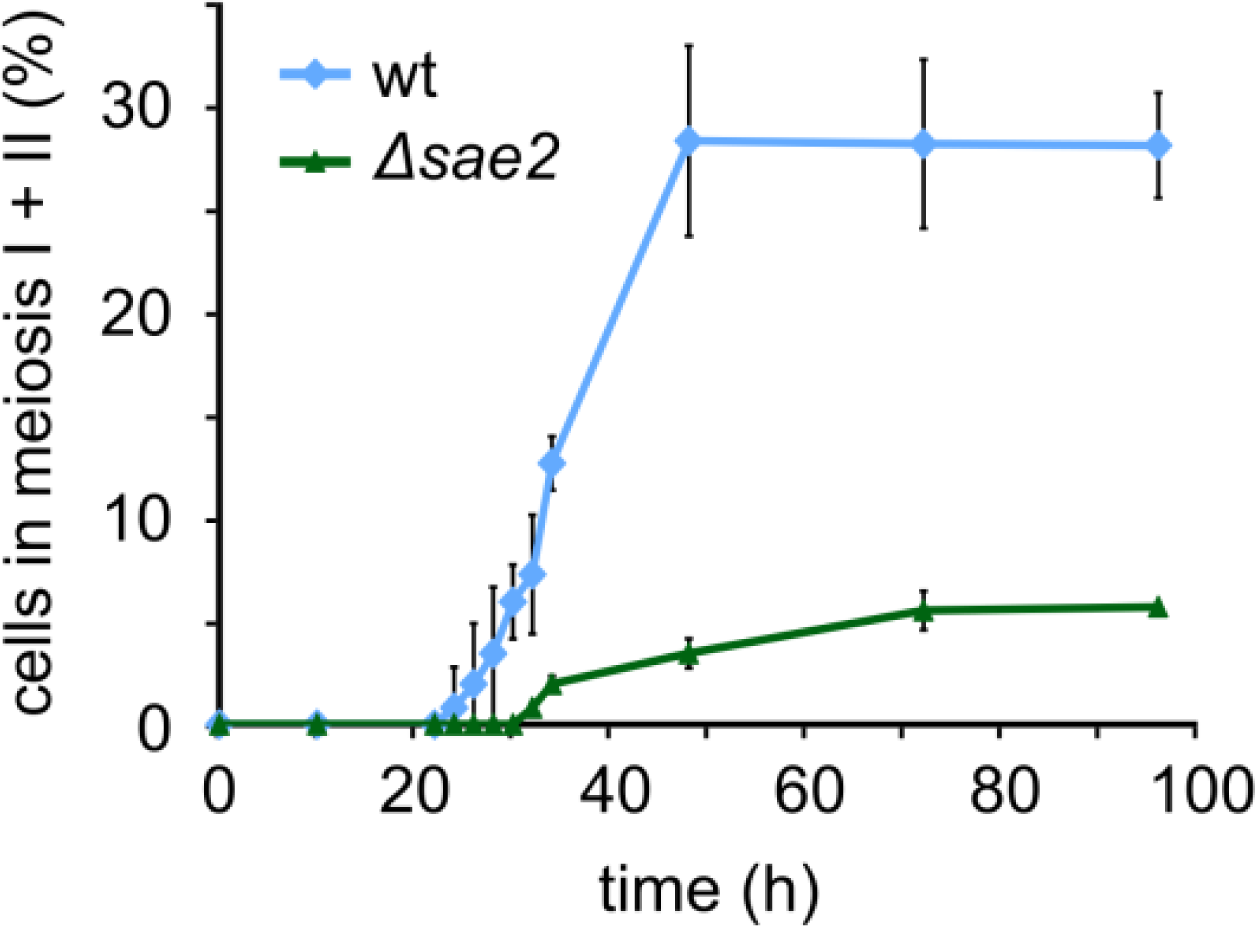
Sd. ludwigii Sae2 is necessary for normal sporulation, similarly to Spo11. Meiotic time-course analysis of a homozygous Δ*sae2* deletion strain in comparison to the wild type. Following induction of cells to enter meiosis, samples were withdrawn at the indicated time points and their cellular DNA content was stained with Hoechst 33258 to determine binucleate (meiosis I) and tetranucleate (meiosis II) cells. Error bars: SD (3 replicates).

**Extended Data Fig. 4:**
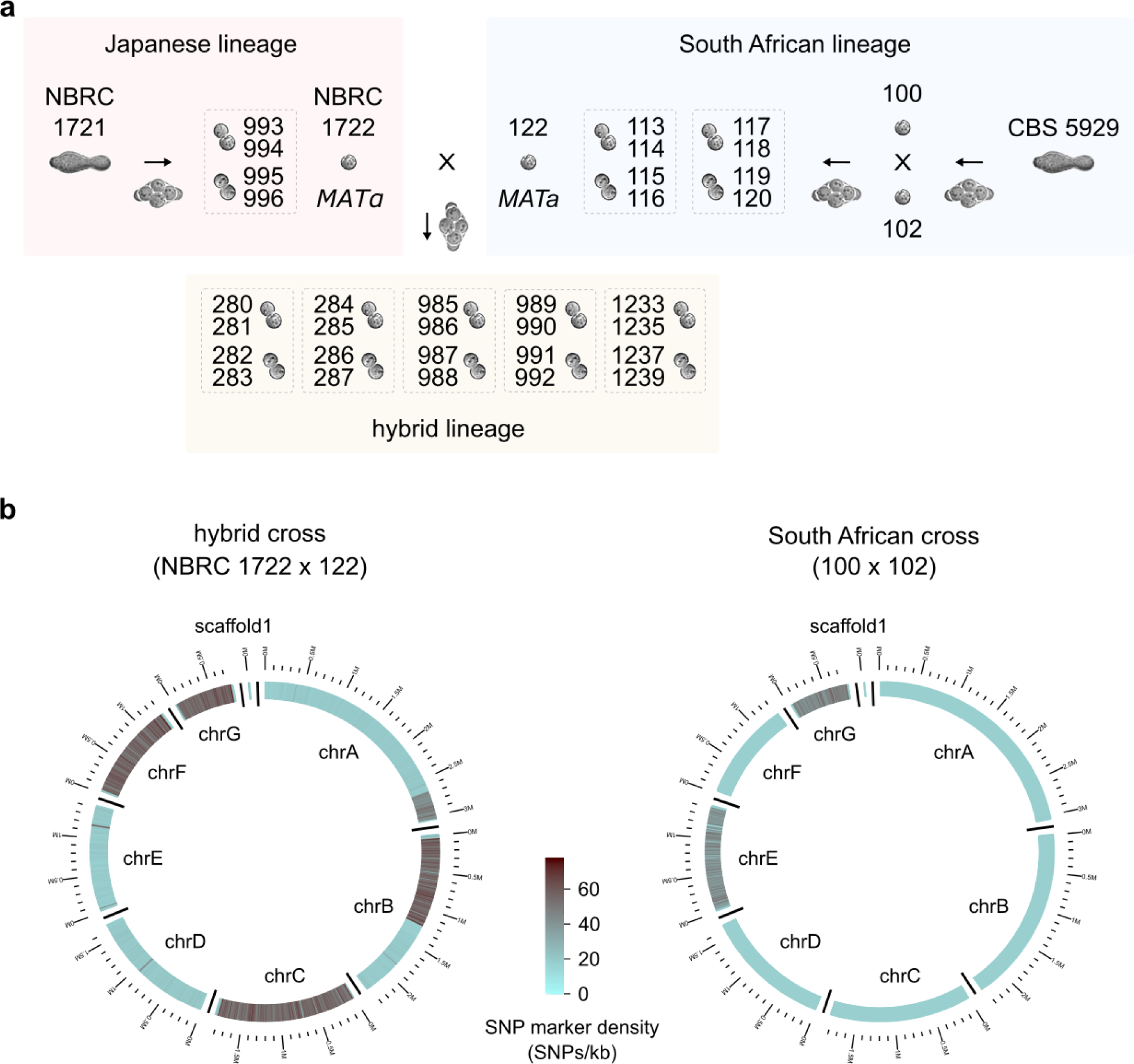
Meiotic SNP segregation analysis in *Sd. ludwigii*. **a**, A detailed scheme of the experimental setup, including the identifiers of all strains used: the wild-type isolates are referred to with their original strain collection IDs, whereas the Knop lab *Sd. ludwigii* strain collection IDs are provided for all other strains that were generated in this study (Supplementary Table 1). Spore viability of strain CBS 5929 was very low (probably due to high load of deleterious mutations); it was thus impossible to obtain full viable tetrads from this strain for the SNP segregation analysis. We used backcrossing to isolate two spores (spores 100 and 102) that gave rise to diploids with high spore viability. **b**, Heterozygosity levels (expressed as the number of filtered high-quality SNPs per kb of genomic sequence) between the parental strains of each cross are plotted here along the chromosomes of the reference genome sequence.

**Extended Data Fig. 5:**
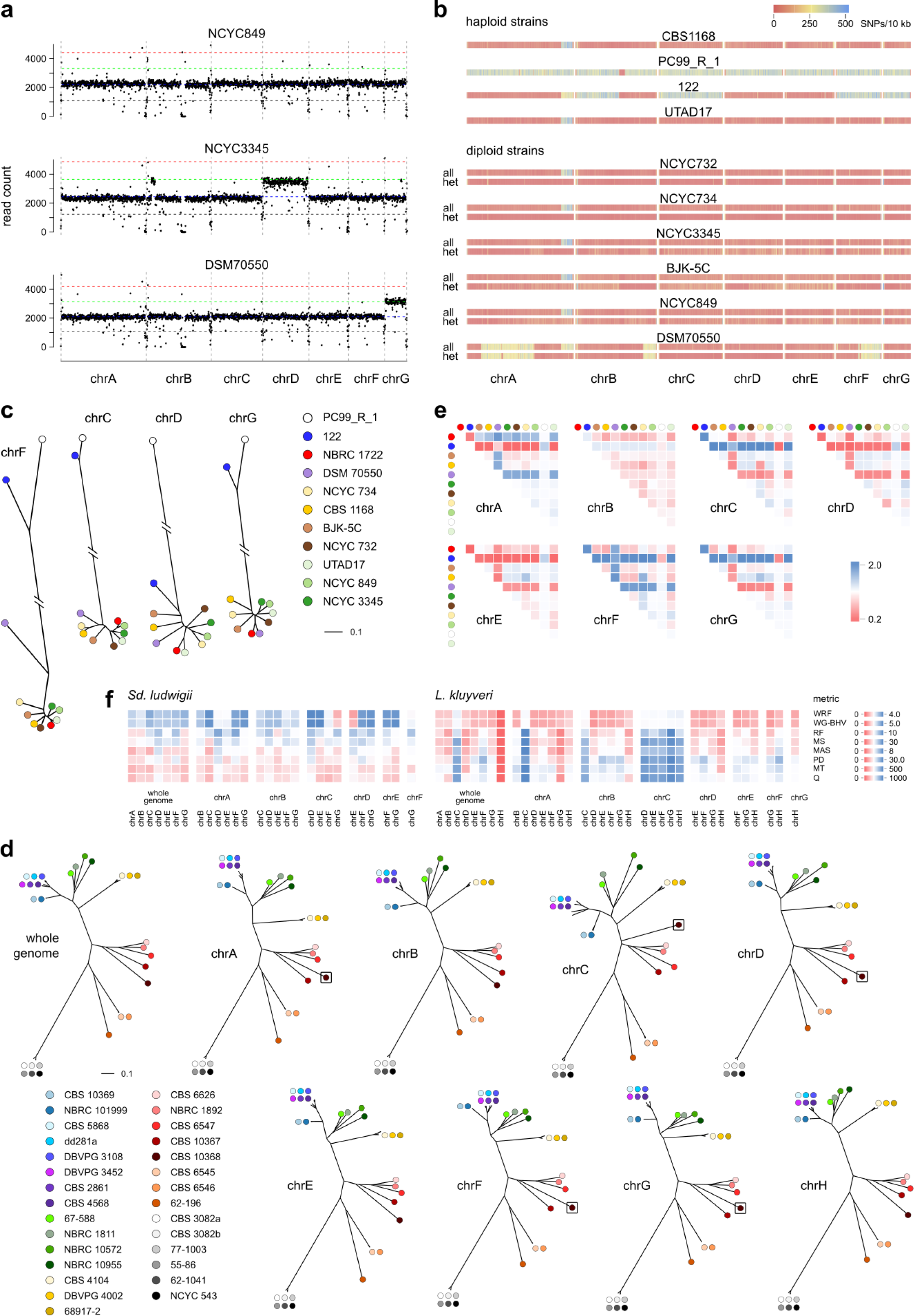
Genomic comparisons of Sd. ludwigii strains. **a**, Examples of euploid (NCYC 849) and aneuploid (chrD and chrG trisomies in NCYC 3345 and DSM 70550, respectively) isolates of *Sd. ludwigii*, determined by read coverage analysis. The gap near the middle of chrB corresponds to the rDNA region. **b**, Genome-wide distribution of SNPs of all *Sd. ludwigii* strains in comparison to the reference strain. For each diploid strain, the total number of SNPs (“all”) is shown in the upper bar and the heterozygous positions (“het”) in the bottom one. **c**, Dendrograms of *Sd. ludwigii* strains, constructed by hierarchical cluster analysis of SNPs of individual chromosomes. The dendrograms of the remaining chromosomes, as well as that of the whole genome, are shown in Fig. 6a. **d**, Similarly constructed dendrograms of *L. kluyveri* strains. **e**, Differences in SNP density of individual chromosomes of *Sd. ludwigii* in comparison to the whole genome, for all strains (color code as in c). **f**, Tree difference scores (seven unrooted metrics, no normalization) between all combinations of whole-genome SNPs and SNPs of individual chromosomes, for *Sd. ludwigii and L. kluyveri.* The analyzed strains are shown in panels (c) and (d), respectively. WRF: weighted Robinson-Foulds distance. WG-BHV: weighted geodesic (BHV) unrooted distance. RF: Robinson-Foulds distance. MS: matching split distance. MAS: unrooted maximum agreement subtree distance. PD: path difference distance. MT: matching triplet distance. Q: quartet distance.

**Extended Data Fig. 6:**
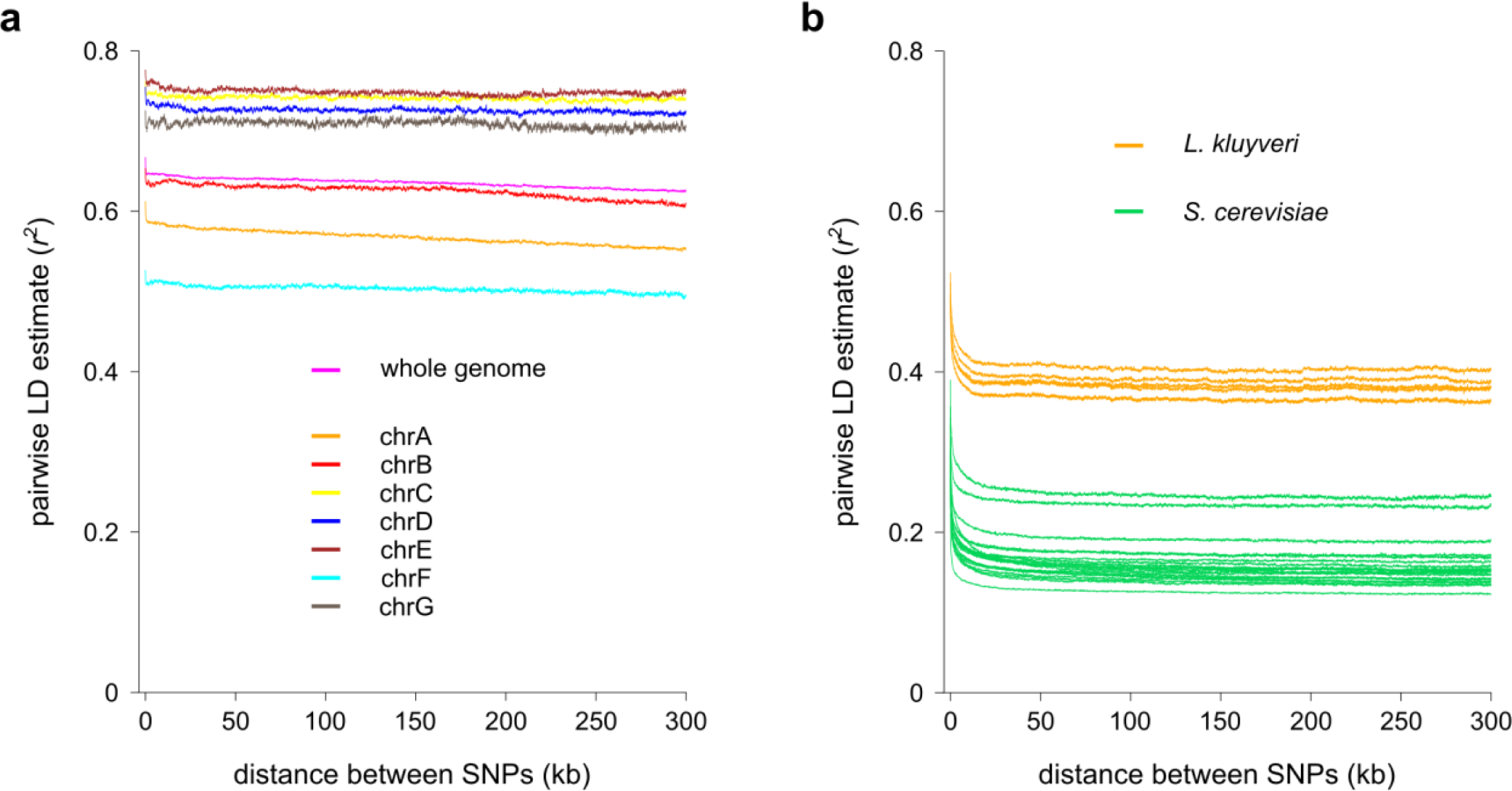
Curves of linkage disequilibrium (LD) decay as a function of physical distance between SNP marker pairs. **a**, LD decay curves of the whole genome in comparison to individual chromosomes of *Sd. ludwigii*. **b**, LD decay curves of 5 *L. kluyveri* and 20 *S. cerevisiae* groups of strains (of the same sample size, n=11).

**Supplementary Table 1:**
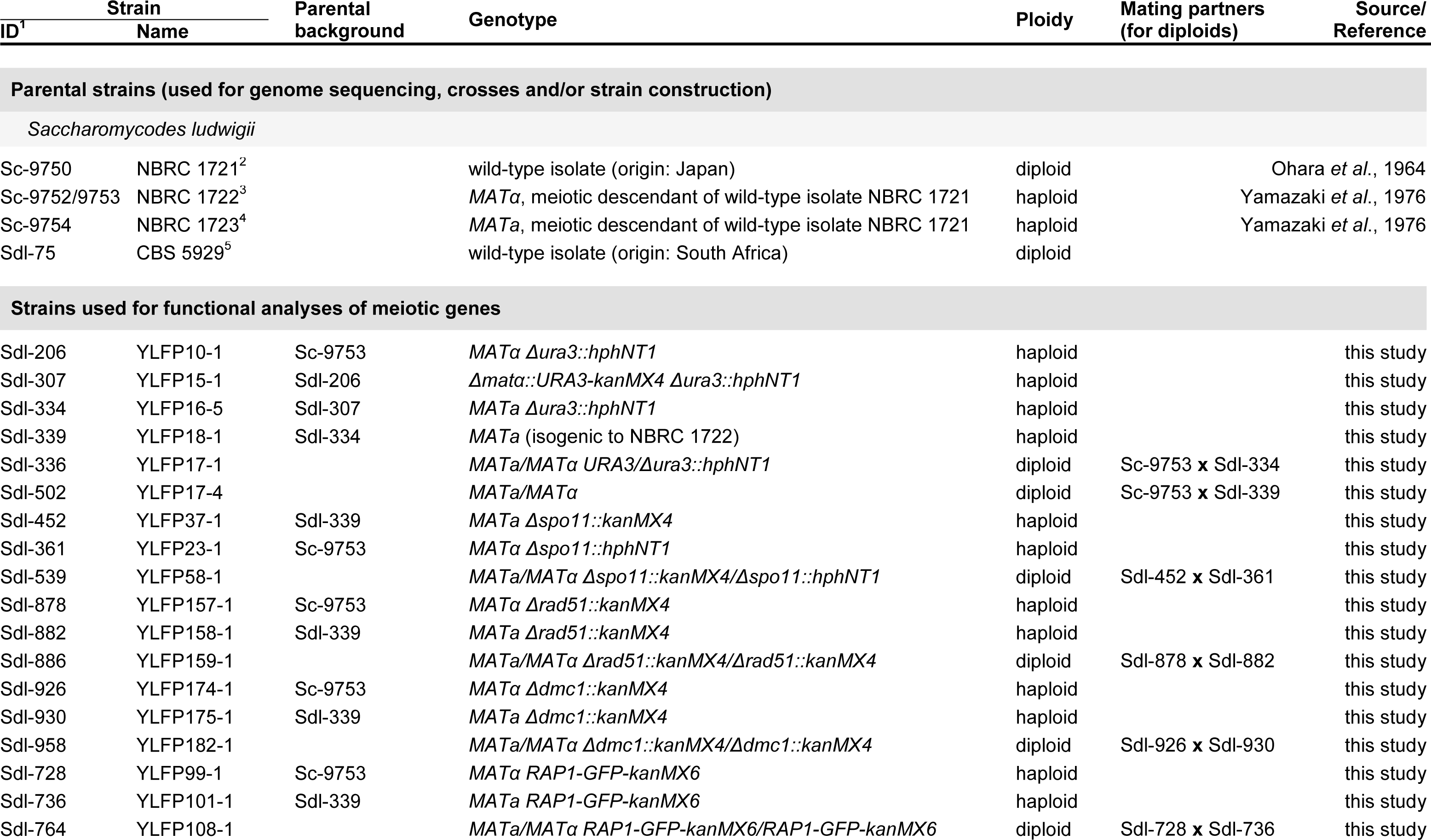
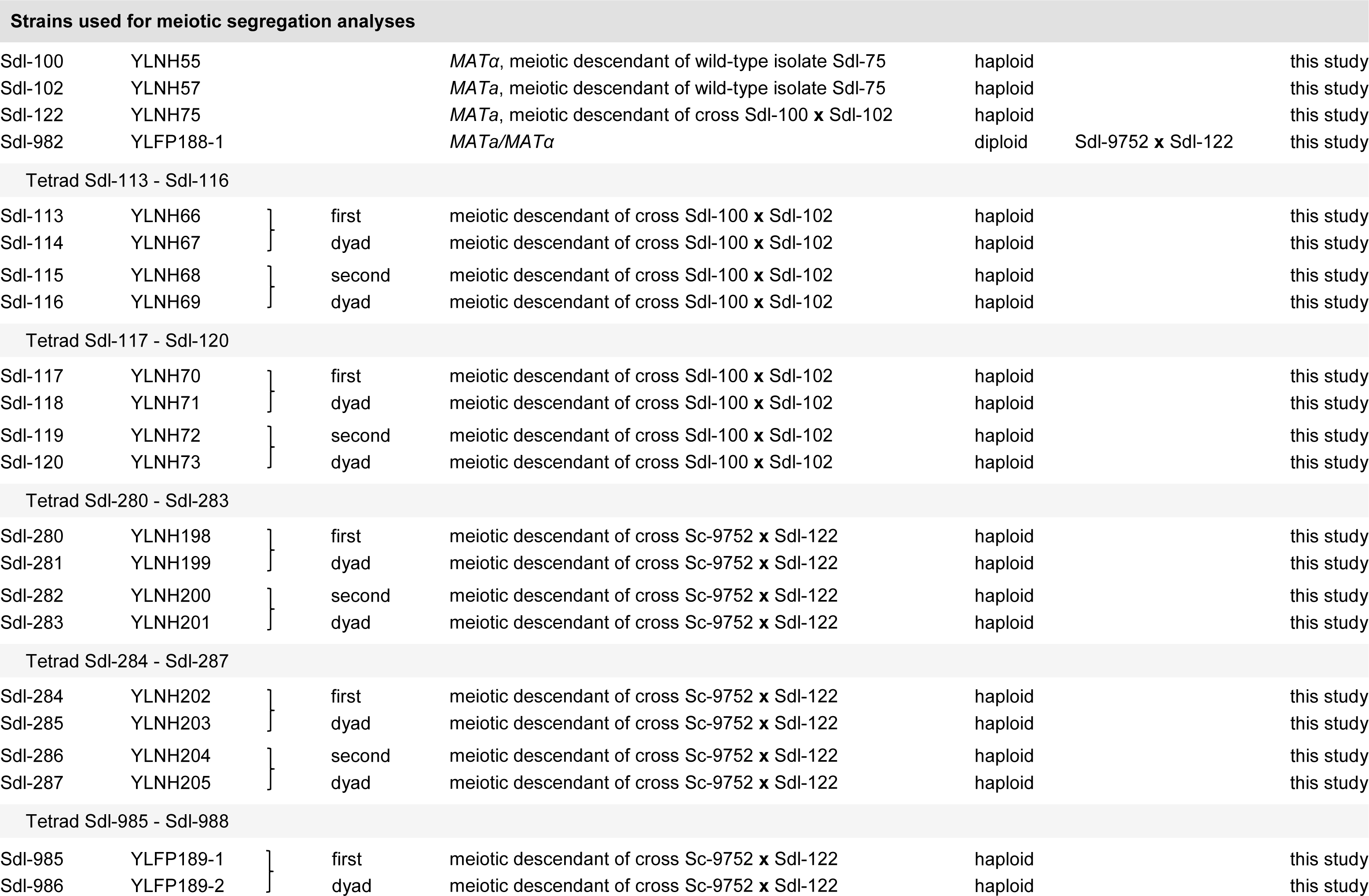
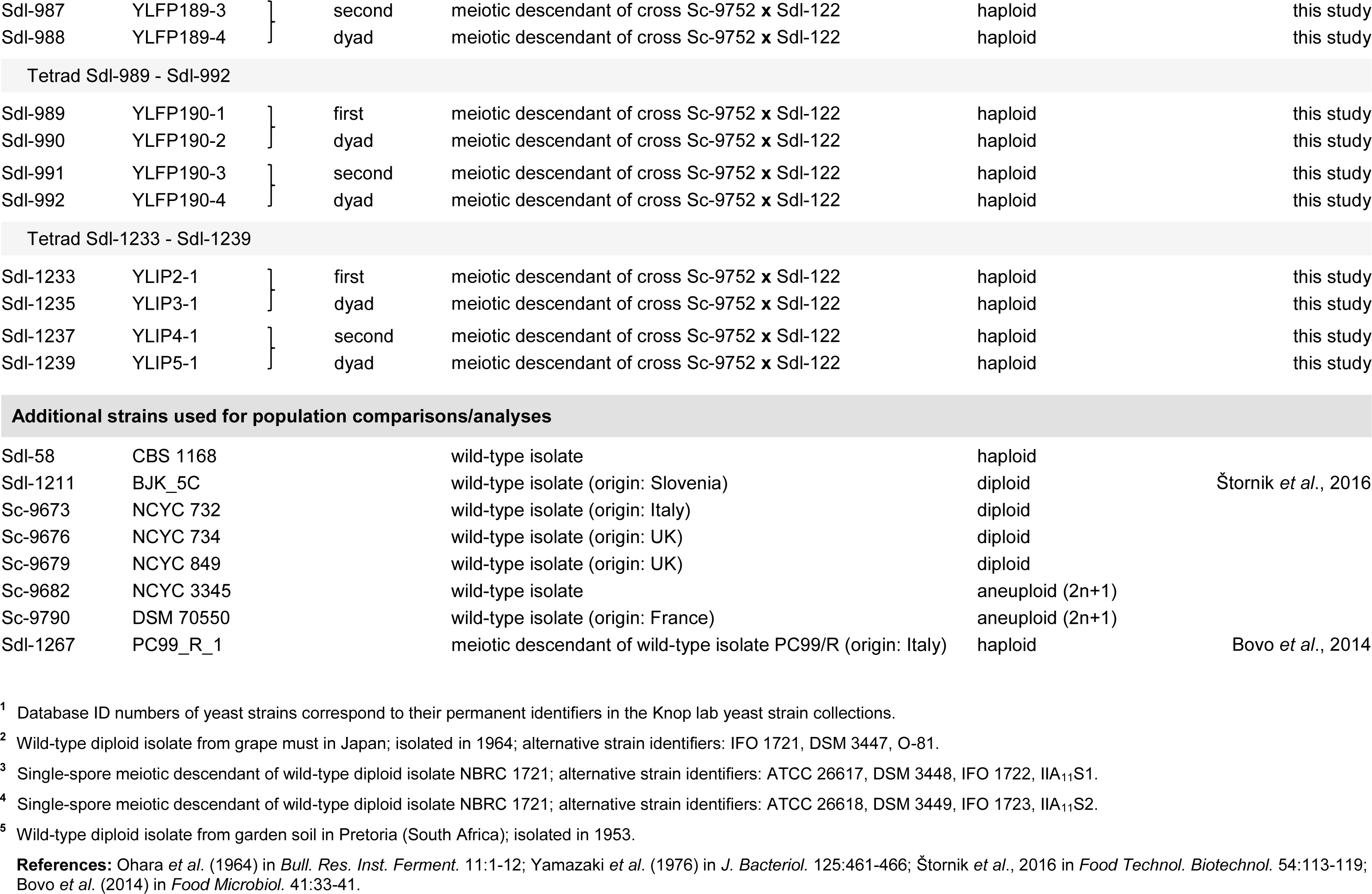
List of yeast strains generated and used in this study.

**Supplementary Table 2:**
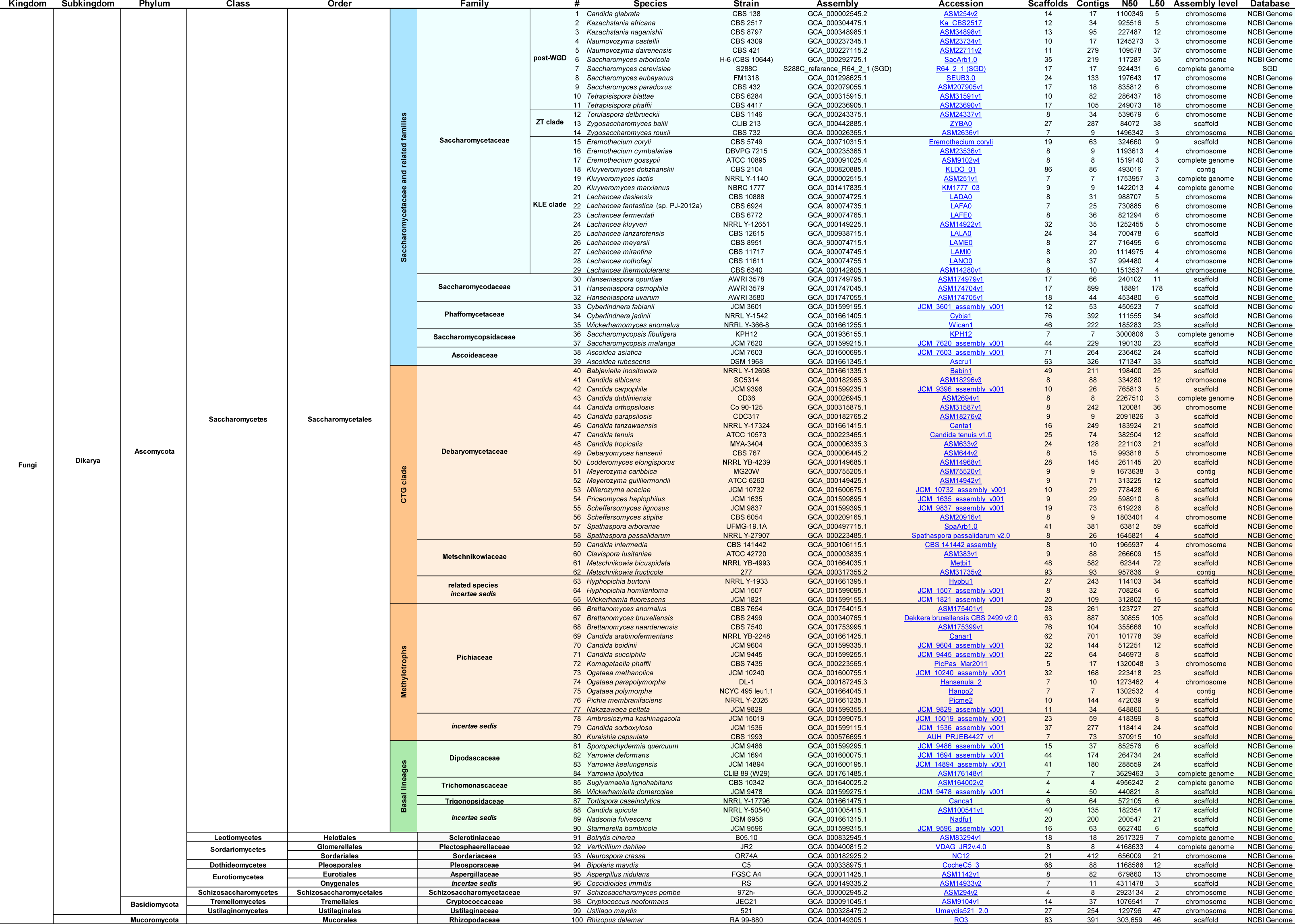
List of genome assemblies of yeasts and filamentous fungi analyzed in this study.

**Supplementary Table 3:**
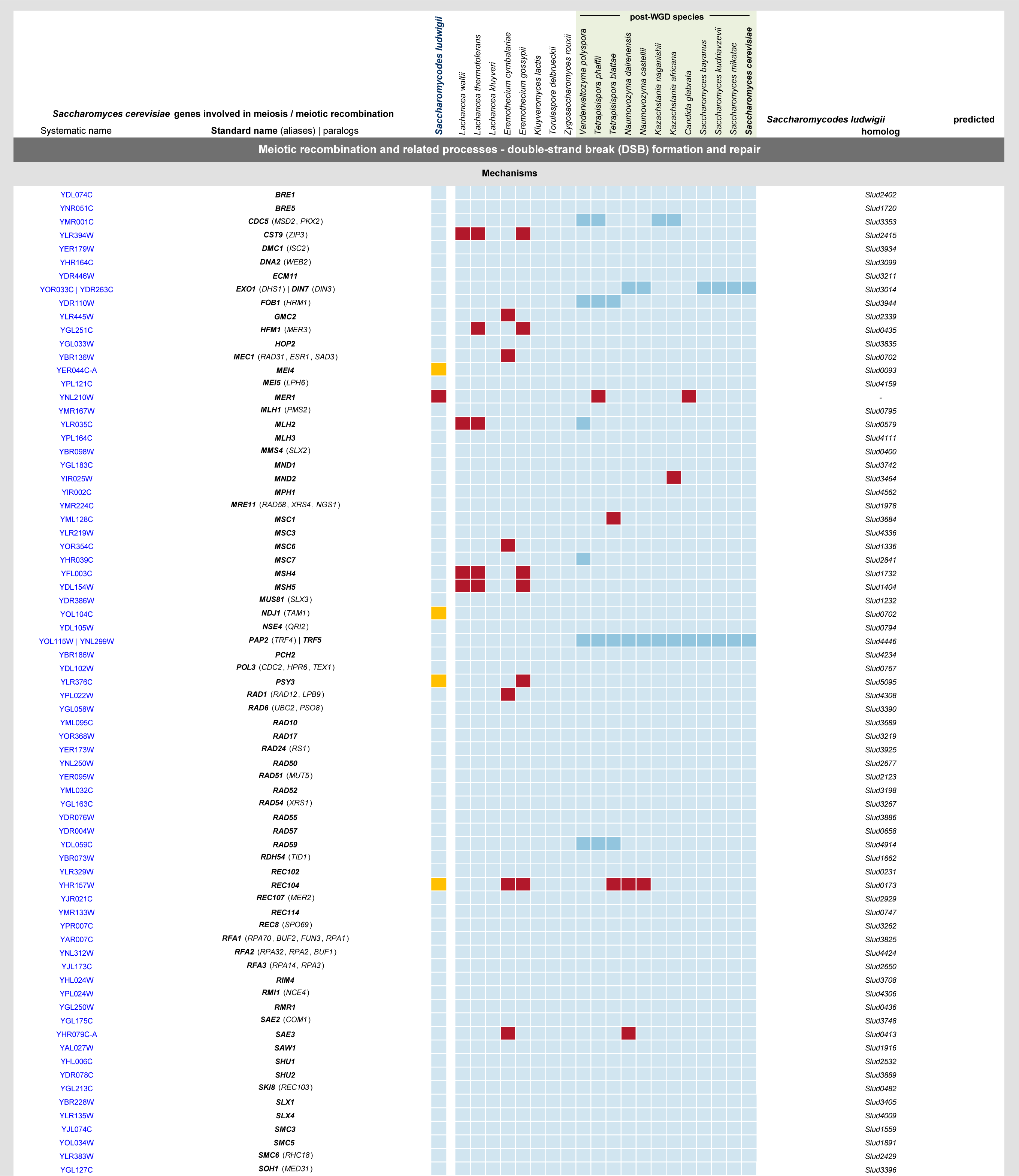
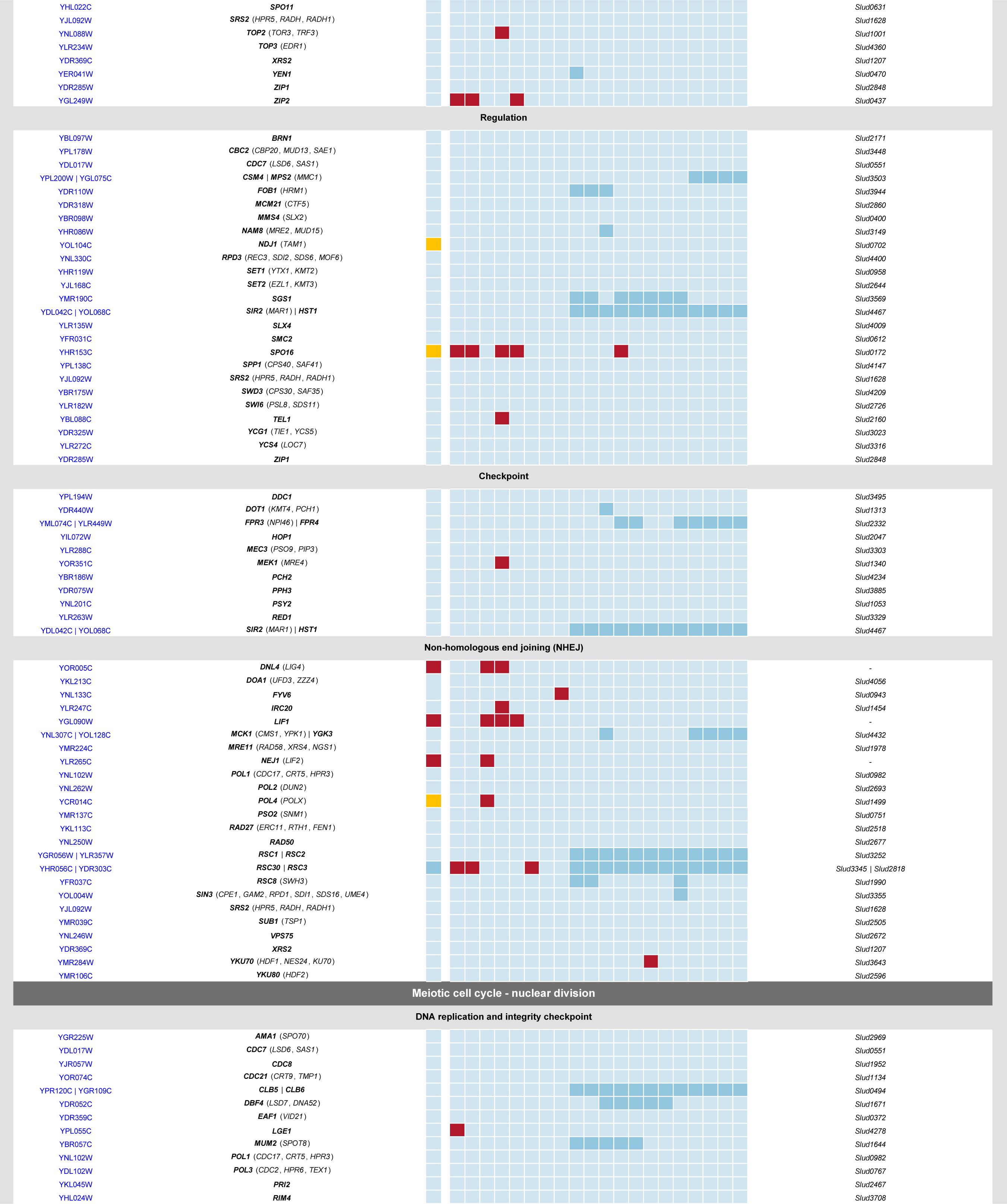
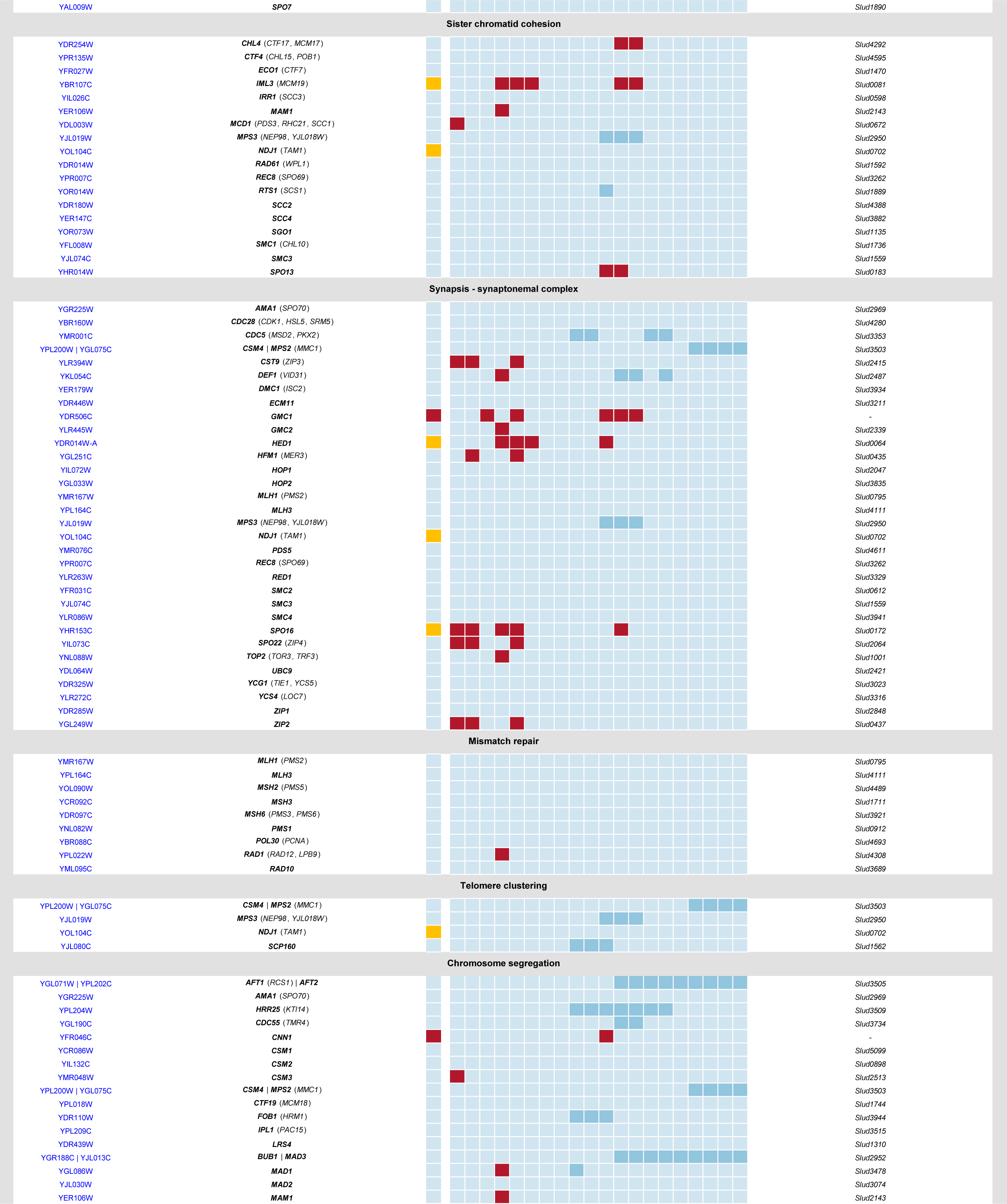
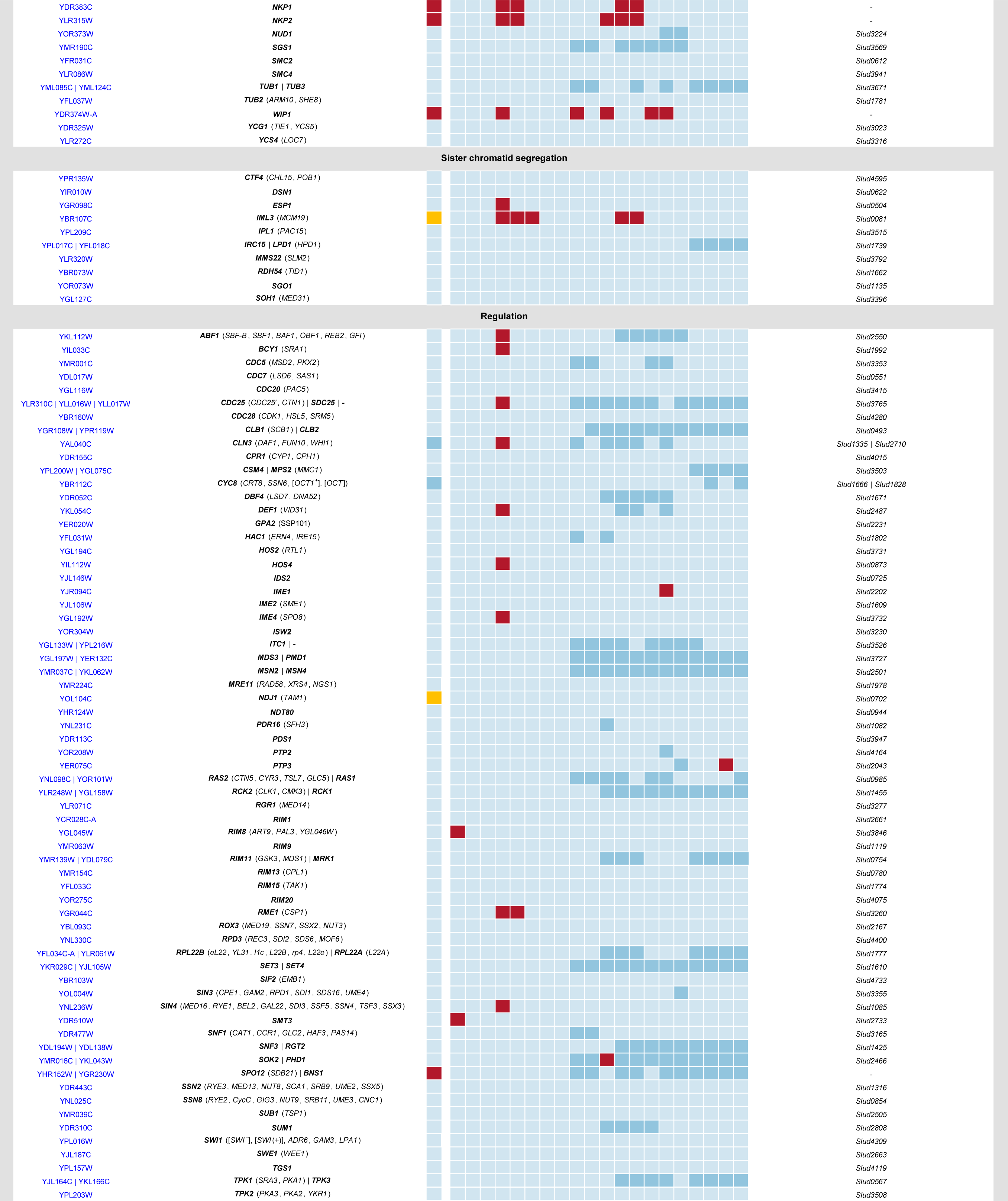
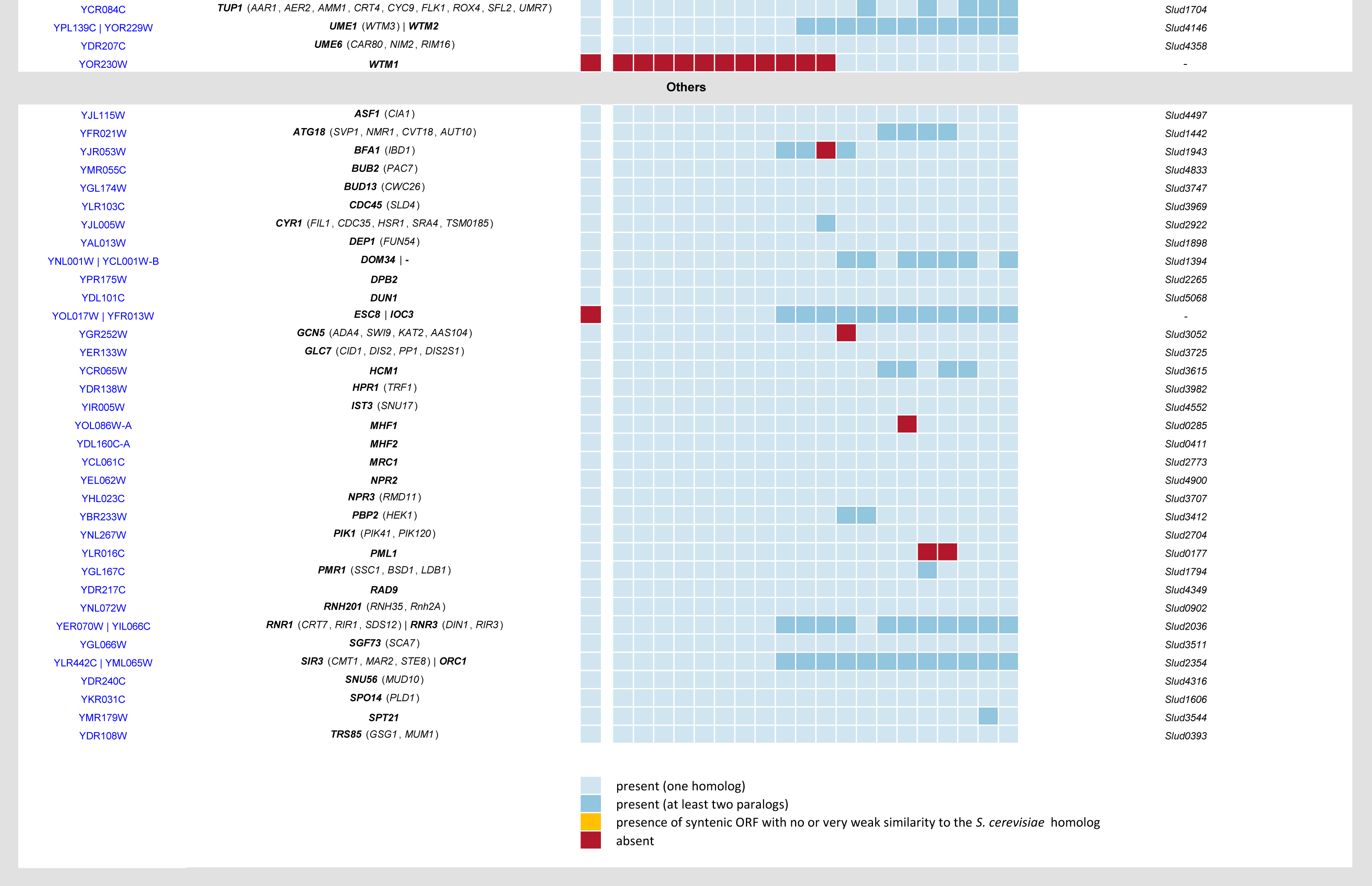
Genes involved in meiosis/meiotic recombination and their presence/absence in *Sd. ludwigii* and other yeasts.

**Supplementary Table 4:** Population analysis - read mapping statistics.

| Strain | Ploidy estimation<br>(flow cytometry) | Mapping on reference genome |  |  | Mapping on masked reference genome |  |  | Mapping on masked reference genome<br>(subtelomeres removed) |  |  |
| --- | --- | --- | --- | --- | --- | --- | --- | --- | --- | --- |
|  | Ploidy | # SNPs | # homozygous SNPs | # heterozygous SNPs | # SNPs | # homozygous SNPs | # heterozygous SNPs | # SNPs | # homozygous SNPs | # heterozygous SNPs |
| CBS 1168 | haploid | 41,533 | 35,052 | 6,481 | 32,832 | 26,991 | 5,841 | 27,466 | 25,455 | 2,011 |
| PC99_R_1 | haploid | 443,046 | 435,913 | 7,133 | 318,604 | 312,832 | 5,772 | 311,830 | 308,351 | 3,479 |
| 122 | haploid | 204,068 | 197,741 | 6,327 | 149,054 | 142,238 | 6,816 | 144,655 | 140,038 | 4,617 |
| UTAD17 | haploid<sup>1</sup> | 31,189 | 24,704 | 6,485 | 25,026 | 19,182 | 5,844 | 21,628 | 18,360 | 3,268 |
| NCYC 732 | diploid | 31,867 | 27,474 | 4,393 | 24,924 | 21,155 | 3,769 | 21,360 | 20,065 | 1,295 |
| NCYC 734 | diploid | 42,188 | 35,306 | 6,882 | 33,419 | 27,449 | 5,970 | 27,588 | 25,948 | 1,640 |
| NCYC 3345 | aneuploid (2n+1)<sup>2</sup> | 37,319 | 20,655 | 16,664 | 29,335 | 15,791 | 13,544 | 26,252 | 15,151 | 11,101 |
| BJK-5C | diploid | 46,611 | 25,013 | 21,598 | 37,367 | 19,283 | 18,084 | 33,375 | 18,268 | 15,107 |
| NCYC 849 | diploid | 40,172 | 17,423 | 22,749 | 31,809 | 13,309 | 18,500 | 28,065 | 12,879 | 15,186 |
| DSM 70550 | aneuploid (2n+1)<sup>3</sup> | 115,551 | 28,924 | 86,627 | 86,940 | 22,559 | 64,381 | 80,351 | 20,553 | 59,798 |

<sup>1</sup> The isolate was not available in our study, but it is presumed to be haploid (GenBank assembly accession: GCA\_900491785.1).
<sup>2</sup> Based on read coverage, the isolate was determined to have an additional (third) copy of chromosome D.
<sup>3</sup> Based on read coverage, the isolate was determined to have an additional (third) copy of chromosome G.

**Supplementary Table 5:**
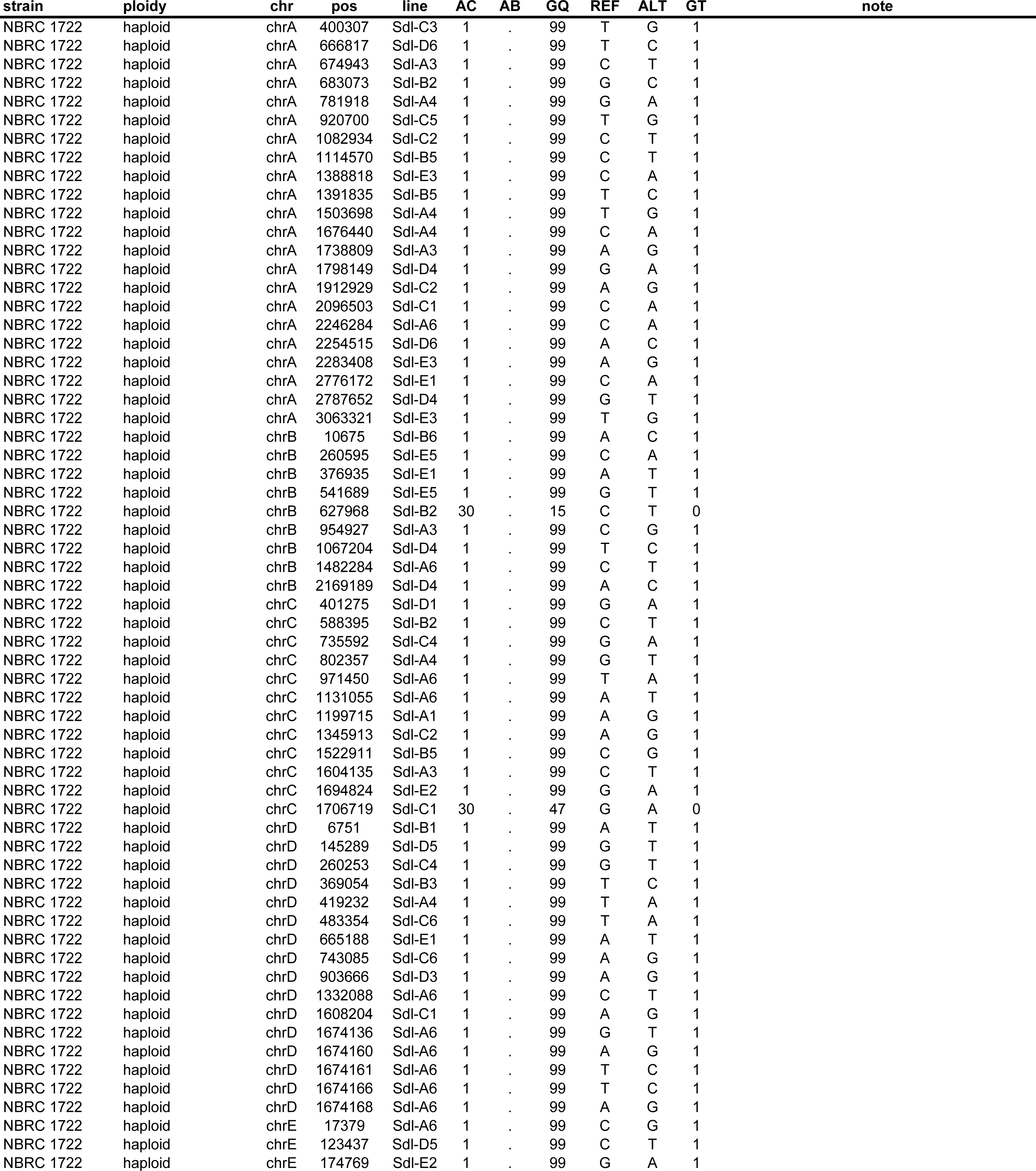
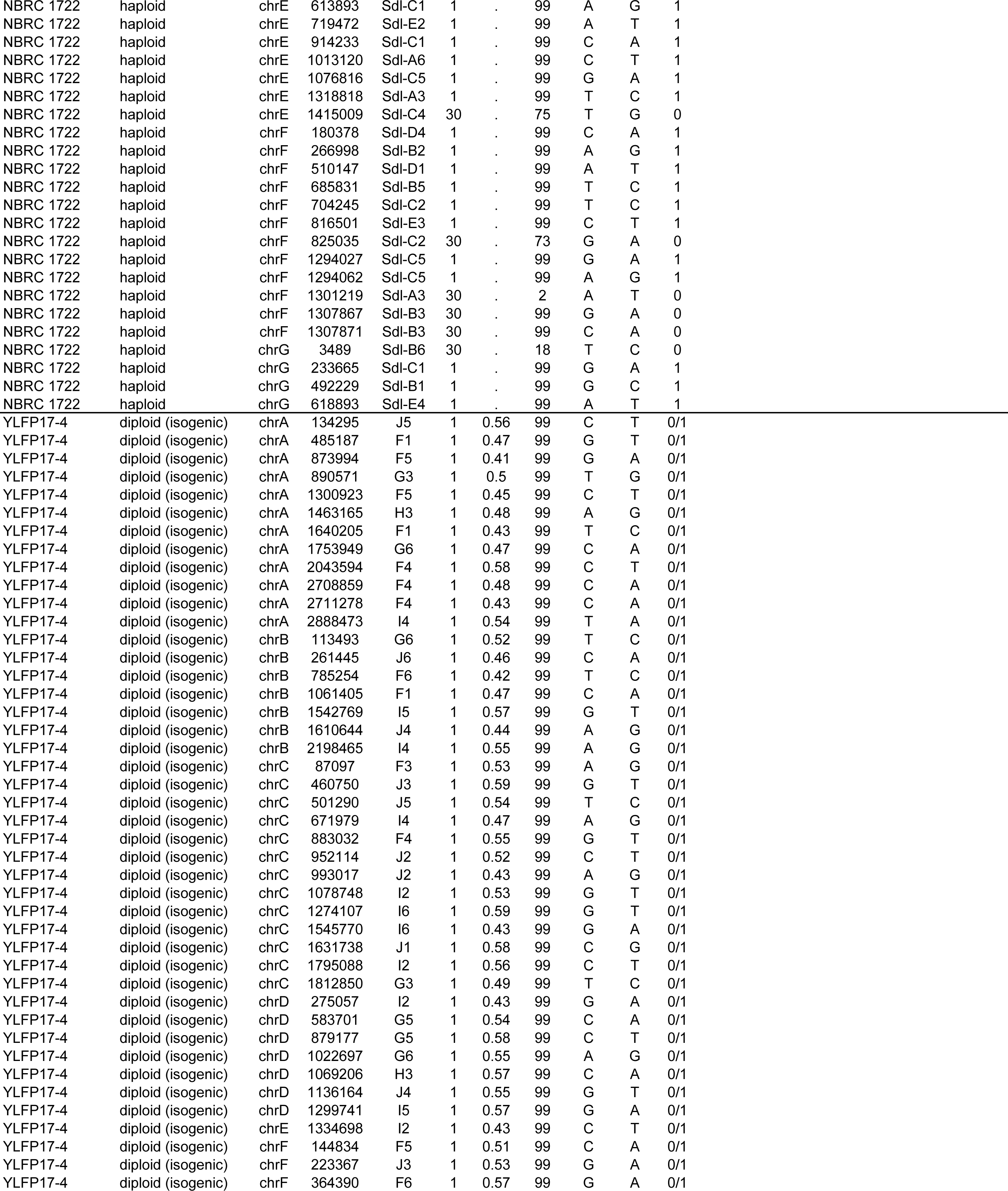
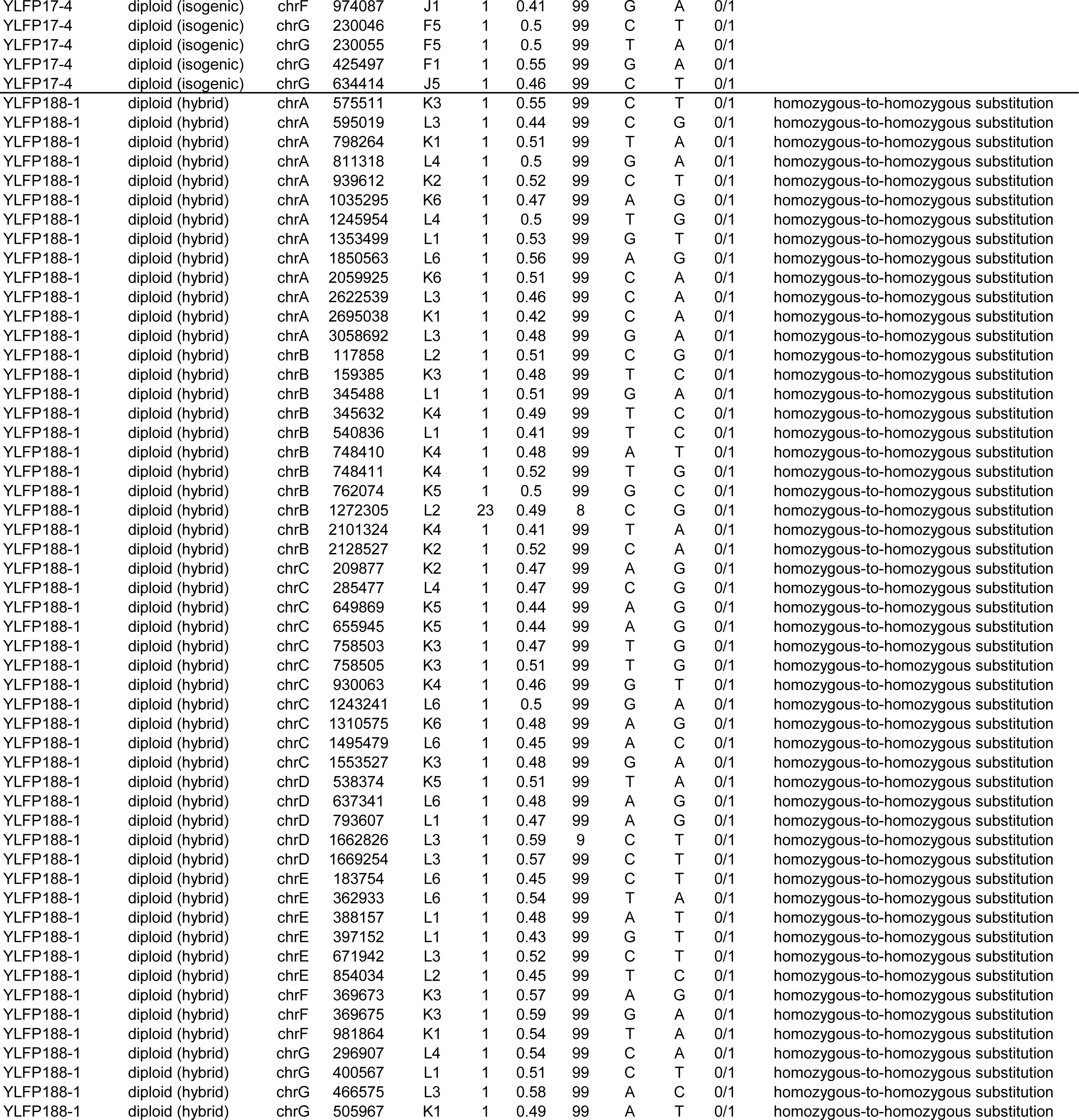
Novel mutations that arose during the mutation accumulation experiment.

| strain | ploidy | chr | pos | line | AC | AB | GQ | REF | ALT | GT | note |
| --- | --- | --- | --- | --- | --- | --- | --- | --- | --- | --- | --- |
| NBRC 1722 | haploid | chrA | 400307 | Sdl-C3 | 1 | . | 99 | T | G | 1 |  |
| NBRC 1722 | haploid | chrA | 666817 | Sdl-D6 | 1 | . | 99 | T | C | 1 |  |
| NBRC 1722 | haploid | chrA | 674943 | Sdl-A3 | 1 | . | 99 | C | T | 1 |  |
| NBRC 1722 | haploid | chrA | 683073 | Sdl-B2 | 1 | . | 99 | G | C | 1 |  |
| NBRC 1722 | haploid | chrA | 781918 | Sdl-A4 | 1 | . | 99 | G | A | 1 |  |
| NBRC 1722 | haploid | chrA | 920700 | Sdl-C5 | 1 | . | 99 | T | G | 1 |  |
| NBRC 1722 | haploid | chrA | 1082934 | Sdl-C2 | 1 | . | 99 | C | T | 1 |  |
| NBRC 1722 | haploid | chrA | 1114570 | Sdl-B5 | 1 | . | 99 | C | T | 1 |  |
| NBRC 1722 | haploid | chrA | 1388818 | Sdl-E3 | 1 | . | 99 | C | A | 1 |  |
| NBRC 1722 | haploid | chrA | 1391835 | Sdl-B5 | 1 | . | 99 | T | C | 1 |  |
| NBRC 1722 | haploid | chrA | 1503698 | Sdl-A4 | 1 | . | 99 | T | G | 1 |  |
| NBRC 1722 | haploid | chrA | 1676440 | Sdl-A4 | 1 | . | 99 | C | A | 1 |  |
| NBRC 1722 | haploid | chrA | 1738809 | Sdl-A3 | 1 | . | 99 | A | G | 1 |  |
| NBRC 1722 | haploid | chrA | 1798149 | Sdl-D4 | 1 | . | 99 | G | A | 1 |  |
| NBRC 1722 | haploid | chrA | 1912929 | Sdl-C2 | 1 | . | 99 | A | G | 1 |  |
| NBRC 1722 | haploid | chrA | 2096503 | Sdl-C1 | 1 | . | 99 | C | A | 1 |  |
| NBRC 1722 | haploid | chrA | 2246284 | Sdl-A6 | 1 | . | 99 | C | A | 1 |  |
| NBRC 1722 | haploid | chrA | 2254515 | Sdl-D6 | 1 | . | 99 | A | C | 1 |  |
| NBRC 1722 | haploid | chrA | 2283408 | Sdl-E3 | 1 | . | 99 | A | G | 1 |  |
| NBRC 1722 | haploid | chrA | 2776172 | Sdl-E1 | 1 | . | 99 | C | A | 1 |  |
| NBRC 1722 | haploid | chrA | 2787652 | Sdl-D4 | 1 | . | 99 | G | T | 1 |  |
| NBRC 1722 | haploid | chrA | 3063321 | Sdl-E3 | 1 | . | 99 | T | G | 1 |  |
| NBRC 1722 | haploid | chrB | 10675 | Sdl-B6 | 1 | . | 99 | A | C | 1 |  |
| NBRC 1722 | haploid | chrB | 260595 | Sdl-E5 | 1 | . | 99 | C | A | 1 |  |
| NBRC 1722 | haploid | chrB | 376935 | Sdl-E1 | 1 | . | 99 | A | T | 1 |  |
| NBRC 1722 | haploid | chrB | 541689 | Sdl-E5 | 1 | . | 99 | G | T | 1 |  |
| NBRC 1722 | haploid | chrB | 627968 | Sdl-B2 | 30 | . | 15 | C | T | 0 |  |
| NBRC 1722 | haploid | chrB | 954927 | Sdl-A3 | 1 | . | 99 | C | G | 1 |  |
| NBRC 1722 | haploid | chrB | 1067204 | Sdl-D4 | 1 | . | 99 | T | C | 1 |  |
| NBRC 1722 | haploid | chrB | 1482284 | Sdl-A6 | 1 | . | 99 | C | T | 1 |  |
| NBRC 1722 | haploid | chrB | 2169189 | Sdl-D4 | 1 | . | 99 | A | C | 1 |  |
| NBRC 1722 | haploid | chrC | 401275 | Sdl-D1 | 1 | . | 99 | G | A | 1 |  |
| NBRC 1722 | haploid | chrC | 588395 | Sdl-B2 | 1 | . | 99 | C | T | 1 |  |
| NBRC 1722 | haploid | chrC | 735592 | Sdl-C4 | 1 | . | 99 | G | A | 1 |  |
| NBRC 1722 | haploid | chrC | 802357 | Sdl-A4 | 1 | . | 99 | G | T | 1 |  |
| NBRC 1722 | haploid | chrC | 971450 | Sdl-A6 | 1 | . | 99 | T | A | 1 |  |
| NBRC 1722 | haploid | chrC | 1131055 | Sdl-A6 | 1 | . | 99 | A | T | 1 |  |
| NBRC 1722 | haploid | chrC | 1199715 | Sdl-A1 | 1 | . | 99 | A | G | 1 |  |
| NBRC 1722 | haploid | chrC | 1345913 | Sdl-C2 | 1 | . | 99 | A | G | 1 |  |
| NBRC 1722 | haploid | chrC | 1522911 | Sdl-B5 | 1 | . | 99 | C | G | 1 |  |
| NBRC 1722 | haploid | chrC | 1604135 | Sdl-A3 | 1 | . | 99 | C | T | 1 |  |
| NBRC 1722 | haploid | chrC | 1694824 | Sdl-E2 | 1 | . | 99 | G | A | 1 |  |
| NBRC 1722 | haploid | chrC | 1706719 | Sdl-C1 | 30 | . | 47 | G | A | 0 |  |
| NBRC 1722 | haploid | chrD | 6751 | Sdl-B1 | 1 | . | 99 | A | T | 1 |  |
| NBRC 1722 | haploid | chrD | 145289 | Sdl-D5 | 1 | . | 99 | G | T | 1 |  |
| NBRC 1722 | haploid | chrD | 260253 | Sdl-C4 | 1 | . | 99 | G | T | 1 |  |
| NBRC 1722 | haploid | chrD | 369054 | Sdl-B3 | 1 | . | 99 | T | C | 1 |  |
| NBRC 1722 | haploid | chrD | 419232 | Sdl-A4 | 1 | . | 99 | T | A | 1 |  |
| NBRC 1722 | haploid | chrD | 483354 | Sdl-C6 | 1 | . | 99 | T | A | 1 |  |
| NBRC 1722 | haploid | chrD | 665188 | Sdl-E1 | 1 | . | 99 | A | T | 1 |  |
| NBRC 1722 | haploid | chrD | 743085 | Sdl-C6 | 1 | . | 99 | A | G | 1 |  |
| NBRC 1722 | haploid | chrD | 903666 | Sdl-D3 | 1 | . | 99 | A | G | 1 |  |
| NBRC 1722 | haploid | chrD | 1332088 | Sdl-A6 | 1 | . | 99 | C | T | 1 |  |
| NBRC 1722 | haploid | chrD | 1608204 | Sdl-C1 | 1 | . | 99 | A | G | 1 |  |
| NBRC 1722 | haploid | chrD | 1674136 | Sdl-A6 | 1 | . | 99 | G | T | 1 |  |
| NBRC 1722 | haploid | chrD | 1674160 | Sdl-A6 | 1 | . | 99 | A | G | 1 |  |
| NBRC 1722 | haploid | chrD | 1674161 | Sdl-A6 | 1 | . | 99 | T | C | 1 |  |
| NBRC 1722 | haploid | chrD | 1674166 | Sdl-A6 | 1 | . | 99 | T | C | 1 |  |
| NBRC 1722 | haploid | chrD | 1674168 | Sdl-A6 | 1 | . | 99 | A | G | 1 |  |
| NBRC 1722 | haploid | chrE | 17379 | Sdl-A6 | 1 | . | 99 | C | G | 1 |  |
| NBRC 1722 | haploid | chrE | 123437 | Sdl-D5 | 1 | . | 99 | C | T | 1 |  |
| NBRC 1722 | haploid | chrE | 174769 | Sdl-E2 | 1 | . | 99 | G | A | 1 |  |

|  |  |  |  |  |  |  |  |  |  |  |
| --- | --- | --- | --- | --- | --- | --- | --- | --- | --- | --- |
| NBRC 1722 | haploid | chrE | 613893 | Sdl-C1 | 1 | . | 99 | A | G | 1 |
| NBRC 1722 | haploid | chrE | 719472 | Sdl-E2 | 1 | . | 99 | A | T | 1 |
| NBRC 1722 | haploid | chrE | 914233 | Sdl-C1 | 1 | . | 99 | C | A | 1 |
| NBRC 1722 | haploid | chrE | 1013120 | Sdl-A6 | 1 | . | 99 | C | T | 1 |
| NBRC 1722 | haploid | chrE | 1076816 | Sdl-C5 | 1 | . | 99 | G | A | 1 |
| NBRC 1722 | haploid | chrE | 1318818 | Sdl-A3 | 1 | . | 99 | T | C | 1 |
| NBRC 1722 | haploid | chrE | 1415009 | Sdl-C4 | 30 | . | 75 | T | G | 0 |
| NBRC 1722 | haploid | chrF | 180378 | Sdl-D4 | 1 | . | 99 | C | A | 1 |
| NBRC 1722 | haploid | chrF | 266998 | Sdl-B2 | 1 | . | 99 | A | G | 1 |
| NBRC 1722 | haploid | chrF | 510147 | Sdl-D1 | 1 | . | 99 | A | T | 1 |
| NBRC 1722 | haploid | chrF | 685831 | Sdl-B5 | 1 | . | 99 | T | C | 1 |
| NBRC 1722 | haploid | chrF | 704245 | Sdl-C2 | 1 | . | 99 | T | C | 1 |
| NBRC 1722 | haploid | chrF | 816501 | Sdl-E3 | 1 | . | 99 | C | T | 1 |
| NBRC 1722 | haploid | chrF | 825035 | Sdl-C2 | 30 | . | 73 | G | A | 0 |
| NBRC 1722 | haploid | chrF | 1294027 | Sdl-C5 | 1 | . | 99 | G | A | 1 |
| NBRC 1722 | haploid | chrF | 1294062 | Sdl-C5 | 1 | . | 99 | A | G | 1 |
| NBRC 1722 | haploid | chrF | 1301219 | Sdl-A3 | 30 | . | 2 | A | T | 0 |
| NBRC 1722 | haploid | chrF | 1307867 | Sdl-B3 | 30 | . | 99 | G | A | 0 |
| NBRC 1722 | haploid | chrF | 1307871 | Sdl-B3 | 30 | . | 99 | C | A | 0 |
| NBRC 1722 | haploid | chrG | 3489 | Sdl-B6 | 30 | . | 18 | T | C | 0 |
| NBRC 1722 | haploid | chrG | 233665 | Sdl-C1 | 1 | . | 99 | G | A | 1 |
| NBRC 1722 | haploid | chrG | 492229 | Sdl-B1 | 1 | . | 99 | G | C | 1 |
| NBRC 1722 | haploid | chrG | 618893 | Sdl-E4 | 1 | . | 99 | A | T | 1 |
| YLFP17-4 | diploid (isogenic) | chrA | 134295 | J5 | 1 | 0.56 | 99 | C | T | 0/1 |
| YLFP17-4 | diploid (isogenic) | chrA | 485187 | F1 | 1 | 0.47 | 99 | G | T | 0/1 |
| YLFP17-4 | diploid (isogenic) | chrA | 873994 | F5 | 1 | 0.41 | 99 | G | A | 0/1 |
| YLFP17-4 | diploid (isogenic) | chrA | 890571 | G3 | 1 | 0.5 | 99 | T | G | 0/1 |
| YLFP17-4 | diploid (isogenic) | chrA | 1300923 | F5 | 1 | 0.45 | 99 | C | T | 0/1 |
| YLFP17-4 | diploid (isogenic) | chrA | 1463165 | H3 | 1 | 0.48 | 99 | A | G | 0/1 |
| YLFP17-4 | diploid (isogenic) | chrA | 1640205 | F1 | 1 | 0.43 | 99 | T | C | 0/1 |
| YLFP17-4 | diploid (isogenic) | chrA | 1753949 | G6 | 1 | 0.47 | 99 | C | A | 0/1 |
| YLFP17-4 | diploid (isogenic) | chrA | 2043594 | F4 | 1 | 0.58 | 99 | C | T | 0/1 |
| YLFP17-4 | diploid (isogenic) | chrA | 2708859 | F4 | 1 | 0.48 | 99 | C | A | 0/1 |
| YLFP17-4 | diploid (isogenic) | chrA | 2711278 | F4 | 1 | 0.43 | 99 | C | A | 0/1 |
| YLFP17-4 | diploid (isogenic) | chrA | 2888473 | I4 | 1 | 0.54 | 99 | T | A | 0/1 |
| YLFP17-4 | diploid (isogenic) | chrB | 113493 | G6 | 1 | 0.52 | 99 | T | C | 0/1 |
| YLFP17-4 | diploid (isogenic) | chrB | 261445 | J6 | 1 | 0.46 | 99 | C | A | 0/1 |
| YLFP17-4 | diploid (isogenic) | chrB | 785254 | F6 | 1 | 0.42 | 99 | T | C | 0/1 |
| YLFP17-4 | diploid (isogenic) | chrB | 1061405 | F1 | 1 | 0.47 | 99 | C | A | 0/1 |
| YLFP17-4 | diploid (isogenic) | chrB | 1542769 | I5 | 1 | 0.57 | 99 | G | T | 0/1 |
| YLFP17-4 | diploid (isogenic) | chrB | 1610644 | J4 | 1 | 0.44 | 99 | A | G | 0/1 |
| YLFP17-4 | diploid (isogenic) | chrB | 2198465 | I4 | 1 | 0.55 | 99 | A | G | 0/1 |
| YLFP17-4 | diploid (isogenic) | chrC | 87097 | F3 | 1 | 0.53 | 99 | A | G | 0/1 |
| YLFP17-4 | diploid (isogenic) | chrC | 460750 | J3 | 1 | 0.59 | 99 | G | T | 0/1 |
| YLFP17-4 | diploid (isogenic) | chrC | 501290 | J5 | 1 | 0.54 | 99 | T | C | 0/1 |
| YLFP17-4 | diploid (isogenic) | chrC | 671979 | I4 | 1 | 0.47 | 99 | A | G | 0/1 |
| YLFP17-4 | diploid (isogenic) | chrC | 883032 | F4 | 1 | 0.55 | 99 | G | T | 0/1 |
| YLFP17-4 | diploid (isogenic) | chrC | 952114 | J2 | 1 | 0.52 | 99 | C | T | 0/1 |
| YLFP17-4 | diploid (isogenic) | chrC | 993017 | J2 | 1 | 0.43 | 99 | A | G | 0/1 |
| YLFP17-4 | diploid (isogenic) | chrC | 1078748 | I2 | 1 | 0.53 | 99 | G | T | 0/1 |
| YLFP17-4 | diploid (isogenic) | chrC | 1274107 | I6 | 1 | 0.59 | 99 | G | T | 0/1 |
| YLFP17-4 | diploid (isogenic) | chrC | 1545770 | I6 | 1 | 0.43 | 99 | G | A | 0/1 |
| YLFP17-4 | diploid (isogenic) | chrC | 1631738 | J1 | 1 | 0.58 | 99 | C | G | 0/1 |
| YLFP17-4 | diploid (isogenic) | chrC | 1795088 | I2 | 1 | 0.56 | 99 | C | T | 0/1 |
| YLFP17-4 | diploid (isogenic) | chrC | 1812850 | G3 | 1 | 0.49 | 99 | T | C | 0/1 |
| YLFP17-4 | diploid (isogenic) | chrD | 275057 | I2 | 1 | 0.43 | 99 | G | A | 0/1 |
| YLFP17-4 | diploid (isogenic) | chrD | 583701 | G5 | 1 | 0.54 | 99 | C | A | 0/1 |
| YLFP17-4 | diploid (isogenic) | chrD | 879177 | G5 | 1 | 0.58 | 99 | C | T | 0/1 |
| YLFP17-4 | diploid (isogenic) | chrD | 1022697 | G6 | 1 | 0.55 | 99 | A | G | 0/1 |
| YLFP17-4 | diploid (isogenic) | chrD | 1069206 | H3 | 1 | 0.57 | 99 | C | A | 0/1 |
| YLFP17-4 | diploid (isogenic) | chrD | 1136164 | J4 | 1 | 0.55 | 99 | G | T | 0/1 |
| YLFP17-4 | diploid (isogenic) | chrD | 1299741 | I5 | 1 | 0.57 | 99 | G | A | 0/1 |
| YLFP17-4 | diploid (isogenic) | chrE | 1334698 | I2 | 1 | 0.43 | 99 | C | T | 0/1 |
| YLFP17-4 | diploid (isogenic) | chrF | 144834 | F5 | 1 | 0.51 | 99 | C | A | 0/1 |
| YLFP17-4 | diploid (isogenic) | chrF | 223367 | J3 | 1 | 0.53 | 99 | G | A | 0/1 |
| YLFP17-4 | diploid (isogenic) | chrF | 364390 | F6 | 1 | 0.57 | 99 | G | A | 0/1 |
| YLFP17-4 | diploid (isogenic) | chrF | 974087 | J1 | 1 | 0.41 | 99 | G | A | 0/1 |  |
| YLFP17-4 | diploid (isogenic) | chrG | 230046 | F5 | 1 | 0.5 | 99 | C | T | 0/1 |  |
| YLFP17-4 | diploid (isogenic) | chrG | 230055 | F5 | 1 | 0.5 | 99 | T | A | 0/1 |  |
| YLFP17-4 | diploid (isogenic) | chrG | 425497 | F1 | 1 | 0.55 | 99 | G | A | 0/1 |  |
| YLFP17-4 | diploid (isogenic) | chrG | 634414 | J5 | 1 | 0.46 | 99 | C | T | 0/1 |  |
| YLFP188-1 | diploid (hybrid) | chrA | 575511 | K3 | 1 | 0.55 | 99 | C | T | 0/1 | homozygous-to-homozygous substitution |
| YLFP188-1 | diploid (hybrid) | chrA | 595019 | L3 | 1 | 0.44 | 99 | C | G | 0/1 | homozygous-to-homozygous substitution |
| YLFP188-1 | diploid (hybrid) | chrA | 798264 | K1 | 1 | 0.51 | 99 | T | A | 0/1 | homozygous-to-homozygous substitution |
| YLFP188-1 | diploid (hybrid) | chrA | 811318 | L4 | 1 | 0.5 | 99 | G | A | 0/1 | homozygous-to-homozygous substitution |
| YLFP188-1 | diploid (hybrid) | chrA | 939612 | K2 | 1 | 0.52 | 99 | C | T | 0/1 | homozygous-to-homozygous substitution |
| YLFP188-1 | diploid (hybrid) | chrA | 1035295 | K6 | 1 | 0.47 | 99 | A | G | 0/1 | homozygous-to-homozygous substitution |
| YLFP188-1 | diploid (hybrid) | chrA | 1245954 | L4 | 1 | 0.5 | 99 | T | G | 0/1 | homozygous-to-homozygous substitution |
| YLFP188-1 | diploid (hybrid) | chrA | 1353499 | L1 | 1 | 0.53 | 99 | G | T | 0/1 | homozygous-to-homozygous substitution |
| YLFP188-1 | diploid (hybrid) | chrA | 1850563 | L6 | 1 | 0.56 | 99 | A | G | 0/1 | homozygous-to-homozygous substitution |
| YLFP188-1 | diploid (hybrid) | chrA | 2059925 | K6 | 1 | 0.51 | 99 | C | A | 0/1 | homozygous-to-homozygous substitution |
| YLFP188-1 | diploid (hybrid) | chrA | 2622539 | L3 | 1 | 0.46 | 99 | C | A | 0/1 | homozygous-to-homozygous substitution |
| YLFP188-1 | diploid (hybrid) | chrA | 2695038 | K1 | 1 | 0.42 | 99 | C | A | 0/1 | homozygous-to-homozygous substitution |
| YLFP188-1 | diploid (hybrid) | chrA | 3058692 | L3 | 1 | 0.48 | 99 | G | A | 0/1 | homozygous-to-homozygous substitution |
| YLFP188-1 | diploid (hybrid) | chrB | 117858 | L2 | 1 | 0.51 | 99 | C | G | 0/1 | homozygous-to-homozygous substitution |
| YLFP188-1 | diploid (hybrid) | chrB | 159385 | K3 | 1 | 0.48 | 99 | T | C | 0/1 | homozygous-to-homozygous substitution |
| YLFP188-1 | diploid (hybrid) | chrB | 345488 | L1 | 1 | 0.51 | 99 | G | A | 0/1 | homozygous-to-homozygous substitution |
| YLFP188-1 | diploid (hybrid) | chrB | 345632 | K4 | 1 | 0.49 | 99 | T | C | 0/1 | homozygous-to-homozygous substitution |
| YLFP188-1 | diploid (hybrid) | chrB | 540836 | L1 | 1 | 0.41 | 99 | T | C | 0/1 | homozygous-to-homozygous substitution |
| YLFP188-1 | diploid (hybrid) | chrB | 748410 | K4 | 1 | 0.48 | 99 | A | T | 0/1 | homozygous-to-homozygous substitution |
| YLFP188-1 | diploid (hybrid) | chrB | 748411 | K4 | 1 | 0.52 | 99 | T | G | 0/1 | homozygous-to-homozygous substitution |
| YLFP188-1 | diploid (hybrid) | chrB | 762074 | K5 | 1 | 0.5 | 99 | G | C | 0/1 | homozygous-to-homozygous substitution |
| YLFP188-1 | diploid (hybrid) | chrB | 1272305 | L2 | 23 | 0.49 | 8 | C | G | 0/1 | homozygous-to-homozygous substitution |
| YLFP188-1 | diploid (hybrid) | chrB | 2101324 | K4 | 1 | 0.41 | 99 | T | A | 0/1 | homozygous-to-homozygous substitution |
| YLFP188-1 | diploid (hybrid) | chrB | 2128527 | K2 | 1 | 0.52 | 99 | C | A | 0/1 | homozygous-to-homozygous substitution |
| YLFP188-1 | diploid (hybrid) | chrC | 209877 | K2 | 1 | 0.47 | 99 | A | G | 0/1 | homozygous-to-homozygous substitution |
| YLFP188-1 | diploid (hybrid) | chrC | 285477 | L4 | 1 | 0.47 | 99 | C | G | 0/1 | homozygous-to-homozygous substitution |
| YLFP188-1 | diploid (hybrid) | chrC | 649869 | K5 | 1 | 0.44 | 99 | A | G | 0/1 | homozygous-to-homozygous substitution |
| YLFP188-1 | diploid (hybrid) | chrC | 655945 | K5 | 1 | 0.44 | 99 | A | G | 0/1 | homozygous-to-homozygous substitution |
| YLFP188-1 | diploid (hybrid) | chrC | 758503 | K3 | 1 | 0.47 | 99 | T | G | 0/1 | homozygous-to-homozygous substitution |
| YLFP188-1 | diploid (hybrid) | chrC | 758505 | K3 | 1 | 0.51 | 99 | T | G | 0/1 | homozygous-to-homozygous substitution |
| YLFP188-1 | diploid (hybrid) | chrC | 930063 | K4 | 1 | 0.46 | 99 | G | T | 0/1 | homozygous-to-homozygous substitution |
| YLFP188-1 | diploid (hybrid) | chrC | 1243241 | L6 | 1 | 0.5 | 99 | G | A | 0/1 | homozygous-to-homozygous substitution |
| YLFP188-1 | diploid (hybrid) | chrC | 1310575 | K6 | 1 | 0.48 | 99 | A | G | 0/1 | homozygous-to-homozygous substitution |
| YLFP188-1 | diploid (hybrid) | chrC | 1495479 | L6 | 1 | 0.45 | 99 | A | C | 0/1 | homozygous-to-homozygous substitution |
| YLFP188-1 | diploid (hybrid) | chrC | 1553527 | K3 | 1 | 0.48 | 99 | G | A | 0/1 | homozygous-to-homozygous substitution |
| YLFP188-1 | diploid (hybrid) | chrD | 538374 | K5 | 1 | 0.51 | 99 | T | A | 0/1 | homozygous-to-homozygous substitution |
| YLFP188-1 | diploid (hybrid) | chrD | 637341 | L6 | 1 | 0.48 | 99 | A | G | 0/1 | homozygous-to-homozygous substitution |
| YLFP188-1 | diploid (hybrid) | chrD | 793607 | L1 | 1 | 0.47 | 99 | A | G | 0/1 | homozygous-to-homozygous substitution |
| YLFP188-1 | diploid (hybrid) | chrD | 1662826 | L3 | 1 | 0.59 | 9 | C | T | 0/1 | homozygous-to-homozygous substitution |
| YLFP188-1 | diploid (hybrid) | chrD | 1669254 | L3 | 1 | 0.57 | 99 | C | T | 0/1 | homozygous-to-homozygous substitution |
| YLFP188-1 | diploid (hybrid) | chrE | 183754 | L6 | 1 | 0.45 | 99 | C | T | 0/1 | homozygous-to-homozygous substitution |
| YLFP188-1 | diploid (hybrid) | chrE | 362933 | L6 | 1 | 0.54 | 99 | T | A | 0/1 | homozygous-to-homozygous substitution |
| YLFP188-1 | diploid (hybrid) | chrE | 388157 | L1 | 1 | 0.48 | 99 | A | T | 0/1 | homozygous-to-homozygous substitution |
| YLFP188-1 | diploid (hybrid) | chrE | 397152 | L1 | 1 | 0.43 | 99 | G | T | 0/1 | homozygous-to-homozygous substitution |
| YLFP188-1 | diploid (hybrid) | chrE | 671942 | L3 | 1 | 0.52 | 99 | C | T | 0/1 | homozygous-to-homozygous substitution |
| YLFP188-1 | diploid (hybrid) | chrE | 854034 | L2 | 1 | 0.45 | 99 | T | C | 0/1 | homozygous-to-homozygous substitution |
| YLFP188-1 | diploid (hybrid) | chrF | 369673 | K3 | 1 | 0.57 | 99 | A | G | 0/1 | homozygous-to-homozygous substitution |
| YLFP188-1 | diploid (hybrid) | chrF | 369675 | K3 | 1 | 0.59 | 99 | G | A | 0/1 | homozygous-to-homozygous substitution |
| YLFP188-1 | diploid (hybrid) | chrF | 981864 | K1 | 1 | 0.54 | 99 | T | A | 0/1 | homozygous-to-homozygous substitution |
| YLFP188-1 | diploid (hybrid) | chrG | 296907 | L4 | 1 | 0.54 | 99 | C | A | 0/1 | homozygous-to-homozygous substitution |
| YLFP188-1 | diploid (hybrid) | chrG | 400567 | L1 | 1 | 0.51 | 99 | C | T | 0/1 | homozygous-to-homozygous substitution |
| YLFP188-1 | diploid (hybrid) | chrG | 466575 | L3 | 1 | 0.58 | 99 | A | C | 0/1 | homozygous-to-homozygous substitution |
| YLFP188-1 | diploid (hybrid) | chrG | 505967 | K1 | 1 | 0.49 | 99 | A | T | 0/1 | homozygous-to-homozygous substitution |

**Supplementary Table 6:** List of LOH tracts detected in the mutation accumulation analysis.

| Mutation accumulation line<br>(hybrid diploid ancestry) | chromosome | beginning | Loss-of-heterozygosity (LOH) tract |  |  | number of SNP markers |
| --- | --- | --- | --- | --- | --- | --- |
|  |  |  | end | minimum size (bp) |  |  |
| K1 | chrA | 411449 | 413791 | 2343 |  | 6 |
| L1 | chrA | 513059 | 522453 | 9395 |  | 20 |
| L3 | chrA | 578207 | 578207 | 1 |  | 1 |
| K1 | chrA | 948826 | 948826 | 1 |  | 1 |
| K6 | chrA | 1222049 | 1223493 | 1445 |  | 5 |
| K5 | chrA | 1505212 | 1506895 | 1684 |  | 9 |
| K3 | chrA | 1661620 | 1661620 | 1 |  | 1 |
| K5 | chrA | 1708548 | 1708548 | 1 |  | 1 |
| L2 | chrA | 2188868 | 2189916 | 1049 |  | 11 |
| L4 | chrA | 3017907 | 3017907 | 1 |  | 1 |
| K1 | chrA | 3066698 | 3066698 | 1 |  | 1 |
| L6 | chrB | 514822 | 514849 | 28 |  | 2 |
| K1 | chrC | 1258 | 182642 | 181385 |  | 4190 |
| L4 | chrC | 281216 | 281366 | 151 |  | 6 |
| L2 | chrC | 336505 | 336505 | 1 |  | 1 |
| L2 | chrD | 344988 | 357464 | 12477 |  | 23 |
| L3 | chrD | 1666276 | 1675901 | 9626 |  | 7 |
| L3 | chrE | 830403 | 830403 | 1 |  | 1 |
| K3 | chrE | 944151 | 944151 | 1 |  | 1 |
| L3 | chrE | 956070 | 956070 | 1 |  | 1 |
| L6 | chrG | 21383 | 21691 | 309 |  | 2 |
| K3 | chrG | 281226 | 281272 | 47 |  | 4 |

**Supplementary Table 7:**
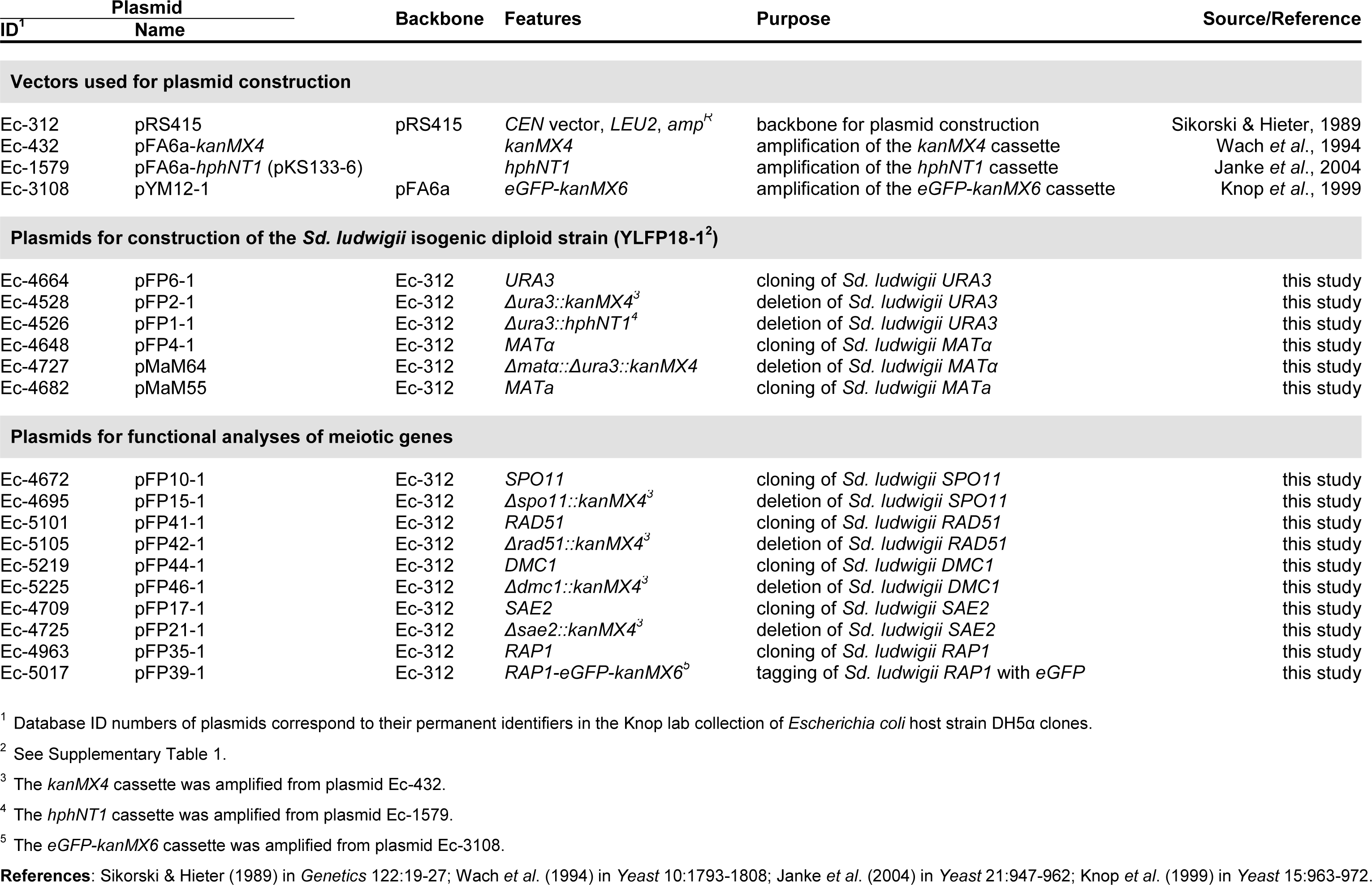
List of plasmids generated and used in this study.

**Supplementary Table 8:**
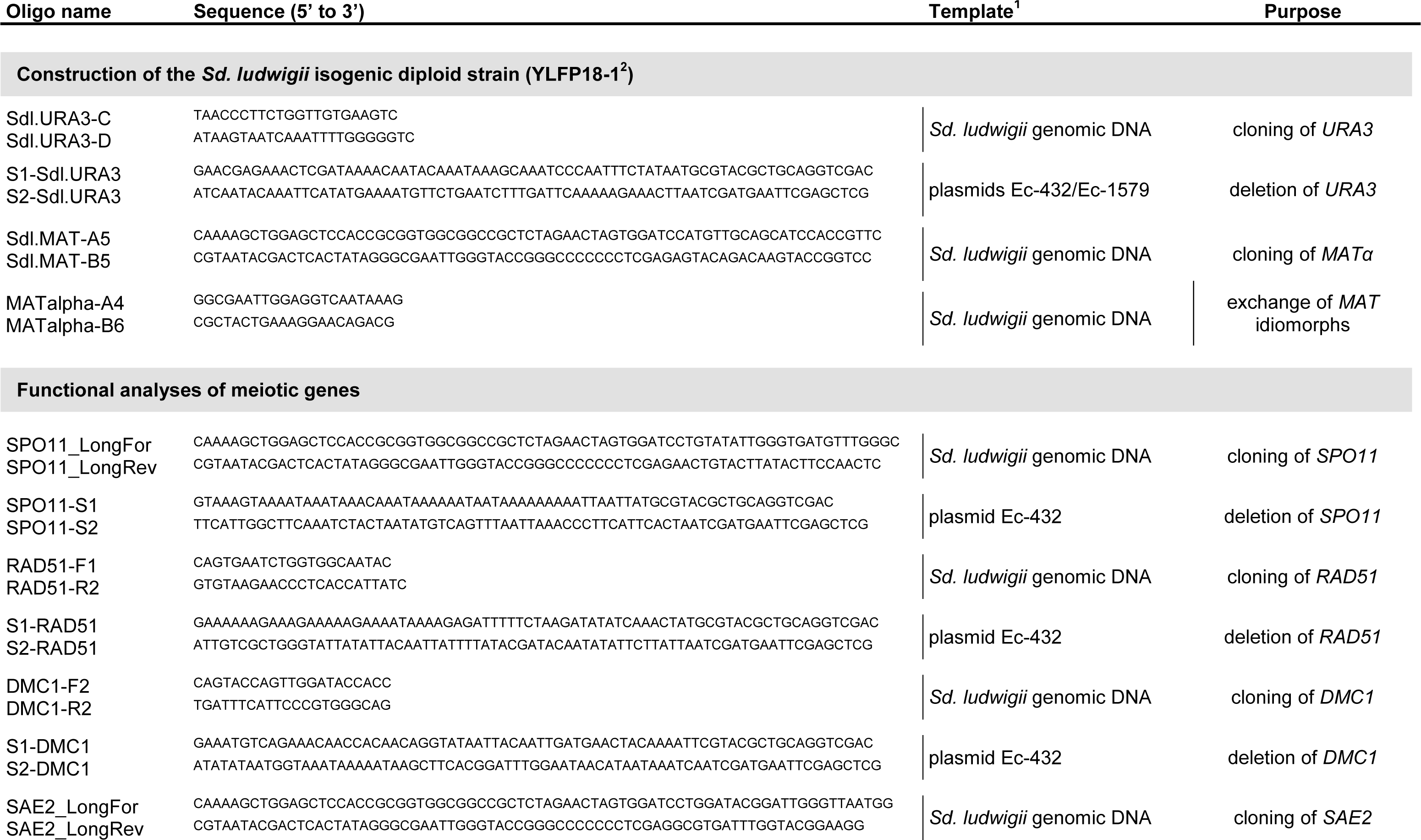
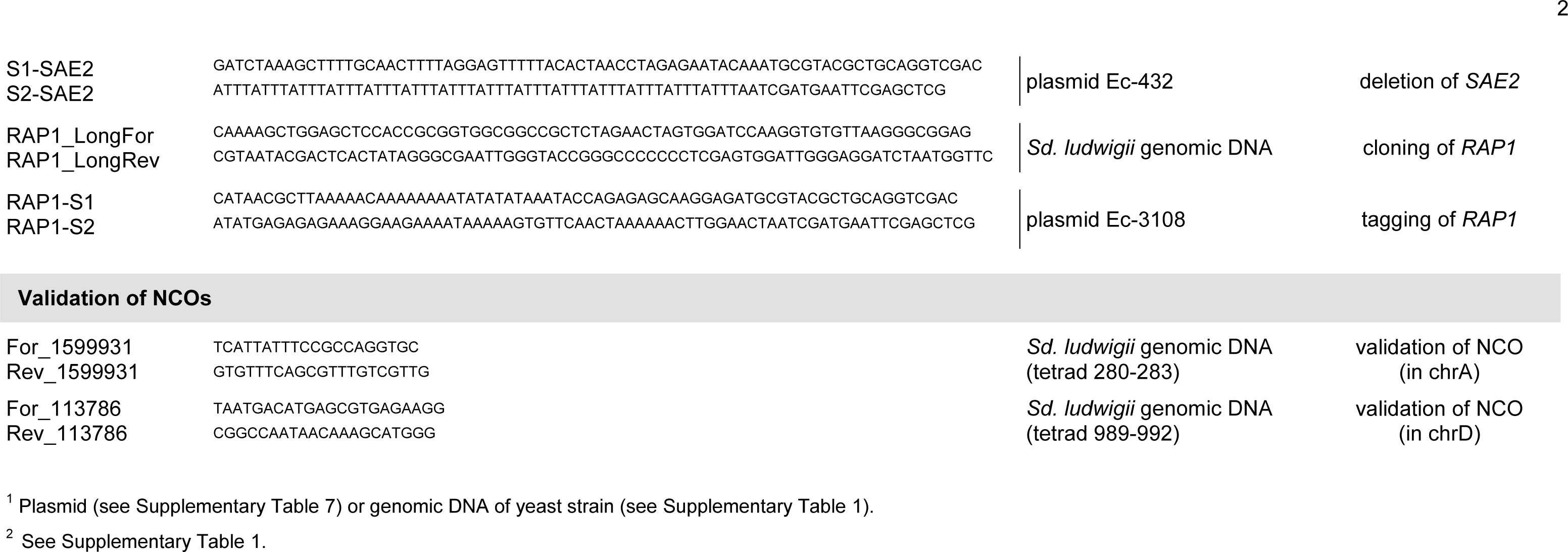
List of DNA oligonucleotides used in this study.

## Notes

### Competing Interest Statement

The authors have declared no competing interest.

